# Genetic diversity and evolution of rice centromeres

**DOI:** 10.1101/2024.07.28.605524

**Authors:** Lingjuan Xie, Yujie Huang, Wei Huang, Lianguang Shang, Yanqing Sun, Quanyu Chen, Shuangtian Bi, Mingyu Suo, Shiyu Zhang, Chentao Yang, Xiaoming Zheng, Weiwei Jin, Qian Qian, Longjiang Fan, Dongya Wu

## Abstract

Understanding the mechanisms driving centromere evolution is crucial for deciphering eukaryotic evolution and speciation processes. Despite their widely recognized characteristics of conserved function in cell division, the centromeres have showed high diversity in composition and structure between species. The mechanism underlying this paradox remain poorly understood. Here, we assembled 67 high-quality rice genomes from *Oryza* AA group, encompassing both Asian and African rice species, and conducted an extensive analysis of over 800 nearly complete centromeres. Through *de novo* annotation of satellite sequences and employing a progressive compression strategy, we quantified the local homogenization and multi-layer nested structures of rice centromeres and found that genetic innovations in rice centromeres primarily arise from internal structural variations and retrotransposon insertions, along with a certain number of non-canonical satellite repeats (*sati*). Despite these rapid structural alterations, the single-base substitution rate in rice centromeres appears relatively lower compared to the chromosome arms. Contrary to the KARMA model for *Arabidopsis* centromere evolution, our model (RICE) suggests that centrophilic LTRs contribute to the decline of progenitor centromeres composed of satellite repeats, and facilitate the formation of evolutionary neo-centromeres, which are enriched with extended CENH3 binding regions beyond the native satellite arrays in plant genomes. In summary, this study provides novel insights into genomic divergence and reproductive barriers among rice species and subspecies, and advances our understanding of plant centromere evolution.

## Introduction

Centromeres are the chromosomal regions that are physically linked to the spindle microtubules during mitosis and meiosis. They are ubiquitous and essential in eukaryotic life for maintaining their genetic stability (Barra & Fachinetti, 2018; Naish & Henderson, 2024). Most centromeres are mainly composed of highly repetitive tandem satellite DNA sequences as the sites of kinetochore assembly and interspersed transposon elements (TEs), but their monomer sequences, satellite copy numbers and array organizations are diversely variable (Naish *et al*., 2021; Ahmed *et al*., 2023; Naish & Henderson, 2024). While canonical high-order repeat (HOR) structures are prevalent in human centromeres, such arrangements are less common in plant centromeres (Naish *et al*., 2021; Altemose *et al*., 2022; Wang *et al*, 2023; Logsdon *et al*., 2024). Conversely, plant centromeres frequently harbour centrophilic transposon insertions, a feature that distinguishes them from centromeres in the human and chicken (Cheng *et al*., 2002; Naish *et al*., 2021; Altemose *et al*., 2022; Chen *et al*., 2023; Huang *et al*., 2023; Lv *et al*., 2023). The functional determination of centromeres often relies on the presence of the centromere-specific histone variant CENH3 or CENP-A, which marks the sites of kinetochore assembly (Malik & Henikoff, 2009; Kursel & Malik, 2016). The genomic regions of centromeres defined by satellite DNA occupancy and CENH3 Chromatin Immunoprecipitation Sequencing (ChIP-seq) enrichment can sometimes show discrepancies, suggesting instances of centromere repositioning and the emergence of evolutionary-new centromeres (ENCs) (Liu *et al*., 2023). However, the mechanisms driving the occurrence of ENCs remain incompletely understood. Despite recent advancements in generating complete centromere assemblies using PacBio high-fidelity (HiFi) sequencing and Oxford Nanopore Technology (ONT) ultra-long sequencing, our understanding to the rapid centromere evolution is still limited, owing to the challenges in comparing highly repetitive sequences among species and the limited sampling within species (Xie *et al*., 2024). So far, comprehensive analyzes on population-scale centromeres with high accuracy and continuity have been only performed in the genomes of human and model species *Arabidopsis*, which display the undiscovered genomic diversity and provide valuable resources for centromere genomics (Wlodzimierz *et al*., 2023; Logsdon *et al*., 2024). Extending such analyses to diverse species will be crucial for capturing the trajectory of centromere evolution and understanding its implications for genome stability and speciation.

Rice is one of the major staple crops globally, serving as a primary energy source for human. It is also one of the major model plant species. Recurrent domestication, improvement and de-domestication events have been parallelly experienced for different species or subspecies in the rice AA genome group (Stein *et al*., 2018). Asian rice (*Oryza sativa*) and African rice (*O. glaberrima*) are the two main crops in *Oryza* AA group with a divergence time less than 1 million years, domesticated from their wild ancestor *Oryza rufipogon* and *Oryza barthii*, respectively (**Fig. 1a**). Three telomere-to-telomere (T2T) genome assemblies of Asian rice (varieties Zhenshan97, Minghui63 and Nipponbare) have been completely assembled (Song *et al*., 2021; Shang *et al*., 2023). Notwithstanding the achievements in generating T2T assemblies, the precise alignment of highly repetitive sequences within centromeres remains a significant challenge. Consequently, systematic comparisons of centromeres across different rice species have been limited. Here we assemble and annotate the centromeres from 70 nearly T2T rice genomes of *Oryza* AA group. We introduce a novel framework for centromere analysis and perform comparative investigations of rice centromeres, probing into the satellite sequence composition, satellite array organization, the impact of transposon invasion and the dynamics of epigenetic modifications among species, subspecies, groups, individuals and chromosomes. Through elucidating the intricate mechanisms governing rice centromere evolution, our study aims to offer new insights into the genetic and epigenetic factors that shape these crucial genomic regions.

**Figure 1.**
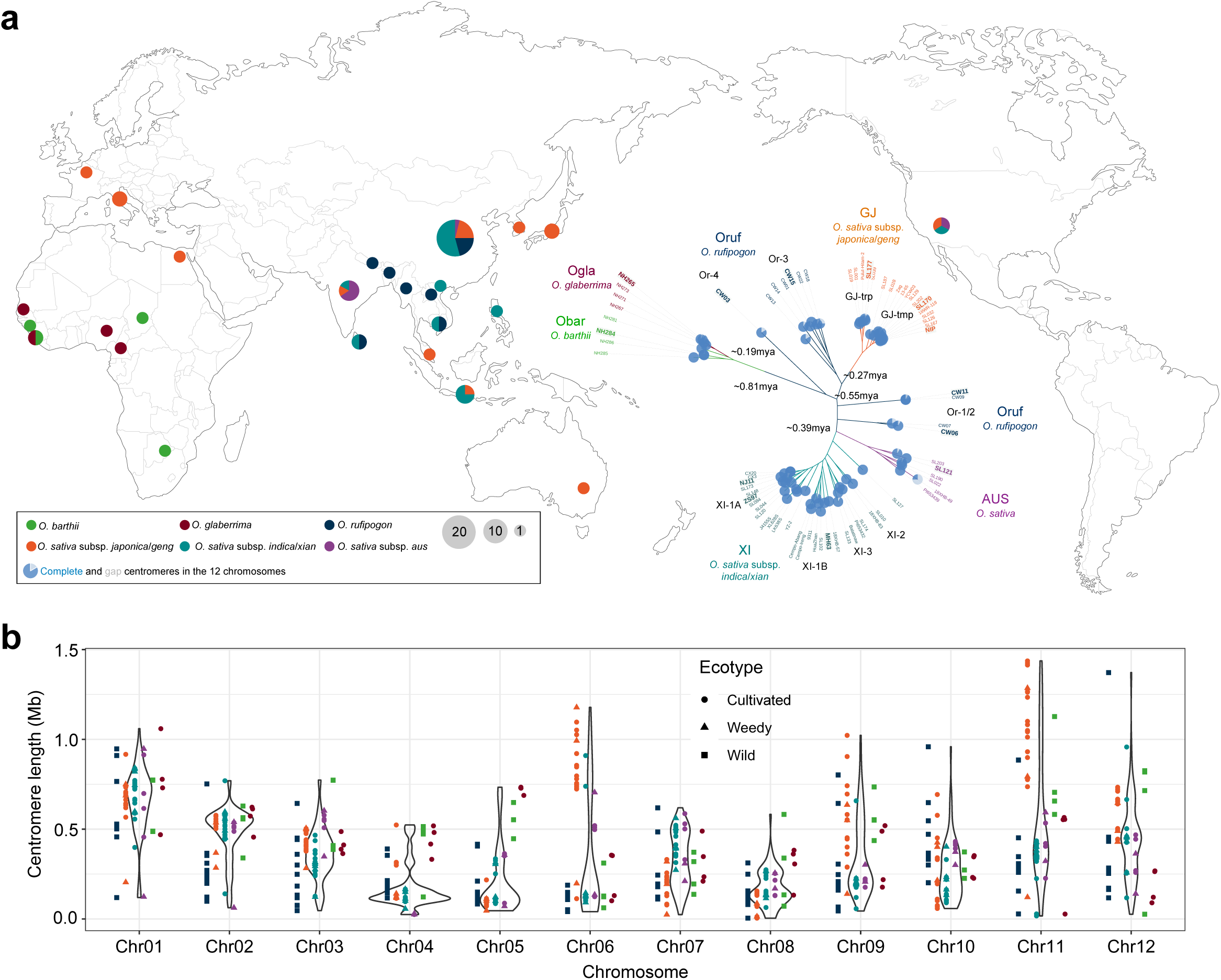
Centromere diversity of *Oryza* AA genomes. **a**, Geographic origin of the 70 rice accessions in the *Oryza* AA group, coloured by taxonomic group membership. A phylogenetic tree constructed using whole-genome SNPs is embedded in the map, with the branches and labels colored according to taxonomic information. Samples selected for performing CENH3 ChIP-seq are highlighted with bold labels on a gray background. Divergence times were adapted from Stein *et al*., 2018. **b**, Variation in centromere length across the 70 accessions per chromosome.

## Results

### Assembling rice centromeres

To fully capture the genomic diversity of rice centromeres, we collected 70 nearly complete sequences of *Oryza* AA genomes, incorporating newly assembled sequences from 67 rice genomes using PacBio HiFi and ONT long reads and three previously released T2T assemblies of Nipponbare (NIP)(Shang *et al*., 2023), Minghui63 (MH63) and Zhenshen97 (ZS97)(Song *et al*., 2021) (**Supplementary Table 1**). Briefly, the 70 rice AA genomes encompassed 11 from wild *O. rufipogon* (hereafter referred to as Oruf), 18 from *O. sativa* subsp. *Geng*/*japonica* (GJ), 25 from *O. sativa* subsp. *Xian*/*indica* (XI), six from *aus* (AUS), along with four wild from *O. barthii* (Obar) and four cultivated from *O. glaberrima* (Ogla) of African rice, collectively covering a wide spectrum of genetic diversity in Asian and African rice (**Fig. 1a**). In detail, Asian GJ rice samples comprised 13 accessions from subgroup temperate GJ (GJ-tmp), five from tropical GJ (GJ-trp), and XI included 12 from XI-1A, seven from XI-1B, two from XI-2 and six from XI-3 (**Fig. 1a**). Wild *O. rufipogon* population was composed of Or-1/2 (*n* = 4, closest ancestor of XI and AUS), Or-3 (*n* = 6, closest ancestor of GJ) and Or-4 (*n* = 1, basal group of all *O. sativa* samples).

Genome assemblies were generated using PacBio HiFi (36.0-fold coverage in average with a N50 length of ∼19.0 Kb) and ONT data (∼83.3-fold with a N50 length of ∼33.3 Kb) (**Supplementary Table 1**). We initially used Hifiasm to generate a highly accurate backbone of genome assembly, and then integrated the Verkko assembly to fill the gaps (Cheng *et al*., 2022; Rautiainen *et al*., 2023). After polishing using PacBio HiFi and NGS short reads, highly accurate assemblies were generated with an average consensus quality value (QV) of 64.4 (less than one base error per Mb)(**Supplementary Fig. 1**; **Supplementary Table 1**). Whole-genome synteny against the NIP reference assembly confirmed the absence of large structural errors. Base-resolution assembly quality assessment indicators revealed the platinum-level quality of the new assemblies. The average whole-genome GCI (Chen *et al*., 2024), an assembly continuity index derived from contig N50, was 71.1, surpassing previous gapless assemblies of MH63 (49.9) and ZS97 (54.1). Assembly quality indicators (AQI) (Li *et al*., 2023) for regional errors (R-AQI, 99.0) and structural errors (S-AQI, 98.1) further supported the high quality of the assemblies (**Supplementary Fig. 1**; **Supplementary Table 1**).

Centromere regions were delineated based on the presence of the rice centromere-specific satellite repeat sequence *CEN155* (also known as *CentO,* ∼155 bp in length) (Cheng *et al*., 2002) (**Supplementary Table 2**). In total of 839 centromeres were identified across the 70 genomes, with one exception on chromosome Chr07 in wild accession CW06, where no *CEN155*-defined centromere was found. Notably, 808 (96.2%) of these centromeres were contiguous without assembly gaps (**Fig. 1a**; **Supplementary Fig. 1**). Base accuracy assessments revealed that 66.2% of assemblies had "+inf" (infinite) QV scores for all 12 centromere sequences, indicating no erroneous bases in the centromere assemblies (**Supplementary Fig. 2a**). At the chromosome level, 92.6% of centromeres exhibited QV values greater than 60, corresponding to an estimated base accuracy exceeding 99.9999% (**Supplementary Fig. 2a**). Analysis using VerityMap identified few discordant *k*-mers between the centromere assemblies of cultivated accessions (*O. sativa* and *O. glaberrima*) and PacBio HiFi reads (Mikheenko *et al*., 2020), whereas a few potential erroneous *k*-mer clusters in wild accessions (*O. rufipogon* and *O. barthii*) were observed, likely due to the higher heterozygosity in the wild populations (**Supplementary Fig. 2b**). The average GCI score for all centromeres was 95.6 (**Supplementary Fig. 2**c), with only 39 centromeres showing potential gaps after stringent filtering of low-quality alignments (**Supplementary Table 3**). In other words, 95.2% of assembled centromeres were confirmed to be gapless. To avoid bias among assemblers when handling highly repetitive sequences (Logsdon *et al*., 2024), we compared the centromere assemblies generated by Hifiasm and Verkko, and found them to be highly consistent (**Supplementary Fig. 2**d). 414 (67.3%) out of 615 centromeres were completely assembled by both Hifiasm and Verkko, with *CEN155* array length differences less than 0.1%, and 538 (87.5%) had differences less than 1%. In summary, we have yielded a high-quality atlas of rice centromere assemblies with high accuracy and continuity.

### Centromere length variation

The L/S ratio (long arm/short arm) and centromere region synteny against the NIP reference assembly revealed the conserved positioning of *CEN155* satellite arrays for each chromosome among individuals (**Extended Data Fig. 1a**). Most chromosomes were metacentric or subcentric, whose centromeres are located near the middle of chromosomes, except Chr04 and Chr09, generally in line with the previous cytological characterization (Cheng *et al*., 2001). Coincidentally, the rice 45S ribosomal DNA (rDNA) repeat arrays were positioned on the short arm of acrocentric Chr09, consistent with the pattern observed in the human genome, implying the potential impact of the 45S rDNA array positioning on the evolution of chromosome karyotypes (Altemose *et al*., 2022; Shang *et al*., 2023).

Centromere lengths varied significantly among rice species, subspecies, individuals and chromosomes (**Fig. 1b**). Across individual genomes, centromere sequences occupied approximately 1.11% of the total genome, ranging from 0.57% to 1.82%, considerably smaller compared to the ∼3% (occupied by 171-bp alpha satellites) in the human genome (Altemose *et al*., 2022) and the ∼9.6% in the *Arabidopsis thaliana* genome (occupied by 178-bp satellites) (Naish *et al*., 2021)(**Supplementary Table 2**). Among chromosomes, we observed a positive correlation between centromere length and chromosome length, indicating that longer chromosomes tend to have longer centromere sequences (**Extended Data Fig. 1b**; **Supplementary Fig. 3**). Interestingly, this correlation was not significant in wild Asian rice and wild African rice groups, in contrast to cultivated groups (**Supplementary Fig. 3**). Despite the overall positive correlation with chromosome length, centromere length contributed minimally to the variation of chromosome size within the same chromosome, and even showed a negative correlation in some cases (**Supplementary Fig. 4a**). In contrast to the significant role of centromere length in chromosome size variation in *Arabidopsis thaliana* (Lian *et al*., 2024), TEs were identified as the primary source of chromosome size variation in rice (**Supplementary Figs. 4b and 4c**).

At the whole-genome level, Asian rice exhibited longer centromeres compared to African rice, although variations were observed across chromosomes (**Fig. 1b**). Among the different groups, as the outgroup of *O. sativa* and other *O. rufipogon*, CW03 from Or-4 had the longest whole-genome centromere sequence (∼6.94 Mb), followed by GJ accessions (∼5.58 Mb), while XI (∼3.73 Mb), AUS (∼4.58 Mb), wild Oruf (∼3.38 Mb) and cultivated African rice (∼5.00 Mb) exhibited shorter centromeres (**Fig. 1b**; **Supplementary Table 2**). Within the GJ group, GJ-tmp centromeres (∼5.93 Mb) were longer than those of GJ-trp (∼4.73 Mb). Different chromosomes showed distinct variations in lengths (**Fig. 1b**). For instance, GJ had significantly longer centromeres than other groups on chromosomes Chr06 and Chr11, potentially due to lineage-specific expansions. The three longest centromeres were all from chromosomes Chr11 of GJ-tmp, each exceeding 1.4 Mb in length. Interestingly, a 6-Mb inversion spanning the centromere and one nested 1-Mb inversion near the centromere on Chr06 were observed (**Extended Data Fig. 1c**). The smaller nested inversion likely originated in the common ancestor of GJ-trp and GJ-tmp, while the larger inversion occurred specifically in the GJ-tmp lineage. These structural difference in Chr06 centromeres between subspecies may contribute to the reproductive isolation in natural hybridization and low recombination rates in artificially recombinant populations (Wei *et al*., 2024).

### Centromere introgression and fission

Sequence similarity between intra-group centromeres was higher that that between inter-group centromeres, highlighting the distinct evolutionary trajectories between XI and GJ (e.g. Chr02) (**Extended Data Fig. 2a; Supplementary Fig. 5**). Notably, Chr05 centromeres showed exceptionally high similarity between GJ and XI but lower between GJ and AUS, supporting the hypothesis that a large Chr05 segment in the XI genomes, spanning the centromere, was introgressed from GJ during speciation or initial domestication (Wu *et al*., 2022) (**Extended Data Fig. 2b**). Aligning to GJ reference assembly NIP further supported these findings, revealing reduced structural variations in the flanking sequences of Chr05 centromeres across most XI samples, relative to other chromosomes. Additionally, Chr10 centromeres in subgroup GJ-trp showed high similarity to a subgroup of XI, aligning with the documented introgression events from XI to GJ-trp (Gong & Han, 2022) (**Supplementary Fig. 5**). Considering the frequent inter-subspecies hybridization in rice breeding, centromere introgression events affecting various chromosomes have been observed, including well-known cultivars like ZS97, MH63 and Kasalath (**Supplementary Table 4**). For instance, the Chr04 centromere in XI cultivar J4155S (alias Jing4155S) displayed a GJ-type centromere, in line with the historical records of the utilization of GJ (Nongken58 from Japan and Lemont from USA) in the breeding progress of J4155S (**Extended Data Fig. 2c**).

Although it is widely believed that the meiotic recombination in centromere regions is severely repressed, we analyzed the centromere sequences in sliding windows to explore potential centromeric rearrangement events. Representative centromere haplotypes were defined for each chromosome and then we categorized all the centromere blocks into different haplotypes based on pairwise similarity (**Supplementary Fig. 5**). Besides the whole-centromere introgression events above, rare centromere splitting or fission events were observed. Chr04 centromeres in XI, for example, exhibited two distinct types (X1 and X2) (**Supplementary Fig. 5**) and correspondingly the identity between X1 and X2 showed diagonal inconsistency in the heat-map plot, for which X1 was deleted from X2 with a shorter length (**Extended Data Fig. 2c**). Similarly, the Chr12 centromere of SL044 (Nantehao, one of the most essential backbone parents in the hybrid rice breeding), comprised a combination of two XI-type haplotypes (X3 and X4)(**Fig. 2a**). Collinearity among SL044, X3 (e.g. CW09) and X4 (e.g. NJ11) centromeres and internal sequence similarity within each centromere supported the structural combination (**Fig. 2b**). An identical (99.40%) intact LTR element *RETROSAT-2C* was identified both at the end of X3 and the beginning of X4 (**Fig. 2b**; **Supplementary Fig. 6**). Phylogenetic analysis using centromere upstream and downstream SNPs (∼1 Mb) indicated consistent topology among SL044 and the two haplotypes, where the flanking sequences of SL044 centromere was both close to haplotype X4 (e.g. NJ11), suggesting that the Chr12 centromere of SL044 likely did not arise from the fusion of X3 and X4 (**Fig. 2c**). To sum up, rapid evolution and innovation in rice centromeres could be driven by direct introgression events and large-scale rearrangements such as fissions, duplications and inversions, although these events were comparatively rare (**Fig. 2d**).

**Figure 2.**
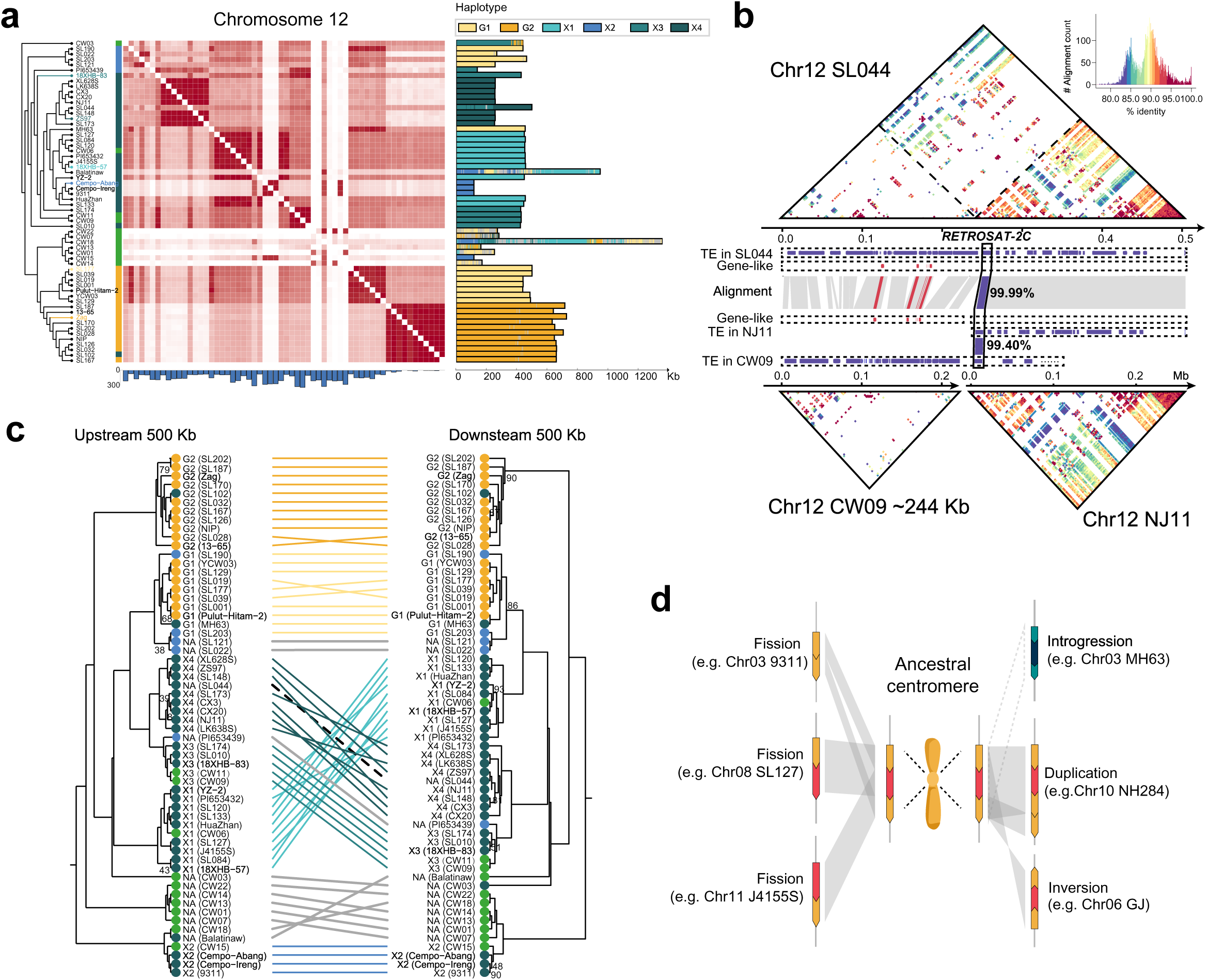
Centromere introgression and splitting. **a**,Centromere sequence similarity, structural variations in centromere flanking regions (compared to NIP), and siding-window haplotyping of Chr12 centromeres. Haplotypes are shown in different colors. **b**, StainedGlass sequence similarity heat maps of Chr12 centromeres from SL044, CW09 and NJ11, and their synteny. TEs (blue) and gene-like elements (red) are shown. Their commonly shared retrotransposon *RETROSAT-2C* is highlighted. **c**, Phylogeny of Asian rice accessions based on the upstream and downstream SNPs of the Chr12 centromere. Taxonomic information is represented by colored circles, with the position of SL044 marked by a dashed line. **d**, Schematic of centromere rapid alterations in structure, including introgression, duplication and fission or splitting.

### Satellite repeat variation

To precisely characterize the sequence compositions of rice centromeres, we annotated the *CEN155* satellites, TEs and potential genes. Across all centromeres, we identified a total of 1.46 million copies of *CEN155* satellites, predominantly distributed around 155 and 165 bp in length (**Supplementary Fig. 7**). The average copy number of *CEN155* satellites varied significantly among genomes, with GJ (∼28,210) showing a significantly higher count compared to XI (∼16,581), AUS (∼18,011) and wild Asian rice Oruf (∼15,203), similar to African rice Ogla (∼27,176) and Obar (∼29,736). These variations correlated with differences in centromere length, but markedly fewer than the abundance of *CEN180* satellites in *Arabidopsis thaliana* centromeres (∼66,131) (Naish *et al*., 2021). The *CEN155* satellite repeats formed long tandem arrays, often exhibiting bias in strand orientation (**Fig. 3a**). Significant divergence in strand preferences was observed among groups on each chromosome, with distinct patterns observed between chromosomes in GJ and XI (**Fig. 3a**).

**Figure 3.**
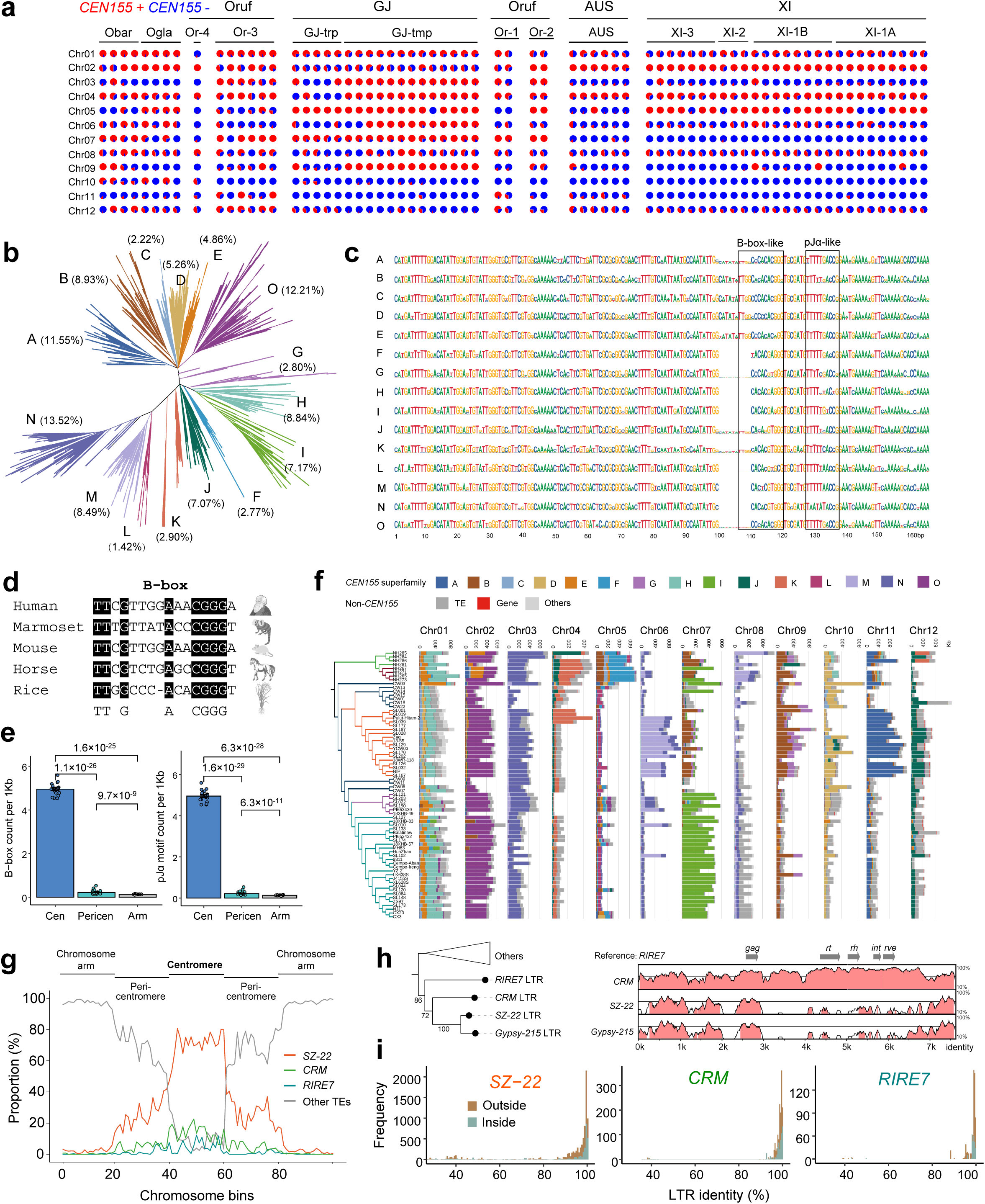
Variation of *CEN155* sequences and centrophilic TEs in centromere regions. **a**, Pie charts illustrating the *CEN155* proportion on the forward (red, +) or reverse (blue, -) strand across chromosomes (rows) and accessions (columns). **b**, A maximum-likelihood phylogenetic tree of representative *CEN155* sequences (*n* = 6,665). Fifteen superfamilies are clustered and colored. **c**, Consensus sequences of the fifteen *CEN155* superfamilies. B-box and pJα-like motifs are highlighted. **d**. Alignment of B-box motifs across human, marmoset, mouse, horse and rice. **e**, Enrichment analysis of B-box in different genomic regions of the rice genome. Significance values were calculated using Student’s *t* test. **f**, Centromere compositions across the 70 rice genomes. Branch colors in the phylogeny tree (left) indicate taxonomic groups. **g**, Abundance enrichment of centrophilic retrotransposons (*SZ-22*, *CRM*, *RIRE7*) in centromere, peri-centromere and chromosome arm regions. Each chromosome is shown in 100 bins, where the window sizes are adjusted depending on the length of each region. **h**, Phylogenetic relationship and sequence similarity among centrophilic LTRs. Bootstrap values of 1000 replicates are shown near the branches. **i**, Insertion ages, measured by sequence identity between the two flanking LTR sequences within each intact *SZ-22*, *CRM* and *RIRE7*, comparing inside and outside the *CEN155* arrays.

To comprehensively profile the structures of centromere monomers and trace the evolutionary footprints, we clustered 6,665 representative *CEN155* sequences (140-170bp in length and >30 occurrences in the centromere collection) into 15 families based on phylogeny (**Fig. 3b**). Low-frequency and divergent monomers were assigned according to their least edit distances to the clustered *CEN155* sequences, and finally 15 superfamilies (SFs) were termed as superfamily A (SF-A) to O (SF-O), with SF-N being the most prevalent across the 70 rice genomes, comprising approximately 200 thousand occurrences, while SF-L exhibited the least abundance with fewer than 20 thousand copies (**Fig. 3b**). Each superfamily can be distinguished by specific motif sequences, which could serve as genetic markers for categorizing *CEN155* satellites (**Fig. 3c**). Notably, a 10-bp structural variation (CATATATTGG) was present in SF-B to SF-E, but was mostly absent in superfamilies from SF-F to SF-O (**Supplementary Fig. 8a**). Pairwise edit distances among monomers within the same superfamily were generally lower than those between *CEN155* satellites from different superfamilies, but the results still showed high diversity of monomers within each superfamily (**Supplementary Fig. 8b**).

The presence of B-box-like and pJα-like motifs can be detected in rice consensus *CEN155* sequences, analogous to motifs found in human centromeres involved in the kinetochore assembly (Rosandic *et al*., 2006; Rice, 2020), despite human and rice belonged to different kingdoms (**Fig. 3c**). Alignment with known B-box-like motifs from various species (marmoset, mouse, house) and rice against the canonical B-box motif in human (5’-*TTCGTTGGAA*CGGGA-3’) refined the conserved B-box motif to 5’-TT*G****A**CGGG-3’ (**Fig. 3d**) (Kipling *et al*., 1994; Kugou *et al*., 2016; Rice, 2020; Cappelletti *et al*., 2023). The presence of B-box and pJα-like elements was significantly enriched in centromere regions (**Fig. 3e**), with divergence among superfamilies primarily attributed to variations at base positions A and C, potentially influencing kinetochore function. Although the 10-bp PAV deleted the TT*G of B-box in the alignment, another TTGG short motif before the 10-bp PAV was present as a substitute for most superfamilies.

Centromeres on different chromosomes exhibited distinct compositions (**Fig. 3f**), similar as the centromere structure of *Arabidopsis* genome (Wlodzimierz *et al*., 2023). For instance, SF-A, D, H, J, and O predominated on Chr11, Chr10, Chr01, Chr07 and Chr02, respectively, illustrating chromosome-specific distribution patterns. Proportions of the 15 superfamilies also varied among taxonomic groups. SF-I was nearly absent in Obar and GJ but comprising 17.5% in XI and AUS genomes, primarily on Chr07 (**Fig. 3f**). Notably, size expansions of GJ centromeres on Chr06 and Chr11 were attributed to increased SF-N and SF-A monomers, respectively, compared to other groups. These whole-genome constitutive differences in superfamilies among groups were predominately influenced by specific chromosomes, harboring chromosome-specific *CEN155* satellites (**Fig. 3f**).

### TE invasion in centromeres

Centromere satellites typically arrange head-to-tail to form satellite arrays but such homogenized structures are occasionally disrupted by TE insertions in plant centromeres (Naish & Henderson, 2024). We conducted an annotation of inter-*CEN155* sequences to profile the TE insertion landscape in rice centromeres. Approximately ∼85.1% of non-*CEN155* centromeric sequences were composed of TEs. TE proportions within centromere regions (∼22.2%) were significantly lower than those across the whole genome (∼51.9%), suggesting stronger purifying selection against TE insertion in centromeres compared to chromosome arms (**Supplementary Fig. 9**; **Supplementary Table 2**). Despite shorter centromeres, XI (∼1.01 Mb) and AUS (∼1.25 Mb) genomes exhibited more TEs within centromeres compared to GJ (∼0.72Mb). About 77.8% (in length) of TEs in the centromeres belonged to long-terminal repeat (LTR) retrotransposon superfamily *Gypsy*, much more intensive than the whole-genome *Gypsy* density (∼40.7%), similar as *Arabidopsis* centromeres (Naish *et al*., 2021) (**Supplementary Fig. 9**).

Across all the 70 genomes, we identified a total of 210,672 intact LTRs and 1,042,341 soloLTRs, with 5,128 intact LTRs and 8,366 soloLTRs located within centromeres. The ratio of intact LTRs *verse* soloLTRs was significantly higher within centromeres than outside (0.61 *versus* 0.20), yet much lower than in the *Arabidopsis* genomes (∼2.5) (Wlodzimierz *et al*., 2023). Of all the centromeric intact LTRs, *SZ-22* (*n* = 3673, covering 8.1% of the total centromere length), *CRM* (*n* = 714, 1.8%) and *RIRE7* (*n* = 257, 0.6%) were the most abundant three families, collectively accounting for 90.6% (**Fig. 3g**; **Supplementary Table 2**). Additionally, 26.0% and 22.1% of *SZ-22*, 31.9% and 27.4% of *CRM*, and 30.2% and 18.2% of *RIRE7* intact elements across the whole genome were significantly enriched within the centromere and pericentromeric regions, respectively, suggesting evolutionary coupling and pairwise selection between centrophilic LTRs and satellite repeats (**Fig. 3g**). Phylogeny of the two flanking LTR sequences of intact retrotransposons revealed close relationships among the three families, while distant when inferring from the internal sequences (**Fig. 3h**; **Supplementary Fig. 10a**). Although *SZ-22* was the most dominant LTR family in centromeres, it was non-autonomous which lacked reverse transcriptase (*rt*), RNase H (*rh*) and integrase (*int*) genes, with a significantly shortened intact length (∼4.5 Kb) compared to *RIRE7* and *CRM* (∼7.7 Kb) (**Fig. 3h**; **Supplementary Fig. 10b**).

Centromeric *SZ-22* insertions were significantly younger than those in chromosome arms (*P* value = 7.5e-4, Wilcoxon test), although a certain number of ancient *SZ-22* insertions in centromeres were retained (**Fig. 3i**). Using the sequences flanking *SZ-22* inserted elements, we localized the integrated sites within the *CEN155* consensus repeat. A total of 2,258 *SZ-22* insertions were found throughout the full length of *CEN155* sequence, and the highest frequency of insertion occurred at positions 100 to 120 base, coincided with the location of B-box motif (**Fig. 3c**; **Supplementary Fig. 11**). As mentioned above that frequent reversal events between forward and reverse strand were observed within the *CEN155* arrays (**Fig. 3a**), we noticed that the junction regions of strand reversal were enriched with TE insertions, where TEs were observed in the 1790 out of all the 2133 (83.9%) junction regions, implying a potential role of TE insertion in triggering strand reversal of *CEN155* satellites. Taken together, these features reveal extensive and recurrent invasion of TEs in the *CEN155* satellite arrays, shaping the structural and evolutionary dynamics of rice centromeres. Furthermore, we annotated a total of 535 candidate centromeric gene models within centromeres across the 70 genomes, none of which showed evidence of gene expression.

### Non-canonical satellite repeats

In addition to TE invasion within the *CEN155* arrays, we identified the presence of 6,809 short intervening sequences dispersed between adjacent canonical *CEN155* satellites repeats (termed as *sati* for non-canonical satellite repeat as short intervals of canonical centromeric repeats) (**Extended Data Fig. 3a**). Some of these intervals were separated by a similar number of *CEN155* satellites, where they formed CEN155+ arrays. The abundance and size of *sati* sequences varied among groups and chromosomes (**Extended Data Fig. 3b**; **Supplementary Table 5**). Importantly, the high-frequency intervals were predominantly derived from the canonical *CEN155* sequences (**Extended Data Fig. 3c**). In three accessions of Or-3 wild Asian rice, a 207-bp interval (termed as *sati207*) between *CEN155* satellites on Chr11 was observed 147 times, which indicated group-specific centromeric variation (**Supplementary Table 5**). The *sati207* sequence consisted of two replicates of 104-bp satellites, but differed from *sati104* (**Extended Data Fig. 3c**). Satellite *sati99* on Chr03, had expanded in cultivated *O. glaberrima* compared with wild *O. barthii* (**Supplementary Table 5**).

Furthermore, in African rice, we identified 531 *sati94* sequences with an approximate length of 94 bp, specially enriched in Chr10 centromeres, and particularly expanded in wild *O. barthii* (e.g. NH284 in **Extended Data Fig. 3d**). Interestingly, structural comparison indicated that two clusters of *sati94* in NH284 (Obar) likely emerged through duplication from a NH265-like centromere (Ogla)(**Extended Data Fig. 3d**). CENH3 ChIP data confirmed the function of CEN155+ arrays composed of canonical *CEN155* and non-canonical *sati94* satellites(**Extended Data Fig. 3d**). Despite the similarity in the composition and structural organization of nearby *CEN155* satellite arrays, the distance distribution between every two adjacent *sati94* satellites within each of the three blocks revealed distinct arrangements, suggesting differential local duplication and homogenization progresses (**Supplementary Fig. 1**2a). Phylogenetic analysis further elucidated the evolutionary trajectory of *sati94* sequences within the three CEN155+ arrays and the mixed distribution of *sati94* repeats confirmed the sharing of common ancestor as well as individual-specific expansion (**Supplementary Fig. 12b**).

### Multi-layer nested structure of centromeres

To uncover the structural characteristics of *CEN155* arrays, we employed a progressive compression strategy simulating the expansion progress in reverse (**Fig. 4a**). This approach revealed localized sequence homogenization at monomer, dimer, and multimer scales, where the number of homogenization blocks decreased exponentially with increasing repeat copy size, although the homogenization patterns varied among groups and individuals (**Fig. 4b**; **Supplementary Fig. 13**). Across all centromeres, 59.7% (8.77 million) of *CEN155* satellites were found within 36,661 monomer homogenization regions (moHRs), defined as regions with >5 consecutive satellites from the same superfamily (**Supplementary Table 6**). *S*uperfamiles A (78.2%), I (95.0%), J (80.5%), K (78.6%) and O (83.7%) exhibited higher propensity to form moHRs, contrasting sharply with SF-C (0.5%) and G (0.9%). Chromosomes Chr02, Chr03, Chr07 and Chr12 displayed over 75% of their *CEN155* satellites organized in moHRs, suggesting higher homogeneity compared to other chromosomes (**Fig. 4c**). Divergence of monomer homogenization among groups and individuals was in line with their abundance in each family (**Fig. 3f**; **Supplementary Fig. 13a**). The longest moHR, composed of 5,288 repeats of SF-A on chromosome Chr11 in NIP, spanned approximately 865 Kb. Although *CEN155* satellites within a moHR belonged to the same SF, finer substructures could be observed similar to the classic HOR structure in the human genome, where an array composed of divergent satellites was repeated for several times (Altemose *et al*., 2022). We took the annotation of two centromeres as an example (**Fig. 4d**). In the centromere of chromosome Chr10 in GJ sample YCW03, despite SF-D being the most abundant, interruptions by SF-C satellites within SF-D arrays hindered direct sequence homogenization, which was not reflected from the sequence similarity directly. Satellites from SF-J were arranged head-to-tail into the largest homogenized moHR in the centromere, composed of 271 *CEN155* repeats (**Fig. 4d**). In the largest three SF-J moHRs longer than 100 satellites, the inner identity dot plots within some moHRs exhibited a similar pattern as HOR arrays. The most typical structure (iii) involved HORs approximately 11 satellites long, with a prominent *k*-mer distance spectrum peak at 1706 bp. In the centromere of chromosome Chr01 in CW03, SF-E satellites accounted for 65.1% of all *CEN155* repeats. Analysis of the four moHRs within this region revealed extensive local homogenization, evident from nearly identical *CEN155* sequences observed in the identify dots and *k*-mer distance spectrum among satellites (**Fig. 4d**).

**Figure 4.**
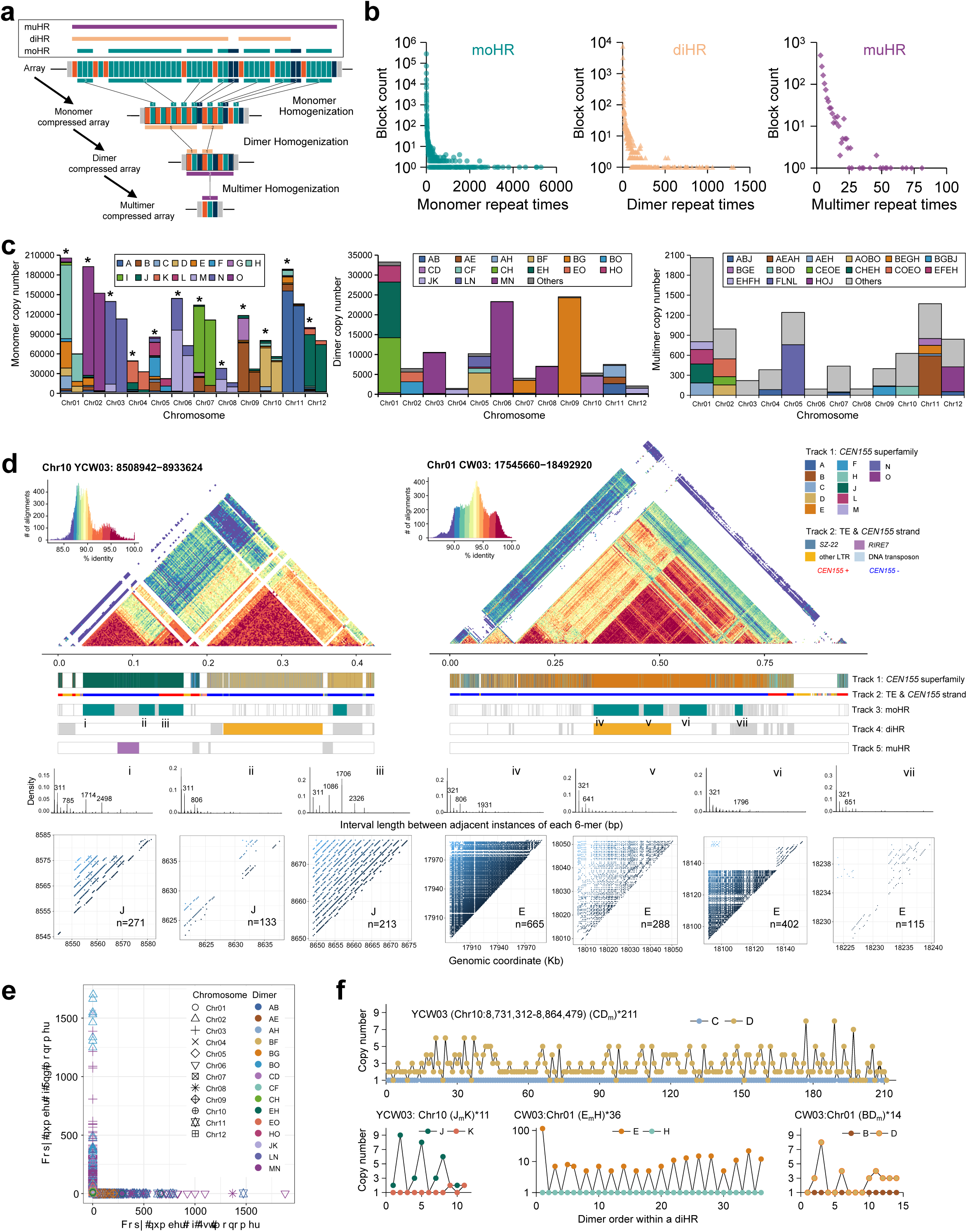
Multi-layer nested structure of rice centromeres. **a**, Schematic illustrating a progressive compression strategy to decompose the structure of *CEN155* arrays. Identical adjacent monomers, dimers and multimers are compressed stepwise, defining local homogenization blocks as moHRs, diHRs and muHRs, respectively. **b**, Distribution of replicate times for monomers, dimers and multimers. **c**, Structural composition of moHR (left), diHR (middle) and muHR (right) in centromeres across chromosomes. The asterisks denote the superfamily composition of all *CEN155* and the other stacked columns represent *CEN155* composition within moHRs. **d**, Centromere structures of two exemplar chromosomes. Annotation tracks display information including superfamily, LTR, strand, moHR, diHR and muHR. Zoomed-in views of seven moHRs illustrate fine HOR-like structures. **e**. Biased copy number distribution for the two monomers within a dimer. **f**, Regularity in the copy number for the two monomers from a dimer along the diHR.

Compressing each moHR into a monomer resulted in a simplified centromere string (**Fig. 4a**). We identified 68,370 dimer arrays in total, notably 4,548 long arrays containing at least 5 consecutive dimer repeats, totaling 135,920 dimers, termed as dimer homogenization regions (diHRs), with an average of 29.9 dimers per diHR (**Fig. 4b**; **Supplementary Fig. 13b**; **Supplementary Table 7**). The longest array was located on chromosome Chr09 of SL187 (GJ) with 1,308 repeats of BG dimer (**Fig. 4b**). The most frequent dimer arrays were MN arrays (40,218 dimers in 754 diHRs), followed by BG, EH and CH dimer. Differential compositions of diHRs were evident across chromosomes (**Fig. 4c**). For instance, MN dimers predominated on chromosomes Chr03, Chr06 and Chr08, while BG dimers were prominent on chromosomes Chr07 and Chr09 (**Fig. 4c**). The assembly of dimers exhibited non-random patterns, with the most abundant 15 dimers collectively accounted for 96.1% of all dimers in diHRs. A similar pattern of dimer organization has been observed in the human alpha satellites, which was assumed to be linked to the presence of B-box or pJα motif (Rice, 2020). We examined the divergence sites between the two monomers for each of the top 15 dimers and identified significant divergence (pairwise allele frequency difference greater than 0.8) at conserved B-box and pJα sites for most dimers, implying the potential relationship between featured motif sequences and satellite dimerized organization (**Extended Data Fig. 4**).

In human centromeres, satellites with and without a B-box motif alternate to form a B-box/no-B-box dimer (Rice, 2020). Interestingly, we observed a significant bias in monomer copy number of dimers, where one monomer showed a relatively constrained and stable copy number (almost always 1) while the other exhibited variable copies (forming moHRs) within a diHR, represented as XY*_m_*or X*_m_*Y (where *m* indicates multiple copies) (**Fig. 4e**). For instance, the centromere of chromosome Chr10 in YCW03 featured four diHRs, with the largest composed of 211 CD repeats. The copy number of satellite SF-C was fixed at one, while SF-D had variable copies ranging from one to eight (**Fig. 4f**). Similarly, another diHR with the JK dimer displayed great variation in SF-J copy numbers but consistently one copy for SF-K. On chromosome Chr01 in CW03, the largest dimer repeat diHR comprised 36 repeats of EH, with monomer SF-E ranging from one to 115 copies, contrasting with a fixed one copy for SF-H (**Fig. 4f**). However, this bias in monomer copy number variation for a dimer was not consistently observed across the whole genome scale (**Supplementary Fig. 14**). As mentioned above, chromosomes Chr03, Chr06 and Chr08 were predominately occupied by MN dimers. However, they exhibited differential copy number biases, with Chr03 primarily showing MN*_m_*, while Chr06 and Chr08 displayed a mixture of MN*_m_* and M*_m_*N (**Supplementary Fig. 1**4). Distinct biases were also noted among groups. For instance, the BO bias was reversed in XI compared with other groups (**Supplementary Fig. 14**). Although the rules governing monomer homogenization within dimers were not universally unified across the whole-genome or chromosome scale or among groups, the local bias was evident (**Supplementary Fig. 15**). Furthermore, in addition to the repetitive occurrences of the same dimer within a diHR, the copy number bias was repeated locally and commonly, strongly suggesting localized duplication as an approach for centromere expansion (**Fig. 4f**; **Supplementary Fig. 15**).

Compressing each diHR into a single dimer yielded a total of 1,383 multimer homogenization regions (muHRs), comprising 418 distinct repeat structures and 8,768 multimer repeats, averaging 6.34 copies per muHR (**Figs. 4a and 4b**; **Supplementary Fig. 13c**; **Supplementary Table 8**). Eight muHRs contained more than 50 replicates of multimer, with the most repetitive multimer being HOJ with 81 copies (**Fig. 4b**; **Supplementary Fig. 16**). The multimers FLNL, AEAH and HOJ were the most frequent, with 754, 590 and 373 repeats, respectively, predominantly chromosomes Chr05, Chr11 and Chr12 (**Fig. 4c**). In summary, we propose a multi-layer nested model of the rice centromere using a progressive compression strategy. Our findings suggest that centromere expansion occurred through local duplication of satellites at the monomer, dimer, multimer and their combination levels, consequently leading to the intricate complexity of the centromere.

### Centromere divergence

Distinct variations in centromere composition and structure are evident among different chromosomes and taxa, as mentioned above. However, the evolutionary dynamics and trajectories at the individual satellite scale remained largely unexplored, either in rice or other species, due to the extremely high similarity among satellites and the challenges in high-resolution synteny characterization among centromeres through direct sequence alignments. Here, we developed a method, SynPan-CEN, designed to identify syntenic satellite pairs among centromeres, by leveraging satellite clustering information and pairwise genetic distances (**Fig. 5a**; **Supplementary Fig. 16**). Initially, we identified syntenic *CEN155* pairs relative to NIP (GJ-tmp) centromeres in the other rice genomes. The synteny ratio decreased rapidly with increasing genetic divergence, with approximately 68.0% and 55.3% of *CEN155* satellites found to be syntenic within GJ-tmp and GJ-trp, respectively, significantly higher than between NIP and AUS (19.0%) or XI (17.4%)(**Supplementary Fig. 17**). A substantially higher syntenic relationship was observed between XI and GJ on chromosome Chr05 (61.7%), compared to other chromosomes (22.2%) (**Supplementary Fig. 18**), in line with the hypothesized introgression events from GJ to XI (Wu *et al*., 2022).

**Figure 5.**
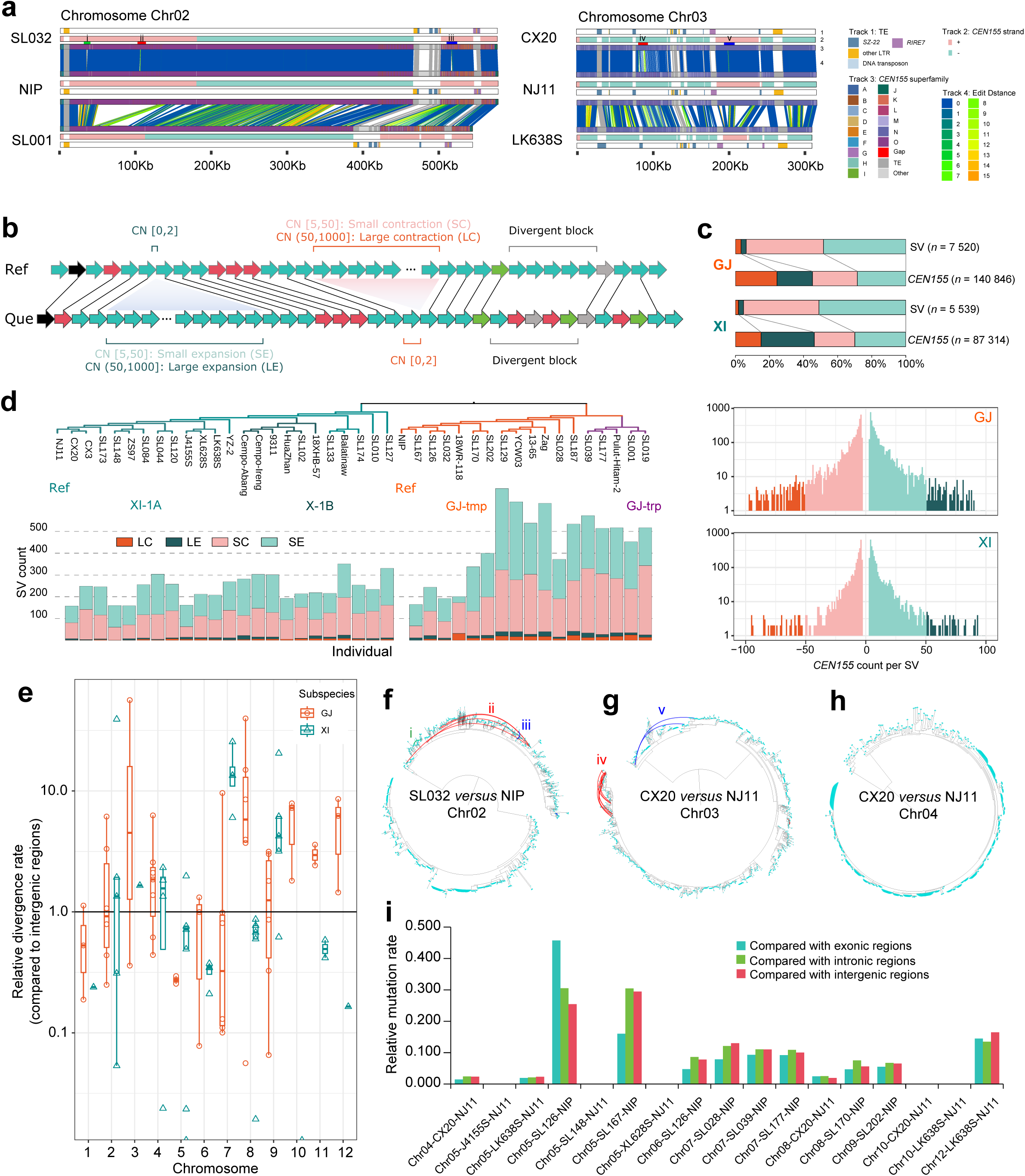
Sequence divergence and mutation rate of rice centromeres. **a**, Centromere synteny inferred using SynPan-CEN. **b**, Schematic of centromere structural variations (CSVs), including contraction and expansion. **c**, CSVs in centromeres of GJ and XI compared to the corresponding reference assemblies NIP and NJ11, respectively. **d**. CSVs in individual genomes associated with the phylogenetic kinship. **e**, Relative sequence divergence in centromeres compared to intergenic regions, estimated from syntenic *CEN155* pairs. **f**, Phylogenetic analysis of *CEN155* sequences from Chr02 centromeres of SL032 and NIP, indicating non-ortholog syntenic pairs (red, blue and green links). **g**, Phylogenetic analysis of *CEN155* sequences from Chr03 centromeres of CX20 and NJ11, indicating non-ortholog syntenic pairs (red and blue links). **h**, Phylogenetic analysis of *CEN155* sequences from Chr04 centromeres of CX20 and NJ11, indicating absolutely orthologous pairs of syntenic *CEN155* satellites. **i**, Relative mutation rates for 17 centromere pairs with absolutely orthologous *CEN155* satellites, compared to non-centromere regions.

We further examined the syntenic relationships of satellites between centromeres, to investigate genomic footprints of local expansion and contraction in GJ centromeres relative to the reference NIP and XI centromeres compared to the reference NJ11 (**Fig. 5b**). We defined centromere structural variations (CSVs) as instances where *CEN155* satellite copy numbers showed large expansions (LE, >50 but <1000 copies) or contractions (LC), and small expansions (SE, <50 but >5 copies) or contractions (SC). Across 17 and 23 samples compared to GJ and XI references, we identified 7,520 and 5,539 CSVs, encompassing approximately 140.8 thousand and 87.3 thousand copies of *CEN155* satellites, respectively (**Fig. 5c**; **Supplementary Table 9**). Large LCs and LEs accounted for 5.7% of all CSVs in count (∼14.7 and ∼13.7 per GJ sample, ∼4.5 and ∼7.2 per XI sample), yet represented 45.6% of CSV satellite copy number variations. On average, 17.0 and 17.9 copies of *CEN155* were included per contraction and expansion variation, respectively. The exponential distribution of CSVs in terms of size was in line with the length distribution spectrum of SVs on the chromosome arms (Qin *et al*., 2021; Shang *et al*., 2022)(**Fig. 5c**). Detected CSV counts ranged from 165 (SL167/Jindao 1, alias B236 in the 3000 Rice Genome Project) to 697 (SL129/Keluoduo B, alias B179 from France) for GJ, and from 158 (CX20) to 351 (Balatinaw, a glutinous and red heritage rice) for XI, respectively, depending on the genetic distances to their respective reference genomes (**Figs. 5a & 5d**). Notably, In XI, the accession closest to NJ11 in centromeric kinship was CX20, consistent with their closest evolutionary relationship at the whole-genome scale (**Figs. 1a, 5a and 5d**). CX20 is a weedy rice accession, presumed to have originated from the de-domestication of the modern cultivar NJ11, widely cultivated throughout South China 30 years ago (Qiu *et al*., 2020).

Centromere regions are known for their accelerated evolutionary rates compared to the chromosome arms, characterized by significant compositional differences observed among closely related species or populations (Bensasson, 2011; Wlodzimierz *et al*., 2023; Logsdon *et al*., 2024; Minton, 2024). For instance, human centromeres exhibit higher mutation rates (1.1∼4.1 times) in the pericentromeric regions compared with non-human primates and the active HOR arrays were presumed to have an increased mutation rate greater than ten times (Logsdon *et al*., 2024). The high repetitiveness of centromere regions complicates biologically meaningful sequence alignments and comparisons. Instead of traditional sliding window alignment strategy, we utilized the satellite-resolved synteny framework between centromeres to profile the evolutionary rates of rice centromeres. Considering the nearly complete turnover of *CEN155* arrays between XI and GJ for certain chromosomes (**Fig. 3f**; e.g. Chr07), comparison between GJ and XI were omitted, focusing instead on the divergence between GJ samples and the reference assembly NIP, and between XI samples and the reference assembly NJ11, respectively. To eliminate false positives from poor alignment, we screened 91 chromosome pairs with a syntenic ratio exceeding 0.9 (**Supplementary Table 10**). We defined “divergence rate” to quantify the sequence difference between alignable syntenic satellites and “mutation rate” to quantify the *de novo* changes in absolute orthologous satellite pairs. In other words, syntenic satellites are not always orthologous. We calculated the biological edit distances (counts of mismatches and gap openings) between DNA sequence alignments from the gene exonic, gene intronic, intergenic regions, and the *CEN155* satellite arrays. Due to the limited evolutionary timescale within GJ and XI groups, we focused on relative evolution rates in centromeres compared to chromosome arms, rather than the absolute evolution rate over generation. Among the 91 chromosome pairs, centromere divergence rate averaged 4.1 times higher than those in chromosome arms, though varied widely (0 to 56.4), depending on chromosomes and individuals (**Fig. 5e**). We zoomed in on the *CEN155* arrays to investigate the source of genetic divergence. Despite similar centromere composition and structure in the compared chromosome pairs, a few of local syntenic *CEN155* pairs were not orthologs. For instance, SL032 and NIP on chromosome Chr02 displayed nearly identical centromeres with only two small CSVs (one 11-copy SC and one 7-copy SE) and one tiny CSV (3-copy contraction) (**Fig. 5a**). Notably, a few of syntenic *CEN155* pairs around the CSVs exhibited higher ED values compared to other pairs. To exam whether the high sequence divergence was contributed from high mutation rate or exaggerated genetic distance, we built the phylogeny using all *CEN155* satellites and found that *CEN155* satellites in the high-ED syntenic pairs were not clustered monophyletically, but instead located distantly in the phylogeny, which suggested that such pairs were not orthologous but paralogous, and thus could not be used to measure the mutation rate (**Fig. 5f**). Besides dispersal high-ED syntenic pairs, we also observed clustered syntenic *CEN155* pairs with evaluated ED values among centromeres. For example, comparing the centromeres of NJ11 and CX20 on chromosome Chr03 revealed a cluster of high-ED pairs (**Fig. 5a**). Phylogenetic analysis confirmed these pairs were distantly related due to non-orthologous origins (**Fig. 5g**). Additionally, the *CEN155* composition differed significantly, with NJ11 containing a higher proportion of *CEN155* satellites from super-family O, possibly due to specific gene conversion or segmental replacement from homologous regions, as evidenced by comparison with outgroup sample LK638S (XI-1B). Therefore, to mitigate the significant impact of overestimated genetic distances between potential paralogous syntenic pairs in the mutation rate analyses, chromosome pairs were excluded where syntenic *CEN155* syntenic satellite pairs were determined non-orthologous by the *CEN155* satellite phylogeny (**Fig. 5h**; **Supplementary Table 10**). Analysis of the 17 chromosome pairs with uniformly orthologous *CEN155* satellites revealed significantly lower mutation rates (∼0.11 times) compared to chromosome arms (**Fig. 5i**). In particular, three centromeres on Chr05 and two on Chr10 in XI samples mirrored reference centromeres in NJ11 (**Supplementary Fig. 19**). In summary, based on the pan-centromere analysis above, rice centromeres exhibit higher sequence divergence rates than chromosome arms, and this divergence primarily stems from non-ortholog gene conversion or segmental substitution rather than *de novo* mutations. Conversely, *de novo* mutation rates in rice centromeres are notably lower than those in chromosome arms, which may be explained by the hyper-methylation nature of centromeres.

### Epigenetic variation and repositioning of centromeres

We conducted ChIP-Seq experiments using specific anti-CENH3 antibody to map the functional centromeres in rice, revealing widespread repositioning and neocentromere formation across multiple genomes (**Supplementary Figs. 20-29**). On average, the CENH3-enriched sequences spanned approximately 8.22 Mb across the thirteen profiled genomes, significantly exceeding the size of centromere regions defined by *CEN155* arrays (∼4.35 Mb), in contrast to the homeostasis of CENH3 loading in *Arabidopsis* (Naish *et al*., 2022; Wlodzimierz *et al*., 2023) and human centromeres (Logsdon *et al*., 2024) but comparable to observations in maize centromeres (Chen *et al*., 2023) (**Supplementary Table 11**). Detailed analysis revealed that on 73 out of 156 chromosomes, the CENH3-enriched regions exceeded 1.5 times the size of the *CEN155*-defined centromere regions. Notably, chromosomes such as Chr01, Chr02, Chr03 and Chr07 exhibited highly consistent positional matching between CENH3-defined and *CEN155*-defined centromere regions, while other chromosomes showed significantly longer CENH3-defined centromeres (**Supplementary Fig. 30**).

To investigate factors related to centromere repositioning and neocentromere formation, we examined the pairwise correlations for each chromosome among chromosome length, *CEN155* array length, CENH3 occupancy length, TE density within and outside the *CEN155* array, and neocentromere formation tendency (NFT) (defined by the length ratio between CENH3 enrichment region and *CEN155* array)(**Fig. 6a**; **Supplementary Figs. 30-36**). Generally, for a given chromosome, the diversity in chromosome length was attributed from TE expansion (**Supplementary Fig. 4)**. Longer chromosome tended to exhibit more intense TE insertion within their centromeres, coupled with shorter *CEN155* array (**Fig. 6a**; **Supplementary Figs. 31 and 32**), suggesting the erosion of the *CEN155* array by TE activity. Longer chromosomes also required extended functional centromeres as defined by CENH3 occupancy, particularly noticeable in chromosomes with significant length variation (**Fig. 6a**; **Supplementary Fig. 33**). We observed positive relationship between centromeric TE density and NFT and negative correlations between the *CEN155* array length and NFT (**Fig. 6a**; **Supplementary Figs. 34 and** 35). Consequently, longer chromosomes among individuals for a given chromosome have higher density of TE insertion, shortened *CEN155* arrays and a greater propensity for outward expansion (**Fig. 6a**; **Supplementary Fig. 36**).

**Figure 6.**
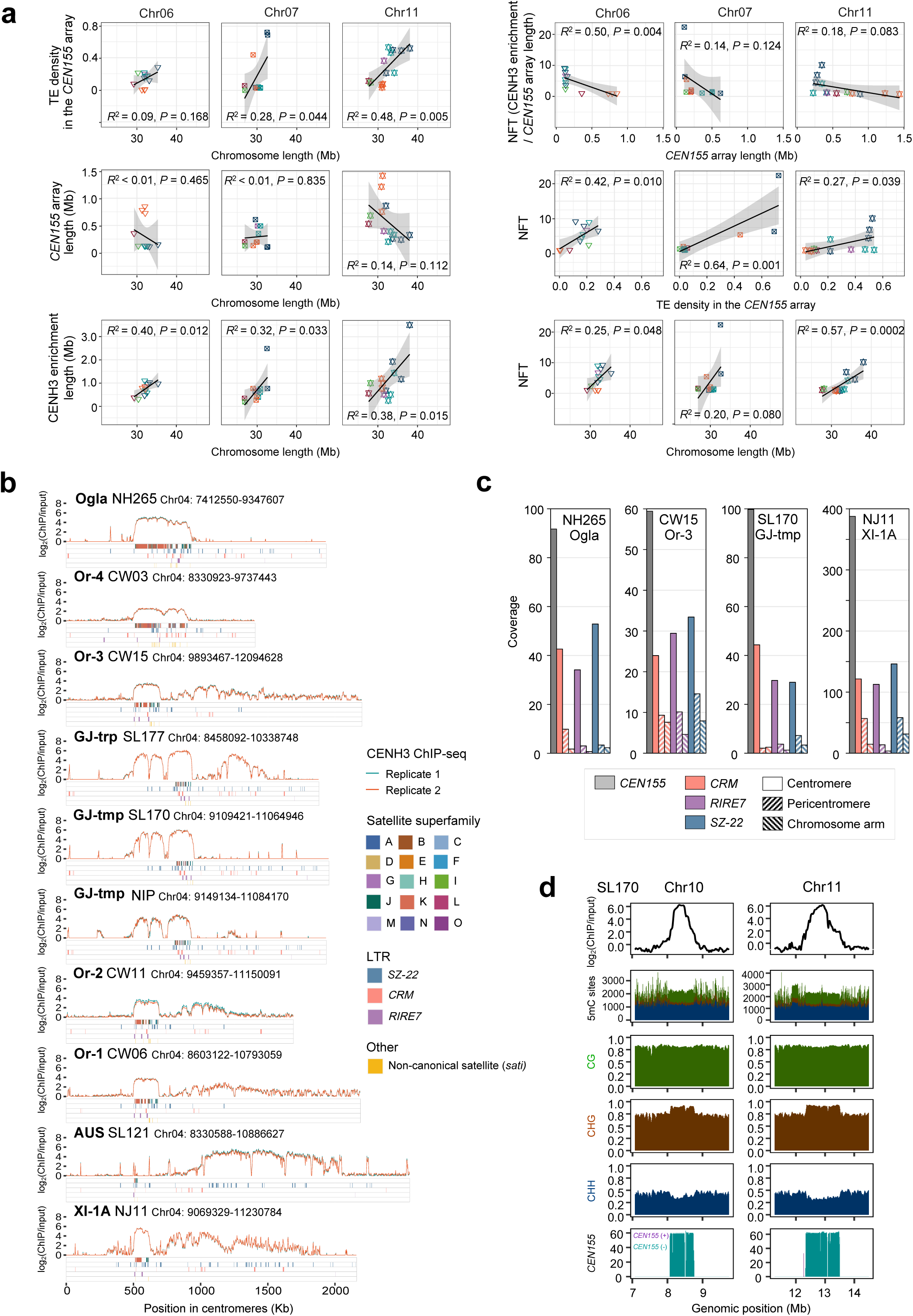
Epigenetic profiling and centromere repositioning. **a**, Correlation analysis showing relationships among variations in chromosome length, *CEN155* array length, CENH3 occupancy signal, TE density and neocentromere formation tendency (NFT). **b**, CENH3 ChIP enrichment and element annotation across Chr04 centromeres. **c**, CENH3 enrichment levels in the *CEN155* satellites, centromeric LTRs, pericentromeric LTRs and LTRs in chromosome arms. **d**, DNA methylation levels in the CG, CHG and CHH sequence contexts within centromeric and flanking regions of chromosomes Chr10 and Chr11 in a GJ accession SL170.

On chromosome Chr04, the CENH3 binding enrichment regions of African rice Ogla (NH265) and Or-4 (CW03) aligned with the *CEN155* arrays (∼400 Kb), indicative of the ancestral state of kinetochore assembly sites (**Fig. 6b**). Conversely, other Chr04 centromeres exhibited significantly shortened *CEN155* arrays (∼131 Kb) yet displayed variable and expanded CENH3 binding regions (∼808 Kb) (**Fig. 6b**; **Supplementary Table 11**). Compared with CW15 (Or-3, ancestral group of GJ), GJ expanded a subregion of CENH3 binding enrichment on the left of its *CEN155* array, accompanied with the contraction of *CEN155* arrays from ∼185 Kb to ∼117Kb. Notably, despite minimal structural variations between the centromeres of GJ-trp (SL177) and GJ-tmp (NIP), GJ-trp (SL177) had a distinct ChIP-seq peak on the right of its *CEN155* array, suggesting either the emergence of a potential neocentromere in GJ-trp or the inactivation of the functional centromere on that side of *CEN155* array in GJ-tmp (**Fig. 6b**). CW11 and CW06, from group Or-1/2 (ancestral wild group of XI and AUS), exhibited discernible CENH3 enrichment within their *CEN155* arrays, with supplementary but robust CENH3 binding signals observed to the right of their *CEN155* arrays in comparison to their progenitor group Or-4 (CW03).

These signals were further amplified in XI (NJ11), implying a functional transition from the *CEN155* arrays to neocentromeres, concomitant with the shortening of *CEN155* array size from ∼172 Kb to ∼136 Kb. Notably, *CEN155* array of AUS (SL121) were drastically degraded (∼25 Kb), with a distinct neocentromere established and functioned on the right side of its *CEN155* array. On chromosome Chr11, distinct neocentromeres were also observed in Or-1/2 (CW11, CW06) and Or-3 (CW15), characterized by markedly shorter CEN155 arrays compared to other groups (**Supplementary Fig. 2**8). These findings collectively suggest a correlation between the emergence of neocentromeres and the demise of native satellite arrays.

Chromosome Chr05 had the lowest density of centrophilic TE insertion among the 12 chromosomes (**Supplementary Fig. 9**). In contrast to the centromeres of outgroup African rice (NH265 and NH284) and Or-4 (CW03), where *CEN155* arrays matched the CENH3 binding regions, all other centromeres showed additional CENH3 binding beyond the *CEN155* arrays (**Extended Data Fig. 6**). Moreover, GJ-trp (SL177) displayed newly evolved CENH3 binding regions compared to GJ-tmp (SL170 and NIP). Interestingly, comparative analysis of centromere structures revealed a specific *SZ-22* LTR insertion within the *CEN155* array of GJ-trp (SL177) (**Extended Data Fig. 6**). Consequently, we propose that TE insertion may trigger and initiate centromere repositioning events leading to neocentromere formation.

The regions enriched with CENH3 occupancy beyond the *CEN155* arrays were predominantly situated near the *CEN155* arrays, centrally distributed within pericentromeric regions, where centromeric LTRs (*SZ-22*, *CRM* and *RIRE7*) were also notably enriched (**Fig. 6b**; **Supplementary Figs. 20-29**). These CENH3 signals showed peaks at the centrophilic LTR loci outside the *CEN155* arrays, particularly in the pericentromeric regions, implying these LTRs may function alternatively as *CEN155* satellites, in spite of relatively lower enrichment intensity compared to *CEN155* satellites and corresponding LTRs within the *CEN155* arrays (**Fig. 6c**). Above evidence implied that the function in kinetochore assembly has driven the evolution of rice centromeric LTRs and centrophilic LTRs appear to mimic the function as centromere satellites as an adaptive strategy to avoid eradication.

Previous studies on human and *Arabidopsis* centromeres have reported a depletion of DNA methylation in centromeres, forming centromere dip regions (CDRs) (Naish *et al*., 2021; Wlodzimierz *et al*., 2023; Logsdon *et al*., 2024). We profiled centromeric DNA methylation using ONT and PacBio HiFi data (**Supplementary Fig. 3**9). Across the entire chromosome, centromere and adjacent pericentromeric regions displayed higher DNA methylation level than the chromosome arms. Within the centromere regions, methylation levels showed no obvious difference among *CEN155* arrays and their flanking regions, with the methylation level of CHH notably lower than CHG and CG methylation (**Fig. 6d**).

Generally, higher CHG methylation, lower CHH methylation and stable CG methylation were observed in the centromere regions compared to the adjacent pericentromeric heterochromatin, particularly in the homeostatic centromeres where the *CEN155* arrays aligned with CENH3 occupancy regions (**Fig. 6d**). The methylation landscape in rice centromeres differs from that in *Arabidopsis* centromeres with a depletion of CHG and CG methylation, possibly associated with the distinctness of satellite sequences and the homeostasis of CENH3 loading (Wlodzimierz *et al*., 2023).

## Discussion

Investigating the mechanisms driving centromere evolution and their impact on genome structure and function is essential for understanding eukaryotic evolution and speciation processes. Based on our comprehensive analyzes, we propose an evolution model of rice centromeres here (termed as <u>R</u>etrotransposon Induced <u>C</u>entromere <u>E</u>volution model, RICE), where genetic and epigenetic dynamics jointly have shaped the rapid evolution of centromeres, emphasizing the importance of LTR insertions in the centromere evolution cycle (**Fig. 7**). Genetic innovation of centromeres in RICE model primarily arise from internal structural variations and the retrotransposon insertion from outside. Via inter-chromosome, individual, group, subspecies and species comparisons in rice centromeres, we have observed diverse structural variations on multiple scales from recurrently local expansions and contractions at the monomer, dimer and multimer levels, to larger-scale centromere splitting and duplication events. Models such as break-induced replication (BIR) model and unequal exchange model provide plausible explanation for the formation of such complex nested structures (Rice, 2019; Talbert & Henikoff, 2022). Despite sharing a common *CEN155* repertoire inherited from a common ancestor, the centromeres in GJ and XI exhibit distinct compositions and structures due to lineage-specific local homogenization progresses, which may have played an important role in the formation of reproduction barriers between them. Contrary to the rapid structural alteration, the single-base substitution rate in centromeres, inferred from absolutely orthologous satellite pairs, appears relatively lower compared to the chromosome arms. Although rice centromere sizes are diverse among individuals, the variation in chromosome length is attributed to TE or LTR insertions. Longer chromosomes typically possess extended functional centromeres, necessary for microtubule attachment during cell division. However, at the same time excessive LTR insertions would invade the *CEN155* arrays, disrupt CENH3 binding continuity of kinetochore assembly and potentially trigger centromere repositioning. These centrophilic LTRs mimic the function of centromere satellites in binding to avoid being purged. In this study, we have observed the emergence of evolutionary neo-centromeres in rice, characterized by CENH3 binding regions that extend significantly beyond the satellite arrays. These neo-centromeres often form in close proximity to the progenitor centromere and are notably enriched with centrophilic LTRs. Unlike the KARMA (kinetochore-associated recombination machine in *Arabidopsis*) mechanism, where LTR insertions primarily contribute to satellite expansion (Wlodzimierz *et al*., 2023), our findings propose a different role for LTRs in plant centromere evolution. Specifically, we suggest that LTR insertions may lead to the decline or loss of progenitor centromeres composed of canonical satellite repeats, thereby facilitating the establishment of neo-centromeres in adjacent regions (**Fig. 7**). For instance, compared to the centromeres of rice AA genomes investigated in this study, the centromeres of rice GG genome (*Oryza granulata*) have no canonical tandem repeats but are primarily occupied by *gypsy*-type LTRs, coresponding to the fact that the rice GG genome has much higher LTR content (Wu *et al*., 2018).

**Figure 7.**
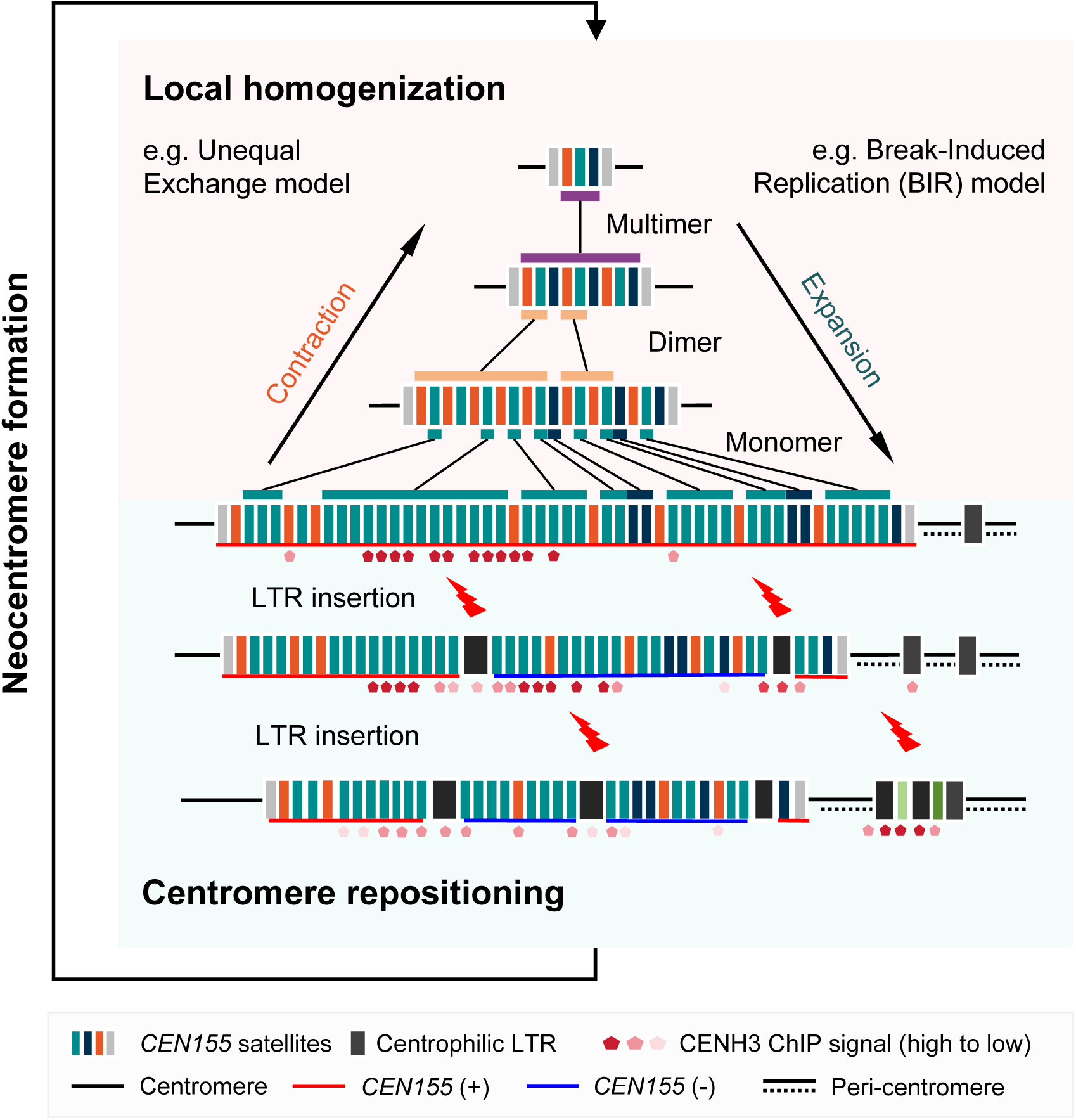
The RICE (Retrotransposon Induced Centromere Evolution) model.

Besides the rice genomes, the proposed model parallels observations in other genomes such as soybean and maize, where high LTR retrotransposon contents have been associated with frequent centromere repositioning (Liu *et al*., 2023; Chen *et al*., 2023). Particularly, wheat centromeres have no canonical satellites but almost are filled by TEs, in line with their extremely high LTR genomic contents (Ahmed *et al*., 2023). In contrast, in the human and chicken genomes, CENH3 or CENP-A binding regions are typically embedded within arrays of higher-order repeats (HORs) at centromeres (Altemose *et al*., 2022; Huang *et al*., 2023). Similarly, in *Arabidopsis*, CENH3 peaks coincide with the presence of satellite *CEN178* or within *CEN178* regions on specific chromosomes (Naish *et al*., 2021; Wlodzimierz *et al*., 2023). Based on these comparative insights, we hypothesize that genomes with higher LTR contents may be more prone to evolving neo-centromeres through mechanisms involving the disruption or replacement of existing centromeric structures. The TE expansion and polyploidization are pivotal adaptive strategies for plants to cope with rapid environmental changes. As a by-product of TE activity to maintain fitness, the rapid centromere alteration may have facilitated the formation of reproduction barrier and speciation to a certain extent, which suggest a novel insight into the evolutionary trade-off between survival and reproduction. Further investigations into the precise molecular mechanisms underlying centromere reorganization in response to LTR insertions will be essential for validating these hypotheses and expanding our understanding of genome evolution in diverse organisms.

## Methods

### Pacbio HiFi and ONT sequencing

A total of 70 rice accessions from *Oryza* AA genome group were including in this study, comprising 51 Asian rice *O. sativa*, 11 wild *O. rufipogon*, four African rice *O. glaberrima* and four wild *O. barthii* samples (**Supplementary Table 1**). The rice plants were cultivated at the China National Rice Research Institute (CNRRI), Hangzhou, China. For Pacbio HiFi sequencing, we newly generated Pacbio HiFi data for 46 accessions, and the HiFi reads for 24 accessions were adopted from previous studies (Wu *et al*., 2023; Shang *et al*., 2023; Zhang *et al*., 2022; Sedeek *et al*., 2023). High-molecular-weight DNA from a single individual was extracted from young seedlings for each accession using the CTAB method. DNA degradation and contamination was scanned on 1% agarose gels. DNA purity and concentration were quantified using a NanoDrop One UV-Vis spectrophotometer (Thermo Fisher Scientific, USA) and a Qubit 4.0 Fluorometer (Invitrogen, USA). According to the methods in the SMRTbell prep kit 3.0 kit manual, HiFi libraries for sequencing were prepared, including DNA quality control, DNA shearing and cleanup, DNA repair and A-tailing reaction, adapter ligation, nuclease treatment and library size selection. Sequencing was conducted on the PacBio Revio platform following the manufacturer’s instructions..PacBio HiFi reads with quality scores over Q20 were generated from raw sequencing reads using PacBio Circular Consensus Sequencing tool ccs (min passes = 3, min RQ = 0.99). The sequencing depth ranged from 23.8× to 49.5× across genomes with an average of 30.4× (**Supplementary Table 1**). For ONT sequencing of ten samples, genomic DNA was extracted using a QIAGEN Genomic DNA extraction kit (Cat #13323, QIAGEN). DNA purity and concentration were determined using a NanoDrop One UV-Vis spectrophotometer (Thermo Fisher Scientific, USA)and a Qubit 3.0 Fluorometer (Invitrogen, USA), respectively. Long DNA fragments were selected using the PippinHT system (Sage Science, USA) via gel cutting. Subsequently, DNA repair and adapter ligation were performed using an SQK-LSK110 kit. The prepared DNA library was then loaded onto a PromethION flow cell (R10.4.1). Base modification information was called using Guppy (v6.4.6). Reads with quality scores less than seven were discarded. For 50 samples, ONT reads were from previous studies (Shang *et al*., 2022; Zhang *et al*., 2022) and downloaded from NCBI Sequence Read Archive (BioProjects PRJNA656318 and PRJNA692836) and National Genomics Data Center (BioProject PRJCA008812). Additionally, available whole-genome NGS data for 64 accessions were adopted from previous studies (Shang *et al*., 2022; Shang *et al*., 2023) (**Supplementary Table 1**).

### CENH3 Chromatin immunoprecipitation and sequencing (ChIP-seq)

ChIP experiments for ten samples were performed following standard protocols (Nagaki *et al*., 2003). Approximately 5 g of 4-week-old seedlings were well grounded in liquid nitrogen for 1 hour and then transferred into 50 ml Eppendorf tube with 20 ml nuclei isolation buffer (10 mM Tris-HCl pH 7.5, 3 mM CaCl_2_, 2 mM MgCl_2_, 0.1 mM PMSF, 1 EDTA-free protease inhibitor cocktail tablets (Roche) and 0.5% Tween 40). After incubating on ice for 15 min, the mixture was filtered through one layer of Miracloth (Calbiochem 475855). The liquid was centrifuged at 600 rcf for 10 min at 4D. Discard the supernatant and resuspend the nuclei pellet with 5 ml 1 × TBS buffer (10 mM Tris-HCl pH 7.5, 3 mM CaCl_2_, 2 mM MgCl_2_, 0.1 mM PMSF, 1 EDTA-free protease inhibitor cocktail tablets (Roche)) with 25% sucrose, and then added on top of 1 × TBS buffer with 50% sucrose gently and centrifuged at 1,500 rcf for 10 min at 4D. Collect the nuclei and digest the chromatin into nucleosome by MNase in MNB buffer (10% sucrose, 50 mM Tris-HCl pH 7.5, 4 mM MgCl_2_, 1 mM CaCl_2_, 0.1 mM PMSF) for 6 min at 37D. The nucleosomes were incubated with pre-immune rabbit serum or CENH3 antibody at 4D overnight, and was then incubated with rProtein A SepharoseTM Fast Flow (GE Healthcare) for 2 hours at 4D. Preimmune rabbit serum was used as the negative control. The rProtein A Sepharose was collected by centrifugation at 13,000 rpm for 1 min at 4D, and washed once using Buffer A (50 mM Tris-HCl pH 7.5, 10 mM EDTA, 0.2 mM PMSF) with 50 mM, 100 mM, and 150 mM NaCl. Immunoprecipitated DNA-protein complexes were eluted twice from the beads with elution buffer (1% SDS, 50 mM NaCl, 20 mM Tris-HCl pH 7.5, 5 mM EDTA) at 65D for 15 min. The DNA was extracted with equal volume of phenol:chloroform (1:1) and the aqueous layer was transferred to new tube. The aqueous layer was then precipitated with isopropanol, sodium acetate, and glycogen at -20D for 2 hours, followed by washing with 70% alcohol once and air-drying. The DNA extraction, precipitation, and washing were performed by centrifugation at 12,000 rpm for 10 min at room temperature, then resolved with TE buffer and stored at -20 D. The anti-CENH3 ChIP-seq library was constructed using rabbit polyclonal anti-CENH3 against the rice-specific peptide. ChIP samples were PCR amplified for 12 cycles and sequenced (2×150bp) on the MGISEQ-200 platform.

### Genome assembly

Among the 70 assemblies analyzed, 67 were newly assembled for this study. Three previously released assemblies (NIP, MH63 and ZS97), assembled completely using extremely high sequencing depth of long reads, were downloaded from the Rice Information GateWay (RIGW) and Rice Super Pan-genome Information Resource Database (Shang *et al*., 2023; Song *et al*., 2021). Newly generated PacBio HiFi reads with adaptor sequences were filtered using HiFiAdapterFilt (Sim *et al*., 2022). For 50 accessions with both PacBio HiFi and ONT reads, we integrated HiFi and Nanopore data together to generate draft *de novo* assemblies using Hifiasm (0.19.5-r587)(Cheng *et al*., 2024) and Verkko (v1.4)(Rautiainen *et al*., 2023), respectively. For the remaining 17 accessions with only PacBio HiFi reads available, we used the HiFi-only mode of Hifiasm and Verkko for assembly, respectively. The scaffolded assembly of Verkko was split into contigs and used to scaffold and extend Hifiasm contigs using a in-house script. The extended contigs (larger than 10 Kb) were then scaffolded on chromosomes using RagTag with no further chimeric splitting (v2.1.0, scaffold -f 2000 --remove-small --aligner minimap2)(Alonge *et al*., 2022), using T2T assembly of NIP as the reference genome (Shang *et al*., 2023). To correct base-level assembly errors and mitigate potential PacBio HiFi sequencing homopolymer errors, we polished the gap-filled assemblies using both HiFi and NGS reads. High-frequency *k*-mers (top 0.02%) were generated using meryl (v1.4)(Rhie *et al*., 2020) and Winnowmap2 (v2.03)(Jain *et al*., 2022) aligned HiFi reads against the assemblies. NextPolish2 (Hu *et al*., 2024) was then employed for polishing the assembly with default parameters. Two *k*-mer datasets (21-mer and 31-mer) were prepared using yak (https://github.com/lh3/yak). For five red rice accessions (Sedeek *et al*., 2023) and one weedy accession YCW03 (Wu *et al*. 2023), assemblies were not polished using short reads, due to their unavailability. To prevent potential over-scaffolding based on homology, we detected and trimmed the potential mis-joins at both ends of each chromosome. Telomere regions were identified by searching the rice telomeric repeat motif TTTAGGG with the occurrence of at least five consecutive repeats. Non-telomeric sequences outside the telomeric region (< 2 Mb) were then pruned from the assemblies.

### Assembly quality assessment

We first manually checked and corrected the potential large structural errors in assembly by aligning to the NIP reference. Assembly quality for each genome was evaluated using a series of indices from completeness, correctness and continuity. For completeness, NGS short reads and long reads (PacBio HiFi and ONT) were aligned to their corresponding chromosomes for each sample with BWA mem (0.7.17-r1188)(Li & Durbin, 2009) and Winnowmap2 (v2.03)(Jain *et al*., 2022) in default parameters, respectively. The mapping statistics was generated by using samtools flagstat (v1.18-15-g9a59467) (Li *et al*., 2009). Averagely 97.62% of short reads were aligned, while wild rice *O. rufipogon* had significantly lower mapping rates (∼95.23%) compared to *O. sativa* (∼98.35%), due to the nature of higher heterozygosity in wild population (**Supplementary Table 1**). ∼99.92% and ∼98.71 of PacBio HiFi and ONT reads were mapped. The mapping coverage along each chromosome was calculated using samtools depth and the coverage distribution was checked visually to detect assembly collapsed regions in 10-Kb windows. For correctness, we used QV and discordant *k*-mers in VerityMap to assess the base accuracy. NGS short reads and Pacbio HiFi reads were split into 21-mers and combined as a *k*-mer library using meryl (v1.4) (Rhie *et al*., 2020) and rare *k*-mers were removed (meryl gt 1). Assembly correctness indicator QV and assembly completeness indicator *k*-mer completeness were calculated using Merqury (v1.3) (Rhie *et al*., 2020). It should be noted that NGS data used here was sequenced after polymerase chain reaction (PCR) amplification, thus we analyzed the potential effects of PCR on QV. We sequenced PCR-free NGS libraries for four samples (CW06, CW15, SL044 and SL177) using Illumina Novaseq 6000. Using only NGS *k*-mer library to calculate QV, PCR NGS data yielded a lower QV than PCR-free data, but varied among samples (46.4 for PCR-free QV *versus* 25.8 for PCR QV in CW06, 47.1 *versus* 23.2 in CW15, 47.4 *versus* 40.1 in SL044 and 43.5 *versus* 44.2 in SL177). We also employed VerityMap (Bzikadze *et al*., 2022) to detect assembly errors and heterozygosity sites in a reference-free mode. VerityMap identifies discordant *k*-mers between the assemblies and PacBio HiFi reads. High frequency of discordant *k*-mers indicate potential errors in local assembly. For continuity, we detected potential assembly gaps with low-confidence read supports using CRAQ (Li *et al*., 2023) and GCI (Chen *et al*., 2024)(**Supplementary Fig. 1**). We aligned the PacBio HiFi reads and NGS reads against corresponding assemblies and calculated AQI values to identify regional and structural assembly errors based on clipped alignment information. GCI is a continuity inspector for complete genome assembly, by utilizing stringently filtered alignments from multiple aligners (minimap2 and Winnowmap2 used in this study) to detect assembly gaps. We quantified the assembly overall continuity using GCI scores at the whole-genome and chromosome levels.

### Evaluation of centromere assembly completeness

To assess the assembly quality of rice centromere sequences, we first determined the centromere regions based on the enrichment of rice *CEN155* satellites. The quality assessment metrics (e.g. QV. GCI) were calculated for each centromere (**Supplementary Fig. 2**). We zoomed in and manually checked the raw and curated reads mapping coverage and the issues reported by GCI and VerityMap in the centromere regions. Additionally, given the challenges in assembling highly repetitive sequences accurately and completely, we compared the centromere sequences solely generated by Hifiasm (Cheng *et al*., 2024) and Verkko (Rautiainen *et al*., 2023) to eliminate the effects of assembly algorithms. For each accession, two sets of contig assemblies from Hifiasm and Verkko with no gap filling and polishing were anchored onto chromosomes using RagTag, respectively. The *CEN155* consensus sequence was then aligned to each chromosome using BLASTN (v2.5.0+, -use_index true -task megablast -evalue 1e-6). We counted the copy numbers of *CEN155* satellites in each centromere and compared the centromeres with no gaps in both Hifiasm and Verkko assemblies (**Supplementary Fig. 2**).

### Phylogeny and ancestry analysis

Short Illumina paired-end reads from each sample were aligned to the T2T NIP reference assembly (Shang *et al*., 2023) using BWA (Li & Durbin, 2009). For six samples lacking NGS short reads (YCW03, Pulut Hitam-2, Zag, Cempo Ireng, Balatinaw, and Cempo Abang), simulated 120-bp reads were generated from their genome assemblies and mapped against the T2T NIP assembly. Whole-genome SNPs across all 70 samples were jointly called following the GATK best practices (McKenna *et al*., 2010). A phylogenetic tree encompassing these 70 rice accessions was constructed using FastTreeMP (Price *et al*., 2009) with 1000 bootstrap replicates. This tree was utilized to elucidate the taxonomic relationships and groupings among the rice accessions.

### Gene annotation

Gene elements were annotated using a homology-based approach. We employed Gmap to align transcripts from the MSU v7.0 annotation of Nipponbare with the parameters ’-n 0 --min-trimmed-coverage 0.7 --min-identity 0.7’, and kept alignments with at least 70% coverage and 90% identity (Wu & Watanabe, 2005). Cleaned RNA-seq reads were mapped to the corresponding genome assembly using TopHat2 (Kim *et al*., 2013), and gene expression was quantified using Cufflinks (Trapnell *et al*., 2010).

### TE annotation

To characterize repeat elements across the 70 high-quality rice genomes, a hybrid approach combining *de novo* prediction and homology-based search strategies was employed. Initially, a *de novo* repeat database was constructed using assemblies from 16 representative rice accessions (including 18XHB-57, 18XHB-83, CW03, CW06, CW11, CW15, MH63, NH271, NH285, NIP, SL028, SL039, SL044, SL121, YZ-2 and ZS97). The *de novo* transposable element (TE) library was generated separately for each accession using RepeatModeler with the parameter ’-LTRStruct’ (Tarailo-Graovac & Chen, 2009). These individual TE libraries were integrated into a raw library consisting of 37,589 records, and subsequently clustered and merged into a non-redundant library. To remove redundant records, pairwise alignment of TEs from each accession and TE library of rice in repbase (Repbase 28.09) using BLAST were performed. Records showing alignment lengths over 60% of the element’s length and identities over 60% were considered as redundant records. A total of 23,541 redundant records were eliminated from the final library. For further filtering, specific accession de novo TEs library and that from Repbase was used to mask 16 representative accessions by RepeatMasker separately (Tarailo-Graovac *et al*., 2009).

TEs, that did not overlap with records in other libraries, had an accumulated length over 5 Kb and more than 10 instances in presence, were selected as candidate records. Subsequently, 341 non-redundant TEs identified in 2 to 15 accessions were clustered based on similarity using CD-HIT (Li & Godzik, 2006). The final non-redundant library comprised 3,847 sequences, including 3,321 records from the rice TE library (Repbase 28.09), 341 homologous TEs and 185 accession-specific TEs. All genome assemblies were masked against the newly-built non-redundant TE library using RepeatMasker. For elements classified as ’UNKNOWN’ by RepeatModeler in more than 50% of cases, a convolutional neural network-based approach (deepTE) was applied for reclassification (Yan *et al*., 2020). Full-length elements were defined based on the annotation results, initially merged based on identity and alignment length to obtain non-redundant or overlapping entries. Specifically, full-length LTR retrotransposons were characterized by the canonical LTR-Internal-LTR structure, with the intervals between LTR and internal units being less than 10% of the consensus length, and the overall length deviating by no more than 10% from the consensus. To ensure accuracy, candidate full-length LTRs meeting these criteria were extracted and compared against consensus sequences, applying a minimum coverage threshold of 90%. Solo-LTRs were defined as units lacking TE annotations of the same family within a distance of 10% of the consensus length.

### Centromere annotation

The Tandem Repeat Annotation and Structural Hierarchy (TRASH) pipeline (Wlodzimierz *et al*., 2023), available at https://github.com/vlothec/TRASH, was utilized for the prediction of centromeric satellite repeats across genomes. The pipeline was configured with parameters "--simpleplot --horclass *CEN155* --frep 2 --par 5 --k 1000". Genomic sequences were segmented into 1-Kb windows to assess local *k*-mer counts and identify repetitive regions. Score of each window was determined based on the proportion of repeated *k*-mers, with regions scoring above a defined threshold considered to contain repeats. Tandem repeats within these windows were characterized by the distances between identical *k*-mers, aiding in the identification of consensus sequences such as *CEN155*. To pinpoint centromeric regions within each rice chromosome, all satellite repeats identified as *CEN155* from the TRASH results were extracted and mapped according to their chromosomal positions. *CEN155* arrays, indicative of centromeric blocks, were defined where the spacing between repeat units did not exceed 150 Kb. The largest contiguous block on each chromosome was considered as the candidate centromeric region. Subsequently, the *CEN155* consensus sequence for each rice accession was aligned back to its respective genome assembly using BLASTN (v2.5.0+) under default settings. These alignments were integrated into the centromeric blocks based on their proximity to adjacent *CEN155* arrays. Final validation of centromeric regions involved manual inspection, confirming regions supported by both TRASH predictions and BLASTN alignments as definitive centromere locations. Synteny analysis was performed to validate the predicted centromere boundaries across 69 rice accessions by extracting and examining the 800-kb flanking regions surrounding the centromere in the NIP reference genome for collinearity with other rice centromere assemblies. Sequence identity heat maps were plotted for each chromosome using StainedGlass (Vollger *et al*., 2022), providing a visualization strategy to confirm centromere boundaries.

### Centromere similarity and haplotyping

Centromere classification was initiated based on pairwise similarity. Firstly, pairwise alignments were performed between all centromeres divided into 2-Kb windows using minimap2 (v.2.24) with parameters (-f 5000 -s 400 -ax ava-ont --dual=yes --eqx) (Li, 2018). Segment windows with identities over 95% were merged, and the cumulative length was designated as the identical length between centromeres. The cumulative length between different centromeres was normalized by the length of the query centromere, highlighting structural variations such as deletions causing asymmetry between the upper and lower triangles in the heat map (**Fig. 2**). Next, specific haplotypes were assigned to each centromere. Haplotypes were manually defined based on identity heat maps generated from 2-Kb windows of different centromeres. Candidate reference haplotypes were established, comprising sequences from XI and GJ that shared at least 95% of full length with two or more accessions. The longest regions among candidate centromeres for each haplotype were selected as reference haplotype sequences. Subsequently, windows split into 2-Kb segments from each centromere were classified into corresponding haplotypes based on the identity among all haplotypes, when the alignment covered at least 85% of the window size. In cases where multiple haplotypes exhibit the same identity, the window was assigned to the haplotype with the fewest mismatches against reference haplotype sequences.

### Structural variations and SNPs around centromeres

HiFi reads from each accession were aligned to the reference assembly of NIP using minimap2 (Li, 2018) with parameters (-ax map-hifi -R "@RG\tID:${pre}\tLB\tSM:${pre}") and SVision (v.1.4) (--min_mapq 10 --min_sv_size 50) (Lin *et al*., 2022) was utilized to generate structural variation (SV) calls. SVs within a 500-Kb upstream and downstream region of each centromere were extracted for statistical analysis. Phylogenetic trees of all accessions were constructed based on SNPs located in the 500-Kb upstream and downstream regions of each centromere with FasttreeMP (Price *et al*., 2009).

### Satellite clustering

The *CEN155* repeats identified by TRASH were classified as canonical satellites if their lengths ranged from 140 bp to 170 bp. Across the 70 rice genomes, a total of 249,952 unique *CEN155* sequences were detected. Initially, satellites occurring frequently (>30 times) across all accessions were selected.

Additionally, satellites predominantly (>90%) found in specific populations (GJ, XI, AUS, Oruf, Obar, and Ogla) were included, despite lower frequencies ranged from 5 to 30. A collection of 6,665 representative *CEN155* repeats were finally obtained. These selected satellites were aligned using MUSCLE (Edgar, 2022) with default parameters, followed by phylogenetic categorization into 15 subfamilies using FastTreeMP (Price *et al*., 2009). To further classify the remaining 243,287 unique satellites, Levenshtein edit distances were calculated in an all-vs-all comparison against the 6,665 categorized satellites. Each unique satellite was assigned to the superfamily of the assigned *CEN155* with the closest edit distance.

### CENP-B box and pJ**_α_** motif analysis

To investigate the DNA binding sites of rice centromeres, we introduced a scoring matrix based on the CENP-B box sequence (5’-PyTTCGTTGGAAPuCGGGA-3’) and the pJα motif (5’-TTCCTTTTPyCACCPuTAG-3’), originally characterized in the human centromeres. Each *k*-mer (*k* = 17) from the query sequences underwent global alignment scoring using a dynamic programming algorithm against the two binding motif sequences. Additional penalties and bonuses were applied at conserved loci. All match scores of all monomer in NIP were used to explore the criteria for defining B-box-like elements. The screening score was set to be top 1% of all scores with a threshold set at 24, denoting the match of all conserved sites. According to the score distribution, the *k*-mers with scores above 25/27 were considered as candidate B-box/pJα-like elements. To enable rapid scanning of genome-wide elements across multiple genomes, the *k*-mers harboring B-box-like elements from NIP genome were integrated using MEME v5.4.1 with parameters ’-w 17 -dna -revcomp’ to construct a B-box-like motif library (Bailey *et al*., 2015). Subsequently, B-box-like elements were identified across the whole genome with *e*-value < 1e-05 against the motif library for B-box and pJα motif separately, utilizing FIMO in MEME with the parameters ’--max-strand’.

### Structural organization analysis of *CEN155* arrays

We resolved the structure of satellite arrays in a progressive compression strategy, merging identical units iteratively (**Fig. 4a**). Homogenization regions of monomer (moHRs) were identified where more than five consecutive satellites from the same family were present. TEs embedded in the satellite arrays were masked. Fine analysis of satellite organization within each moHR was performed using identity dot plots and NTRprism. Pairs of identical satellites within moHRs were selected to investigate internally local collinearity using DAGchainer (Haas *et al*., 2004). NTRprism (Altemose *et al*., 2022) was employed to infer the most abundant tandem repeat periodicities by counting the interval lengths between adjacent instances of each 6-mer. For a canonical HOR array, a distinct peak in the interval spectra was expected at the HOR unit length (Altemose *et al*., 2022). Using the moHR-compressed satellite array, we defined homogenization regions of dimer (diHRs) containing at least five replicates of dimers. For determining homogenization regions of multimer (muHRs), a de Bruijn graph was initially constructed based on the dimer-compressed satellite string of each centromere, where the vertex set comprises all monomer superfamilies and the edge set consists of all pairs of its consecutive monomers superfamily. For instance, for a dimer-compressed centromere sequence “ABCDABCD”, the edge set would include the pairs A-B{2}, B-C{2}, C-D{2}, D-A{1}. Candidate multimers were identified by exhaustively enumerating all cycles in the graph using a depth-first search (DFS) strategy. We observed that ancient multimers might not exhibit a unified head-to-tail pattern like in the human centromeres, due to local expansion, random insertion or deletion. Candidate multimer calls were filtered out based on criteria of multiplicity less than 2, traversal less than 3 and adjacent multimer spacing greater than three times the multimer length. The multiplicity of an edge (A, B) in the graph was defined as the number of times the monomers family A followed the monomer family B in each dimer-compressed centromere. Traversal represented the occurrence frequency of a cycle (multimer) in each dimer-compressed centromere string. To identify multimer hierarchically, candidate nested-multimers were mapped onto the discrete dimer-compressed centromere and detect overlapping regions with continuity and occurrence. If the merged multimer presented better continuity and higher occurrence, it would replace the nested one.

### Centrophilic LTR analysis

To identify centrophilic LTRs, which preferentially localize within centromeres, we conducted a statistical analysis based on the annotation of full-length LTRs. LTRs that constituted more than 90% of all full-length LTRs in centromeres and exhibited a significantly higher preference for the centromere region compared to the chromosome arms were designated as centrophilic LTRs. To characterize the core domains of these LTRs, ORFs within the internal domain of consensus elements from the pan-TE database were extracted using getorf in EMBOSS (Rice *et al*., 2000). Subsequently, domain predictions were performed using HMMER (v3.4) (-E 1e-5) against Pfam hidden Markov models, including PF00026, PF13650, PF08284, PF13975, PF00077, PF09668 for protease; PF03732 for GAG; PF00665, PF13683, PF17921, PF02022, PF09337, PF00552 for integrase; PF17917, PF17919, PF13456 for RNase H; and PF00078 for reverse transcriptase (Mistry *et al*., 2021). Given the presence of non-autonomous LTRs in the pan-TE library, the phylogenetic relationships of LTRs were resolved separately based on the 5’ LTR, 3’ LTR, and internal sequences. Multiple sequence alignments were conducted using MAFFT (--anysymbol –maxiterate 1000), and conserved sites were extracted using trimal (Katoh *et al*., 2013; Capella-Gutiérrez *et al*., 2009). Phylogenetic trees were constructed using FastTreeMP (-nt -gtr) and visualized with iTOL (Price *et al*., 2009; Letunic & Bork, 2021). Pairwise alignments among full-length centrophilic LTRs were performed and visualized using VISTA (Mayor *et al*., 2000). Identity between 5’ LTR and 3’ LTR of centrophilic LTRs was calculated based on full-length LTRs existing within centromeres. Alignments for each LTR pair were conducted using MAFFT, and identities were computed using ape (Paradis & Schliep, 2019). To map the insertion sites of SZ-22 elements, 150-bp regions flanking each element were extracted and queried against the *CEN155* database using BLASTn. Centromeric elements were filtered based on close alignments to the same monomer (2,258 out of 3,673 sequences). The midpoint between aligned flanking sequences determined the insertion positions, and their distribution along the *CEN155* consensus sequence was visualized.

### Pairwise comparison of rice centromeres

We introduced a new algorithm termed SynPan-CEN, to compared satellite arrays in single monomer scale. This approach leverages satellite clustering information and pairwise genetic distances to identify syntenic pairs of *CEN155* satellites between rice centromeres. Initially, we conducted an exhaustive all-to-all comparison of monomer sequences from two satellite arrays using pairwise edit distance (ED) to quantify similarity. We refined synteny construction by conservatively selecting monomer pairs in a correct order, incrementally increasing the ED threshold until the framework encompassed more than 20% of total monomers or 100 monomers. Using DAGchainer software (Haas *et al*., 2004), which computes chains of syntenic genes based on sequence homology and genomic coordinates, we identified syntenic satellite regions and syntelogs. DAGchainer employs a scoring function considering the distance between neighboring elements on each DNA molecule and BLAST E-value scores between homologs, facilitating the identification of maximally scoring chains of ordered element pairs. For the analysis here, DAGchainer parameters (-g 1 -D 20 -A 6 -Z 50 -e -3f) were set to retain blocks with the highest scores, with non-overlapping monomer pairs added to complement the chain backbone. Using a window of ten syntenic pairs, we applied sliding windows along the fixed chain backbone to extract each region. Local syntenic pairs were identified using DAGchainer, maintaining consistency with the parameters mentioned above. Blocks with the highest scores were preserved, and additional non-overlapping syntenic monomer pairs were incorporated. To refine our results, we incrementally tested various ED thresholds and observed a significant impact on identifying optimal syntenic pairs of *CEN155* satellites. Syntenic satellite pairs increased as ED values were below 15, but decreased beyond this threshold. This phenomenon is likely due to the rapid accumulation of potential syntenic paths after exceeding the ED threshold, which complicates the determination of the optimal paths by DAGchainer (Haas *et al*., 2004). In cases where sliding windows presented excessive alignment possibilities, we subdivided windows into two sub-regions, each containing five syntenic pairs, mitigating potential mismatches.

### Sequence divergence and mutation rate in centromeres

Edit distances between syntenic *CEN155* satellite pairs inferred by SynPan-CEN were calculated. Continuous gaps at the beginning or end of a monomer alignment were systematically removed to minimize potential mismatches. Within alignments, consecutive gaps were treated as a single transition, and insertion counts were used exclusively to quantify edit distances. To optimize the alignment process, we compared three widely-used programs— MUSCLE (Edgar, 2022), MAFFT (Rozewicki *et al*., 2019), and ClustalW (Thompson *et al*., 2002)—using default parameters for pairwise monomer sequence alignments. Based on alignment performance and computational efficiency, MAFFT and ClustalW consistently demonstrated superior reliability compared to MUSCLE. Consequently, MAFFT was selected for following analyses. A phylogenetic tree was constructed using all syntenic satellites between chromosomes, where the synteny information was visualized with links in the circle tree plot.

Orthologous pairs were identified based on their proximity to each other in the tree structure. The satellite pair, where the two *CEN155* were clustered as a monoclade, were confirmed as orthologs, while pairs lacking such proximity were more likely to be paralogs.

### DNA methylation analysis using PacBio HiFi data

For the 46 samples whose HiFi reads were newly generated in this study, 5mC methylation probability was determined by combining kinetics from multiple passes in HiFi sequencing. HiFi reads with methylation MM and ML tags were aligned against the corresponding assembly using pbmm2 (v1.2.0) (https://github.com/PacificBiosciences/pbmm2). We employed pb-CpG-tools (https://github.com/PacificBiosciences/pb-CpG-tools) to call base-level methylation status in the model ‘pileup_calling_model.v1’. Given that HiFi-based 5mC signals were predominantly enriched in the CG islands, we focused exclusively on the methylation signals at CG sites along chromosomes, therefore excluding methylation information for CHH and CHG contexts.

### DNA methylation analysis using ONT data

ONT sequencing data was utilized for methylation analysis. Methylation probabilities in CG, CHH, and CHG contexts were called using the ONT reads basecaller Dorado (v.0.5.3) from the raw pod5 files, with the basecalling model options: ‘--modified-bases-models dna_r10.4.1_e8.2_400bps_hac @v4.3.0_5mC_5hmC@v1 --emit-moves’. The base-called read sequences were aligned against their corresponding genome assemblies. Modkit (v.1.0, https://github.com/nanoporetech/modkit) was utilized to generate summary counts of modified and unmodified bases in bedMethyl files. DNA methylation levels were quantified using ‘modkit pileup’ based on only primary alignments. The consistency of methylation quantification between PacBio HiFi reads and ONT reads was assessed by randomly selecting 10,000 CG sites, using Pearson correlation as the evaluation metric.

### ChIP-seq data alignment and processing

CENH3 ChIP-seq reads from 13 accessions (ten newly generated in this study and three previously released) were used to determine the functional centromere regions. Paired-end raw reads were subjected to adapter trimming and filtering using fastp (v.0.23.4) (Chen *et al*., 2023), with the following options: ‘--cut_front_window_size 4 --cut_front_mean_quality 20 -3 --cut_tail_window_size 4 --cut_tail_mean_quality 20 --cut_right --cut_right_window_size 4 --cut_right_mean_quality 20’. Filtered read sequences were aligned to their corresponding genome assembly using Bowtie2 (v.2.3.5.1) (Langmead *et al*., 2019) with parameters ‘--very-sensitive --no-mixed --no-discordant --maxins 800’. Reads that aligned to multiple locations were filtered using samtools (v.1.7) (Li *et al*., 2009). The enrichment level of CENH3 for each 1-Kb window was calculated using bamCompare from the Deeptools package (v.3.5.4.post1) (Ramírez *et al*., 2016) with the parameters ‘--binSize 1000 --outFileFormat bedgraph --operation log2 -p 5 --extendReads’. Base-level CENH3 enrichment was calculated using bamCompare with the parameters ‘--binSize 1 --numberOfProcessors 40 --operation ratio --outFileFormat bedgraph’. Custom R scripts were used to visualize CENH3 occupancy profiles across the 12 chromosomes and *CEN155* satellite repeat arrays.

## Data availability

Raw PacBio HiFi reads for 46 rice accessions, raw ONT sequencing reads for ten accessions, and CENH3 ChIP-seq NGS reads for ten accessions generated in this study have been deposited in NGDC (https://ngdc.cncb.ac.cn/) under the BioProject accession PRJCA025388, with the GSA numbers of CRA016014, CRA016014 and CRA017653, respectively. The newly-generated genome assemblies in this study are available at Zenodo (https://doi.org/10.5281/zenodo.12770803). The genome assemblies of NIP, MH63 and ZS97 are available at the RiceSuperPIRdb (http://www.ricesuperpir.com/) and RIGW (http://rice.hzau.edu.cn/rice_rs3/). The TE and gene annotation files of 70 rice genomes are deposited at Zenodo (https://doi.org/10.5281/zenodo.12698984). Rice centromere annotation and comparison plots for all accessions and chromosomes are supplemented at Zenodo (https://doi.org/10.5281/zenodo.12702715), including similarity heat-map plots generated by StainedGlass for each centromere, whole-genome synteny to the NIP reference assembly, centromere synteny against NIP and NJ11 assemblies and centromere composition for all chromosomes.

## Code availability

The SynPan-CEN code is available at https://github.com/Darlene1997/SynPan-CEN and the scripts for the progressive compression strategy in deciphering the satellite organization and additional in-house codes associated with this study (including assembly, annotation and visualization) are available at https://github.com/dongyawu/CenTools. The similarity heat-map within each centromere was drawn by StainedGlass (https://github.com/mrvollger/StainedGlass). The visualization of centromere annotation and synteny tracks was performed using ggplot2 (https://github.com/tidyverse/ggplot2) in R (v4.3.1, https://www.r-project.org/).

## Acknowledgements

This work was supported by China National Postdoctoral Program for Innovative Talents (BX20220269), China Postdoctoral Science Foundation (No. 2023M743045), Young Scientists Fund of the National Natural Science Foundation of China (No. 32300490) to D.W.; National Key Research and Development Program of China (No. 2019YFA0903904) to W.H.; Biological Breeding-Major Projects (No. 2023ZD04076), Hainan Province Science and Technology Special Fund (No. ZDYF2022XDNY271) and CIC-MIC to L.F. We thanks to Guojie Zhang (Zhejiang University), Yafei Mao (Shanghai Jiao Tong University) and Kun Wu (Zhejiang University) for constructive suggestions.

## Author contribution

D.W. conceived and initiated this study. L.S. collected the samples. D.W., Q.C. and M.S. performed the sequencing data quality control, centromere assembly and quality evaluation. L.X., D.W. and Y.S. performed the analysis of satellite sequence identification and clustering and satellite array organization.

Y.H. and D.W. performed the annotation and centromeric insertion analysis of TEs. W.H., L.X. and S.B. conducted the CENH3 chromatin immunoprecipitation (ChIP) experiments. L.X., D.W. and S.Z. processed the ChIP-seq data and analyzed the epigenetic profiling of rice centromeres. L.F. and D.W. supervised all analyses. Q.Q., W.J., C.Y., L.S. and X.Z. provided suggestions on analysis, organization and writing. D.W., L.X. and Y.H. wrote the manuscripts with the inputs from all co-authors. All authors discussed the results and commented on the manuscript.

---

**Extended Data Figure 1.**
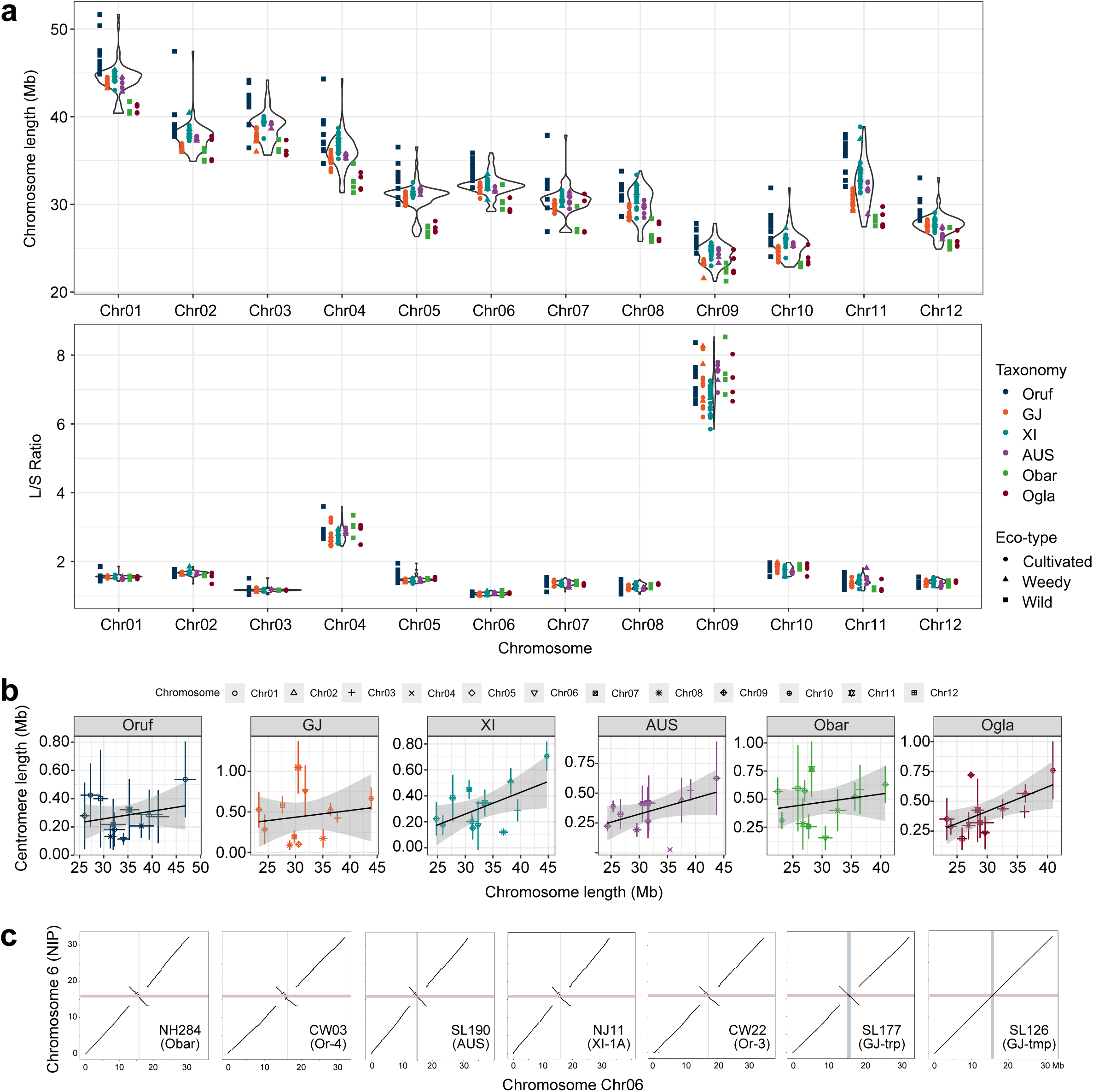
Variation in rice centromere positioning. **a**, Chromosome length and the ratio of long arm length to short arm length across each chromosome. **b**, Relationships between chromosome length and centromere length (defined by the presence of *CEN155* satellites) across different taxonomic groups. **c**, Two megabase-scale inversions observed in the centromere region of chromosome Chr06.

**Extended Data Figure 2.**
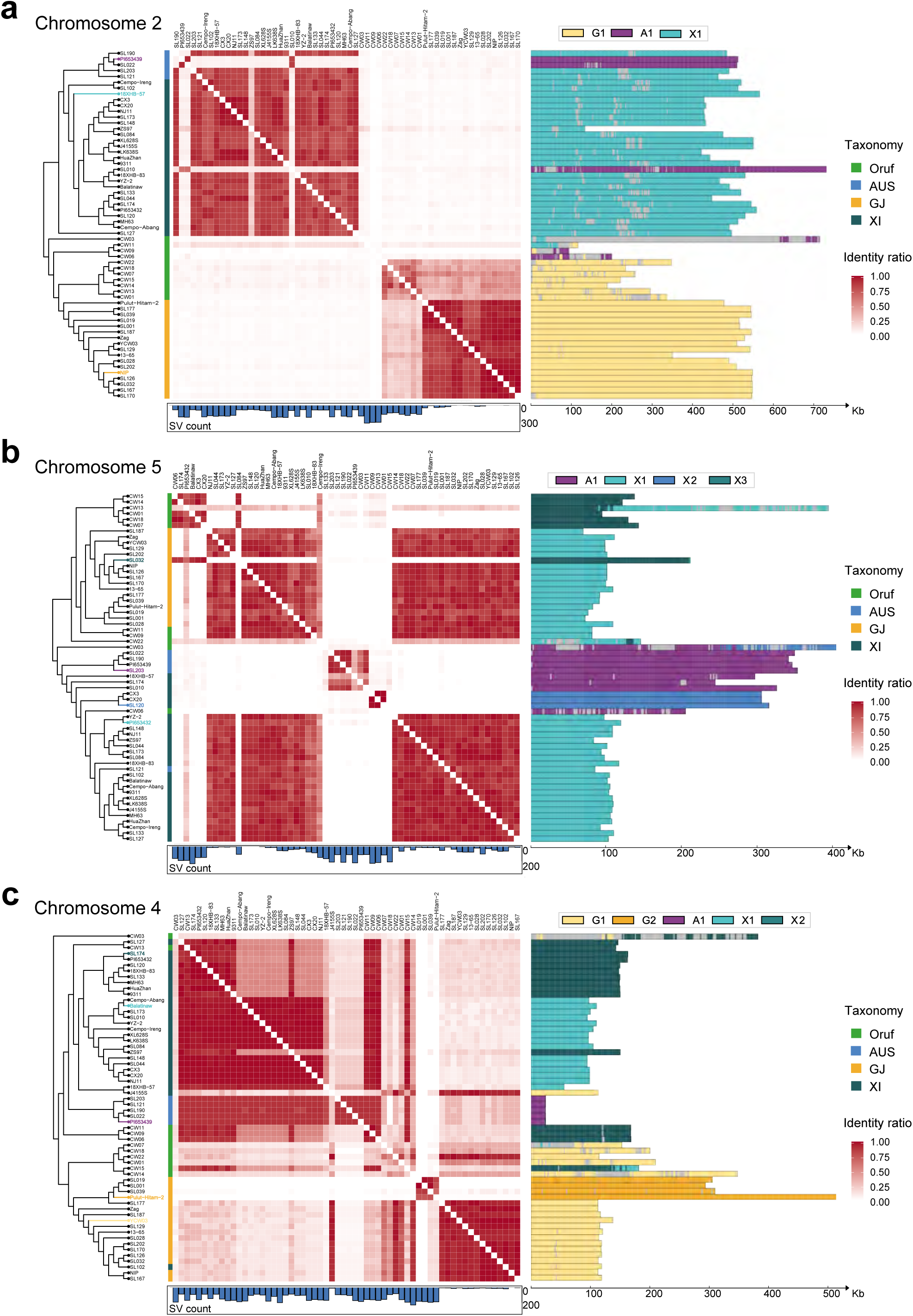
Centromere divergence and introgression in rice evolution. **a**-**c**, Centromere similarity and structural variations compared to NIP on chromosomes Chr02 (**a**), Chr05 (**b**), and Chr04 (**c**).

**Extended Data Figure 3.**
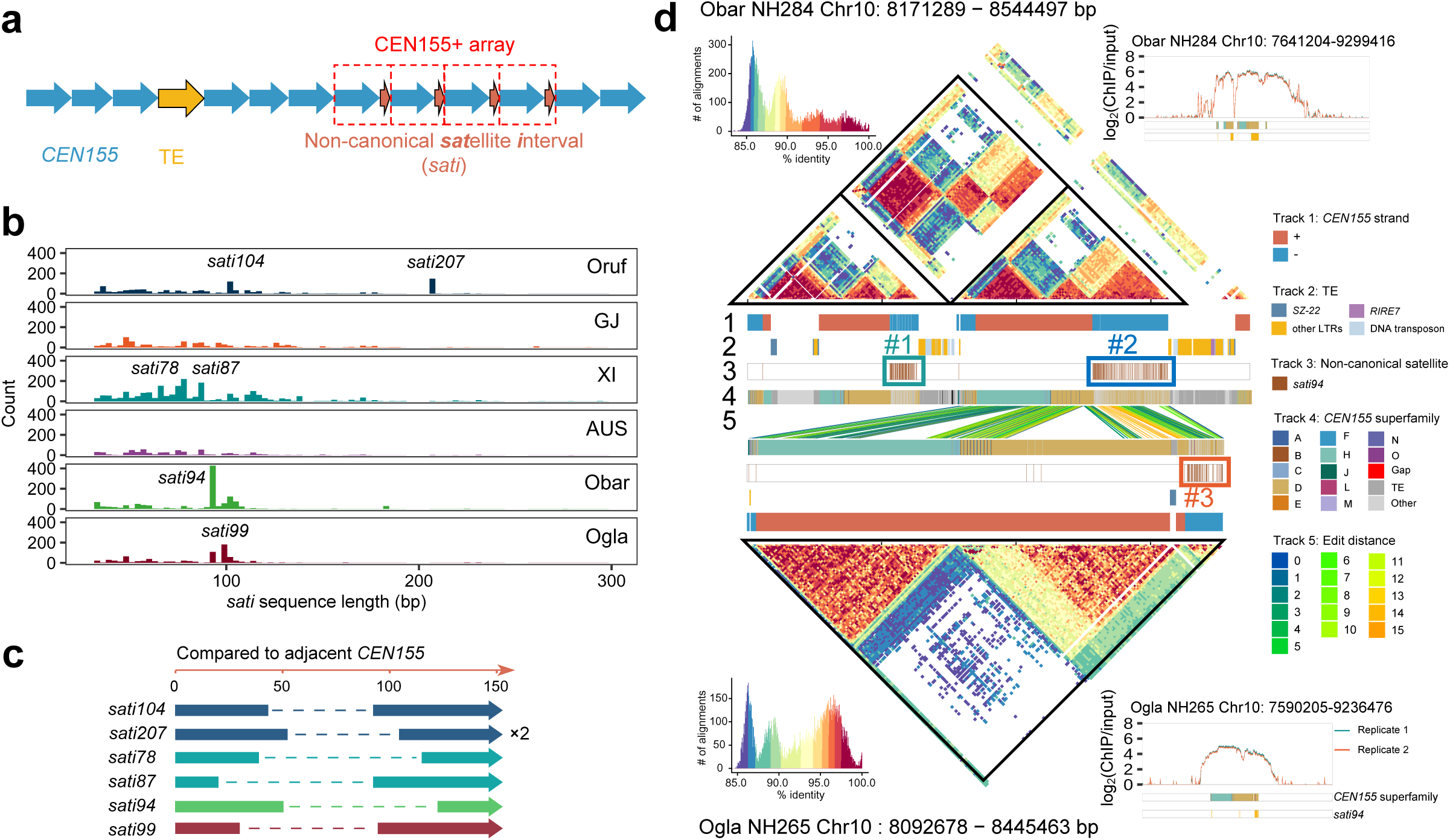
Non-canonical satellites in *CEN155* arrays. **a**, Schematic representation of a centromeric region with non-canonical satellites (*sati*). **b**, Length distribution of *sati* between canonical *CEN155* satellites showing the high-frequency non-canonical satellites. **c**, Origin of *sati* sequences derived from *CEN155* repeats. **d**, Comparison of the two Chr10 centromeres indicating the functional and evolutionary dynamics of CEN155+ arrays incorporating *sati94*. StainedGlass similarity heat maps suggest centromere duplication in NH284 (*O. barthii*), compared to NH265 (*O. glaberrima*), where the CEN155+ arrays composed of *CEN155* and *sati94* are marked from #1 to #3. CENH3 ChIP occupancy plots for the two centromeres indicate the functional consistency of *sati94* as *CEN155*.

**Extended Data Figure 4.**
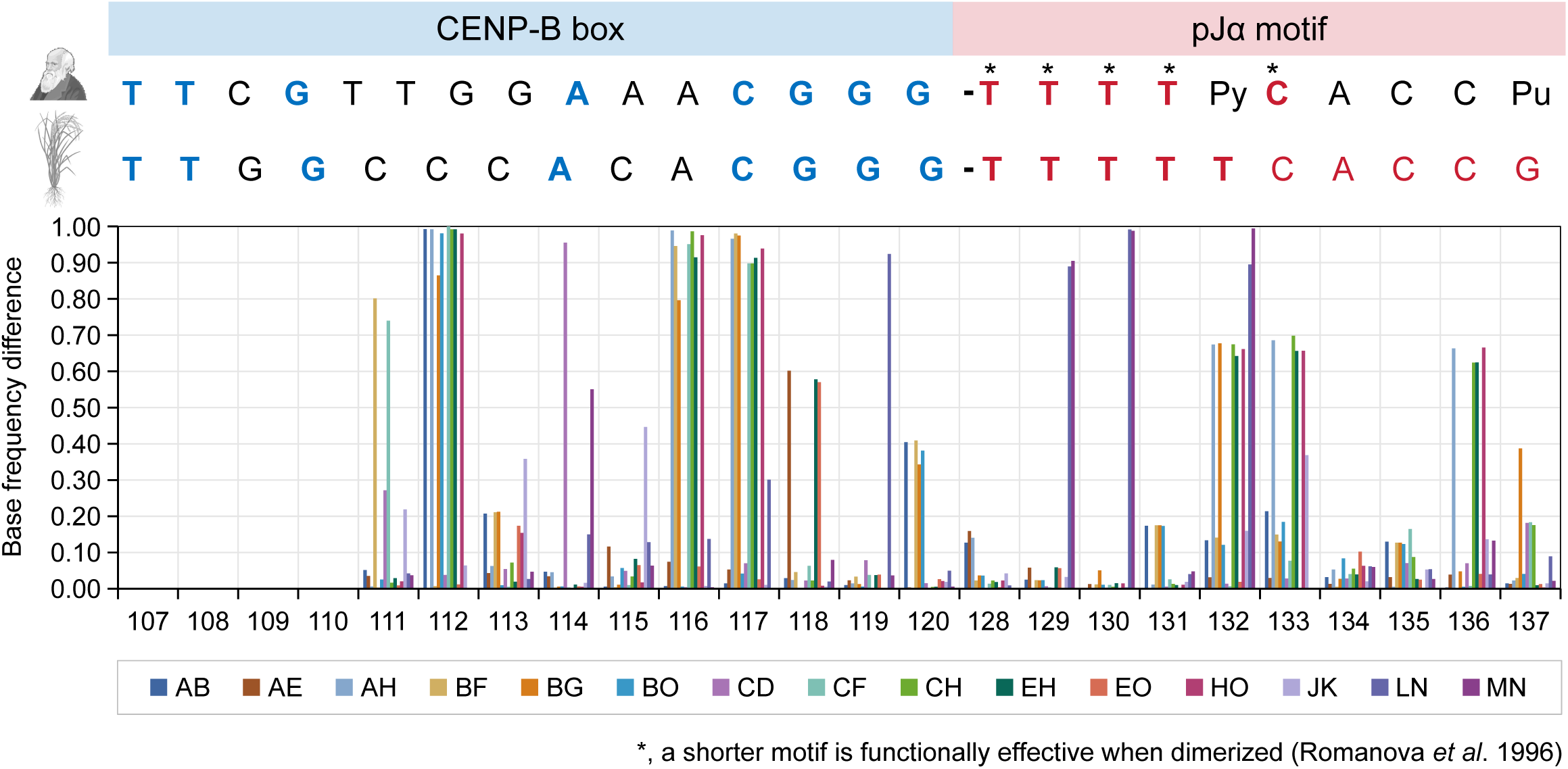
Divergence sites between *CEN155* superfamilies. The CENP-B box-like and pJα-like motif regions are shown.

**Extended Data Figure 5.**
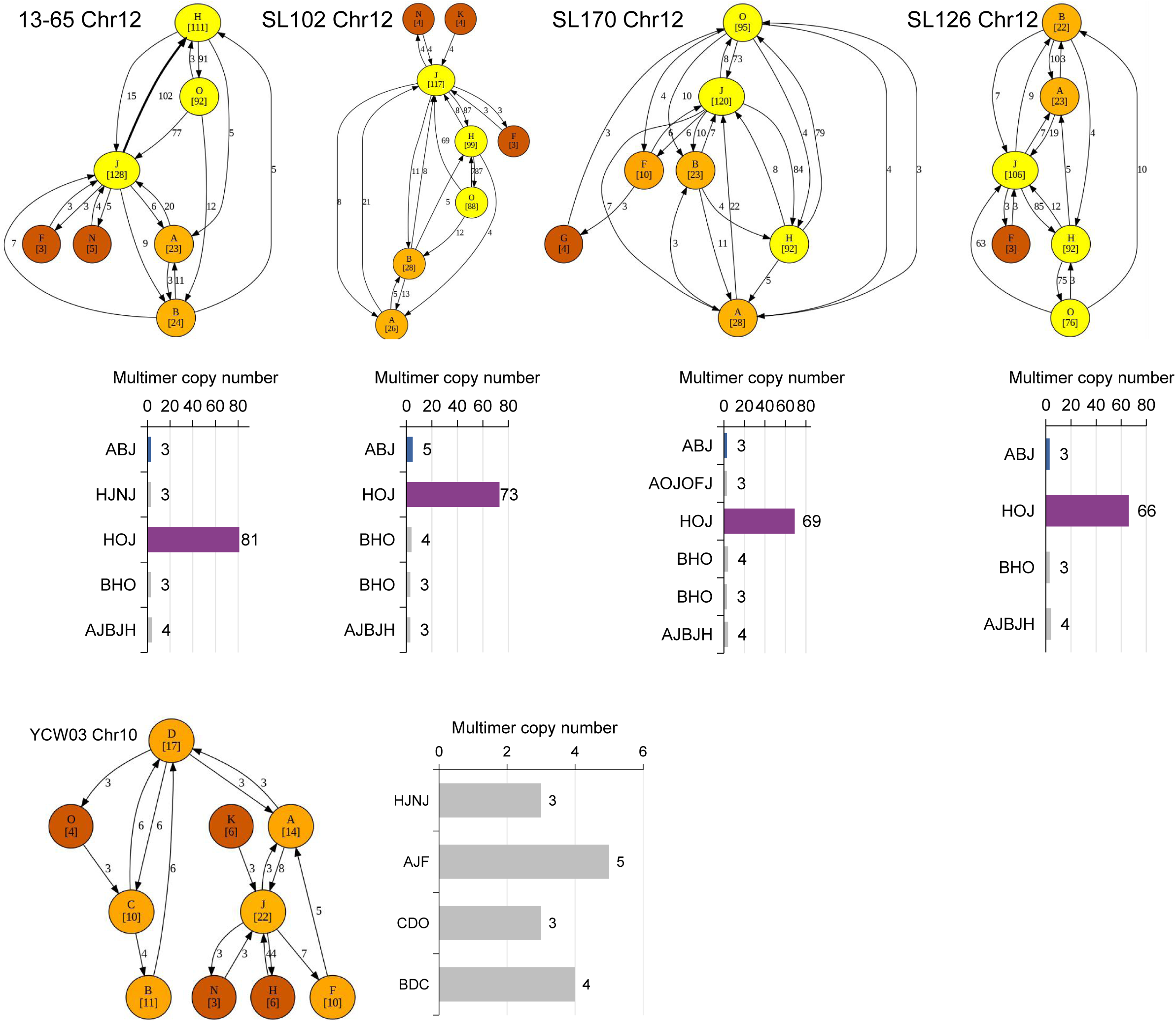
Inference of multimers and muHRs in rice centromeres. The de Bruijn graphs are constructed based on the dimer-compressed satellite string of each centromere.

**Extended Data Figure 6.**
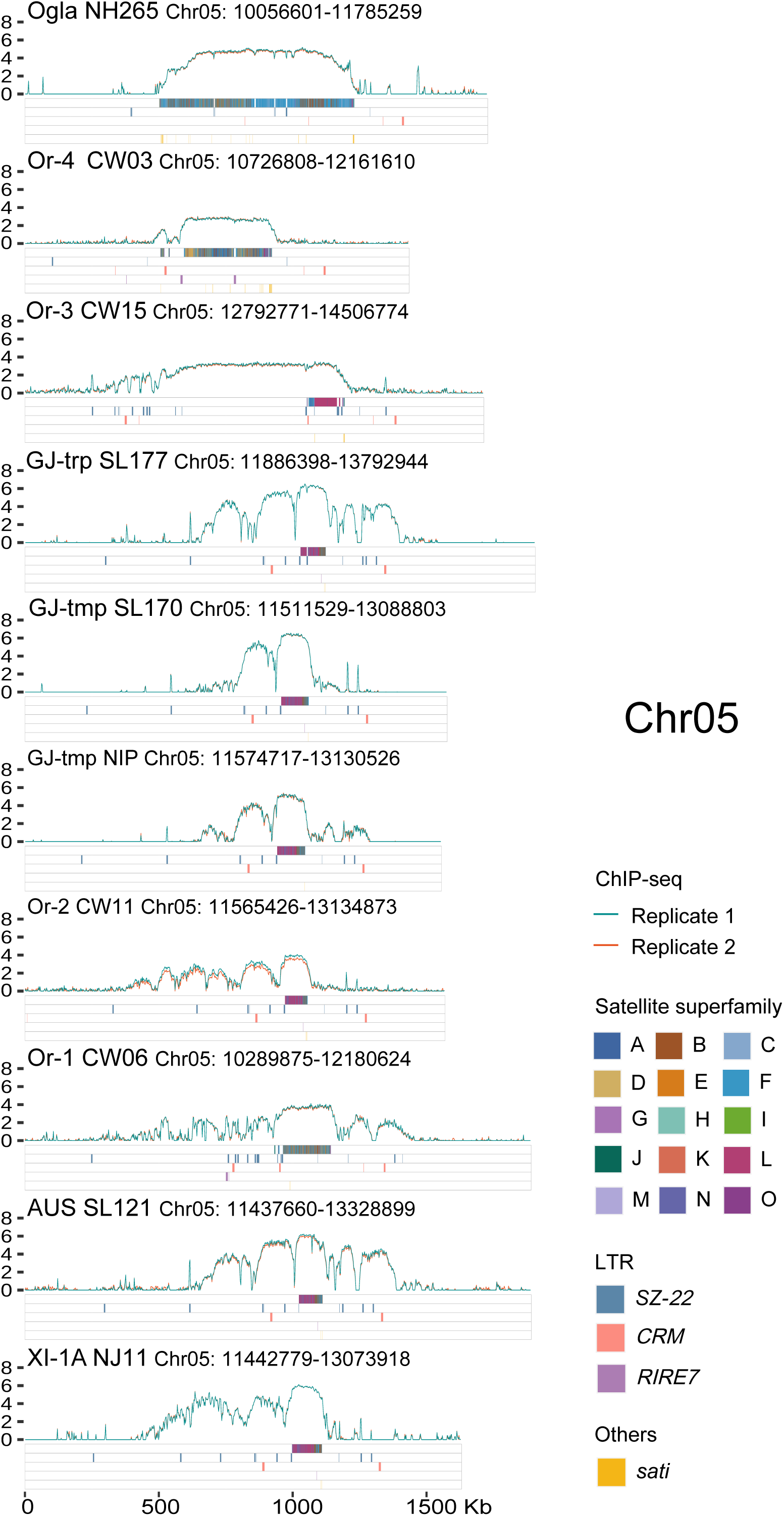
CENH3 ChIP enrichment and element annotation across Chr05 centromeres.

**Supplementary Fig. 1** | Genomic features and quality assessments of rice genomes in this study. QV, consensus quality value; AQI, assembly quality indicator value; GCI, assembly continuity index, derived from contig N50.

**Supplementary Fig. 2 |** Accuracy and continuity assessments of rice centromere assemblies. **a**, Base-level accuracy of centromere sequences represented by QV, where "+inf" indicates no erroneous bases in the centromere assemblies. **b**, Potential *k*-mer discordance in centromere assemblies of CW03 (Or-4), NH265 (Ogla), and NJ11 (XI1A), detected by VerityMap. **c**, CGI scores for rice centromere assemblies. **d**, Comparison of centromere assemblies generated by Hifiasm and Verkko, showing high consistency.

**Supplementary Fig. 3** | Relationship between centromere length and chromosome length across 12 chromosomes among different taxa.

**Supplementary Fig. 4** | Impacts of TEs on chromosome size variation. **a**, Relationship between centromere length and chromosome size within each chromosome. **b**, Relationship between the length of TE sequences outside of centromeres and chromosome size. **c**, Relationship between centromere length and chromosome size in *Arabidopsis thaliana* genomes.

**Supplementary Fig. 5** | Sequence similarity and haplotypes of rice centromeres revealing intra-species introgression and rearrangement events across chromosomes. Left, phylogenetic tree based on chromosome arm SNPs. Middle, heat maps showing the percentage of sequence similarity shared by pairwise centromeres. Right, haplotype assignments of rice centromeres.

**Supplementary Fig. 6** | StainedGlass sequence similarity heat maps comparing within-and between the Chr12 centromeres of CW09, SL044 and NJ11.

**Supplementary Fig. 7** | Distribution of satellite repeat length across the 70 rice genomes in each taxonomic group (Obar, Ogla, Oruf, AUS, GJ, and XI).

**Supplementary Fig. 8** | Characteristics of the fifteen *CEN155* superfamilies. **a**, Distribution of *CEN155* length within each superfamily. **b**, Edit distances among superfamilies.

**Supplementary Fig. 9** | TE invasion landscape in rice centromeres. **a**, TE proportions and accumulated lengths in centromere regions across chromosomes, colored by different TE super-families. **b**, TE proportions among rice genomes, shown according to the phylogenetic tree. From left to right, the circled diagrams represent repeat proportions in centromere, composition in centromeric TEs, and LTR composition in centromeric TEs, respectively. The size of the circle represents the sum of cumulative length. **c**, TE proportions across chromosomes with orange point representing centromere region and blue points stand for genome-wide region.

**Supplementary Fig. 10 |** Features of intact centrophilic LTR families (*SZ-22*, *RIRE7* and *CRM*) in rice centromeres. **a**, Phylogeny of all LTRs in the pan-TE library based on interval sequences, showing distant relationship between *SZ-22* and other two centrophilic LTRs. Triangles in different colors represent the domains of gag, RVT, RNaseH, Integrase_H2C2 and rve, respectively. **b**, Length distributions of *SZ-22*, *RIRE7* and *CRM* identified within the centromere, pericentromere, and chromosome arm regions.

**Supplementary Fig. 11** | Integration frequency of centrophilic LTR *SZ-22* along the *CEN155* consensus sequence.

**Supplementary Fig. 12** | Organization features of non-canonical satellites *sati94*. **a**, The genomic distance (measured by physical distance and *CEN155* copy number) between adjacent non-canonical satellite *sati94* in the three CEN155+ arrays indicating local homogenization. **b**, The maximum-likelihood phylogeny of *sati94* sequences from the three CEN155+ arrays in Chr10 centromeres of NH284 and NH265. Bootstrap values are shown at the branches.

**Supplementary Fig. 13** | Local homogenization at the monomer, dimer, and multimer scales. **a**, Distribution of *CEN155* satellites exhibiting monomer homogenization across the fifteen superfamilies in 70 rice accessions from various taxa. Numbers of dimers and multimers for the top fifteen groups are shown in **b** and **c**. Each rice accession is colored by taxonomic groups at the top, and summed up at the bottom. The total numbers of monomers, dimers, and multimers are showed on the right.

**Supplementary Fig. 14 |** Biased monomer copy numbers across dimers and taxonomic groups. One monomer has a relatively constrained and stabilized copy number (nearly 1), while the other exhibits variable copies (forming moHRs) within a diHR, represented by XY*_m_* or X*_m_*Y.

**Supplementary Fig. 15** | Local homogenization of copy number bias within a dimer through tandem duplication. **a**-**d**, Repeat of bias pattern in monomer copy number of dimers in NIP(GJ), NJ11(XI), SL022(AUS) and NH265(Ogla).

**Supplementary Fig. 16** | Workflow of SynPan-CEN. (1) Stepwise backbone building. All vs all pairwise satellite edit distance (ED) values were used to construct the Directed Acyclic Graph (DAG). To enhance accuracy in constructing synteny, the chain-backbone was built with the most conservative and highly-ordered monomer pairs by gradually increasing the ED value thresholds until the framework contained more than 20% of the total monomers or 100 monomers. This framework was then split by 10 monomer pairs within one window for downstream analysis. (2) Local satellite rescuing. DAGs were used again to rescue chains of syntenic satellite pairs within each window, employing relaxed EDs of 15. The sub-blocks with the highest scores were retained, and any non-overlapping monomer pairs were added to complete the matches. (3) Blocks from the chain-backbone were iteratively merged.

**Supplementary Fig. 17** | Synteny ratio of *CEN155* satellites in rice centromeres against NIP (GJ-tmp) centromeres.

**Supplementary Fig. 18** | Heat map showing the synteny ratio between NIP (GJ-tmp) and other rice centromeres across 12 chromosomes, with Chr05 centromeres showing a significantly higher syntenic relationship between XI and GJ.

**Supplementary Fig. 19 |** Satellite syntenic relationships between centromeres of J4155S, SL148, and XL628S with NJ11 on Chr05, and CX20, LK638S on Chr10. Tracks show centromere annotations, including TE families, *CEN155* strands, *CEN155* superfamilies, and edit distances, respectively.

**Supplementary Fig. 20 |** Genomic features of the functional centromere and its flanking regions on chromosome Chr01. Top, CENH3 ChIP-seq enrichment (log_2_(ChIP/input), two replicates) in 10-Kb windows. Track 1, satellite superfamilies; Track 2, LTRs *SZ-22*; Track 3, LTRs *CRM*; Track 4, LTRs *RIRE7*; Track 5, *sati* (non-canonical satellites).

**Supplementary Fig. 21 |** Genomic features of the functional centromere and its flanking regions on chromosome Chr02. Top, CENH3 ChIP-seq enrichment (log_2_(ChIP/input), two replicates) in 10-Kb windows. Track 1, satellite superfamilies; Track 2, LTRs *SZ-22*; Track 3, LTRs *CRM*; Track 4, LTRs *RIRE7*; Track 5, *sati* (non-canonical satellites).

**Supplementary Fig. 22** | Genomic features of the functional centromere and its flanking regions on chromosome Chr03. Top, CENH3 ChIP-seq enrichment (log_2_(ChIP/input), two replicates) in 10-Kb windows. Track 1, satellite superfamilies; Track 2, LTRs *SZ-22*; Track 3, LTRs *CRM*; Track 4, LTRs *RIRE7*; Track 5, *sati* (non-canonical satellites).

**Supplementary Fig. 23** | Genomic features of the functional centromere and its flanking regions on chromosome Chr06. Top, CENH3 ChIP-seq enrichment (log_2_(ChIP/input), two replicates) in 10-Kb windows. Track 1, satellite superfamilies; Track 2, LTRs *SZ-22*; Track 3, LTRs *CRM*; Track 4, LTRs *RIRE7*; Track 5, *sati* (non-canonical satellites).

**Supplementary Fig. 24 |** Genomic features of the functional centromere and its flanking regions on chromosome Chr07. Top, CENH3 ChIP-seq enrichment (log_2_(ChIP/input), two replicates) in 10-Kb windows. Track 1, satellite superfamilies; Track 2, LTRs *SZ-22*; Track 3, LTRs *CRM*; Track 4, LTRs *RIRE7*; Track 5, *sati* (non-canonical satellites).

**Supplementary Fig. 25** | Genomic features of the functional centromere and its flanking regions on chromosome Chr08. Top, CENH3 ChIP-seq enrichment (log_2_(ChIP/input), two replicates) in 10-Kb windows. Track 1, satellite superfamilies; Track 2, LTRs *SZ-22*; Track 3, LTRs *CRM*; Track 4, LTRs *RIRE7*; Track 5, *sati* (non-canonical satellites).

**Supplementary Fig. 26** | Genomic features of the functional centromere and its flanking regions on chromosome Chr09. Top, CENH3 ChIP-seq enrichment (log_2_(ChIP/input), two replicates) in 10-Kb windows. Track 1, satellite superfamilies; Track 2, LTRs *SZ-22*; Track 3, LTRs *CRM*; Track 4, LTRs *RIRE7*; Track 5, *sati* (non-canonical satellites).

**Supplementary Fig. 27** | Genomic features of the functional centromere and its flanking regions on chromosome Chr10. Top, CENH3 ChIP-seq enrichment (log_2_(ChIP/input), two replicates) in 10-Kb windows. Track 1, satellite superfamilies; Track 2, LTRs *SZ-22*; Track 3, LTRs *CRM*; Track 4, LTRs *RIRE7*; Track 5, *sati* (non-canonical satellites).

**Supplementary Fig. 28** | Genomic features of the functional centromere and its flanking regions on chromosome Chr11. Top, CENH3 ChIP-seq enrichment (log_2_(ChIP/input), two replicates) in 10-Kb windows. Track 1, satellite superfamilies; Track 2, LTRs *SZ-22*; Track 3, LTRs *CRM*; Track 4, LTRs *RIRE7*; Track 5, *sati* (non-canonical satellites).

**Supplementary Fig. 29** | Genomic features of the functional centromere and its flanking regions on chromosome Chr12. Top, CENH3 ChIP-seq enrichment (log_2_(ChIP/input), two replicates) in 10-Kb windows. Track 1, satellite superfamilies; Track 2, LTRs *SZ-22*; Track 3, LTRs *CRM*; Track 4, LTRs *RIRE7*; Track 5, *sati* (non-canonical satellites).

**Supplementary Fig. 30 |** Correlation between the length of CENH3 enriched regions and *CEN155*-defined centromere size for each chromosome.

**Supplementary Fig. 31** | Correlation between chromosome length and TE density across the 12 chromosomes.

**Supplementary Fig. 32** | Correlation between the chromosome length and *CEN155* array length across the 12 chromosomes.

**Supplementary Fig. 33** | Correlation between the chromosome length and CENH3 enrichment length across the 12 chromosomes.

**Supplementary Fig. 34** | Correlation between the *CEN155* array length and neocentromere formation tendency (NFT) across 12 chromosomes.

**Supplementary Fig. 35** | Correlation between the TE density in the *CEN155* array and neocentromere formation tendency (NFT) across 12 chromosomes.

**Supplementary Fig. 36 |** Correlation between the chromosome length and neocentromere formation tendency (NFT) across 12 chromosomes.

**Supplementary Fig. 37** | Comparison of profiled DNA methylation probability from ONT sequencing data and HiFi data. Each point represents a randomly selected 5mC site.

**Supplementary Table 1** | The source, sequencing, assembly and QC information of 70 rice genomes in this study.

**Supplementary Table 2** | Detailed annotations of centromeres and chromosome arms in the 70 rice genomes.

**Supplementary Table 3** | Gaps in the rice centromere assemblies.

**Supplementary Table 4** | Putative centromere introgression events in rice evolution. **Supplementary Table 5** | Frequency of non-canonical satellites (*sati*) in rice centromeres. **Supplementary Table 6** | Satellite annotation in moHRs.

**Supplementary Table 7** | Satellite annotation in diHRs.

**Supplementary Table 8** | Satellite annotation in muHRs.

**Supplementary Table 9** | Centromere structural variations (CSVs) in rice genomes.

**Supplementary Table 10** | Relative divergence and mutation rate estimation for rice centromeres.

**Supplementary Table 11** | Functional centromeres in rice genomes defined by CENH3 ChIP data.

