## Supplementary Figs for "Genetic diversity and evolution of rice centromeres"

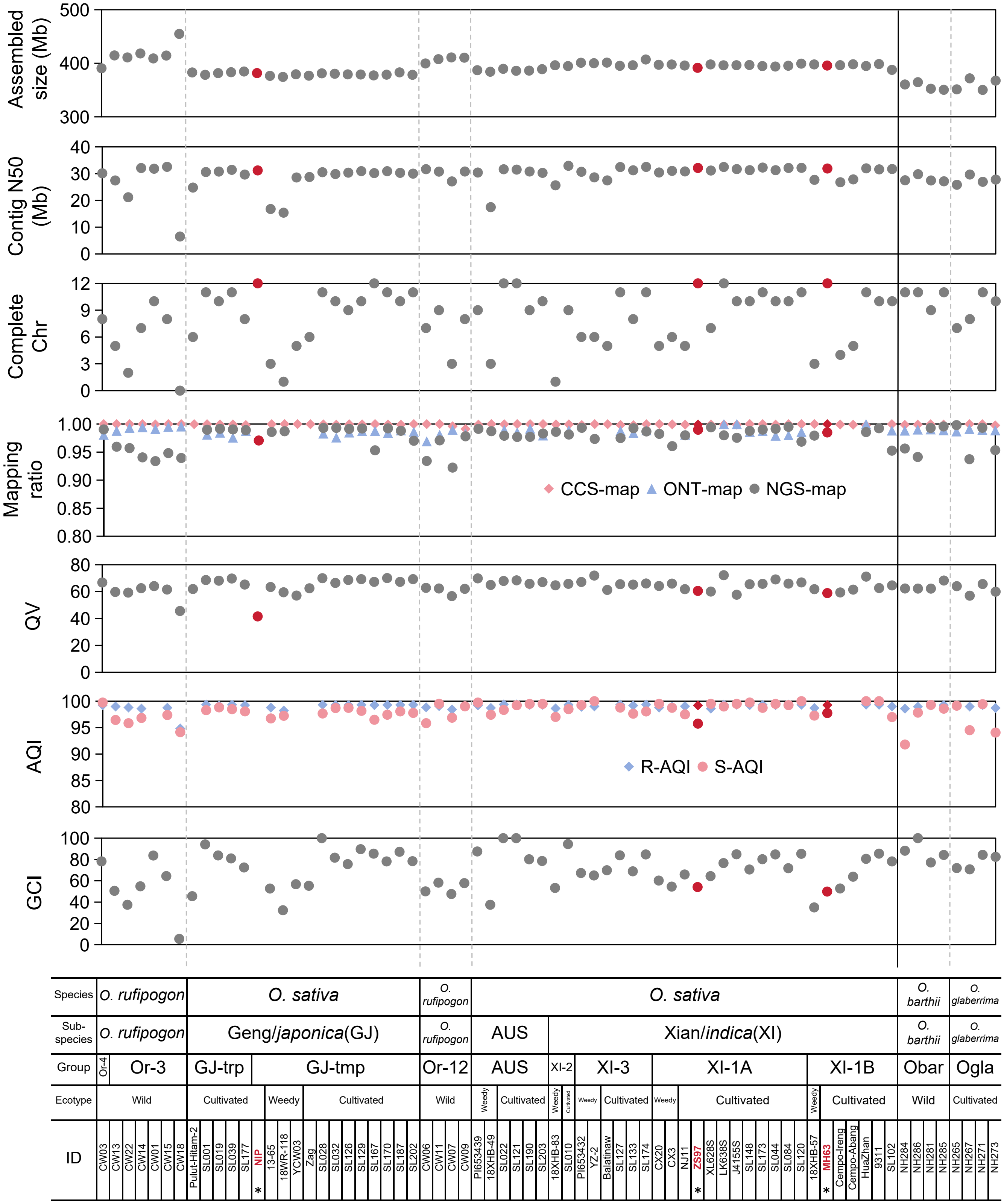


**Supplementary Fig. 1** | Genomic features and quality assessments of rice genomes in this study. QV, consensus quality value; AQI, assembly quality indicator value; GCI, assembly continuity index, derived from contig N50.

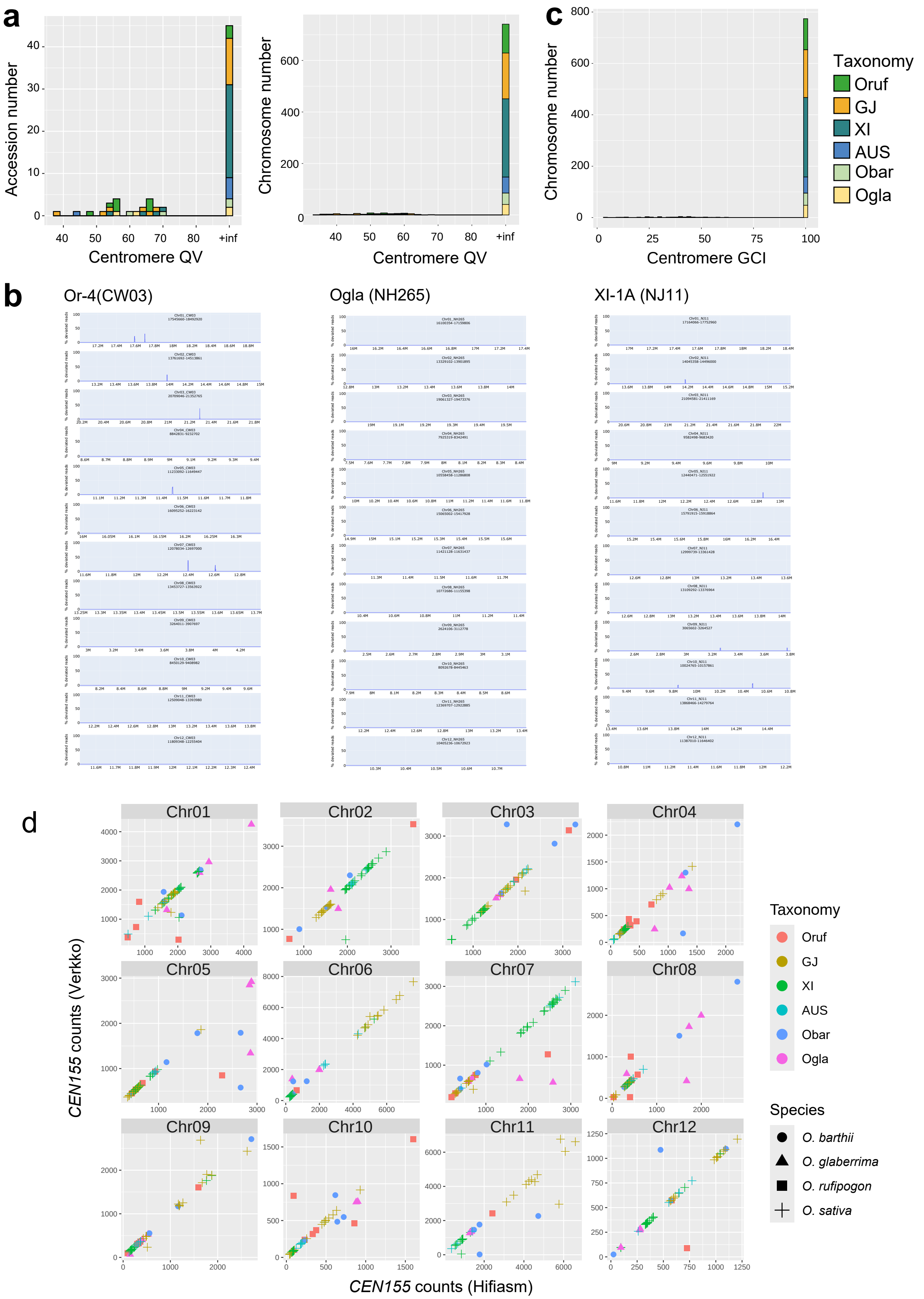


**Supplementary Fig. 2** | Accuracy and continuity assessments of rice centromere assemblies. **a**, Base-level accuracy of centromere sequences represented by QV, where "+inf" indicates no erroneous bases in the centromere assemblies. **b**, Potential *k*-mer discordance in centromere assemblies of CW03 (Or-4), NH265 (Ogla), and NJ11 (XI1A), detected by VerityMap. **c**, CGI scores for rice centromere assemblies. **d**, Comparison of centromere assemblies generated by Hifiasm and Verkko, showing high consistency.

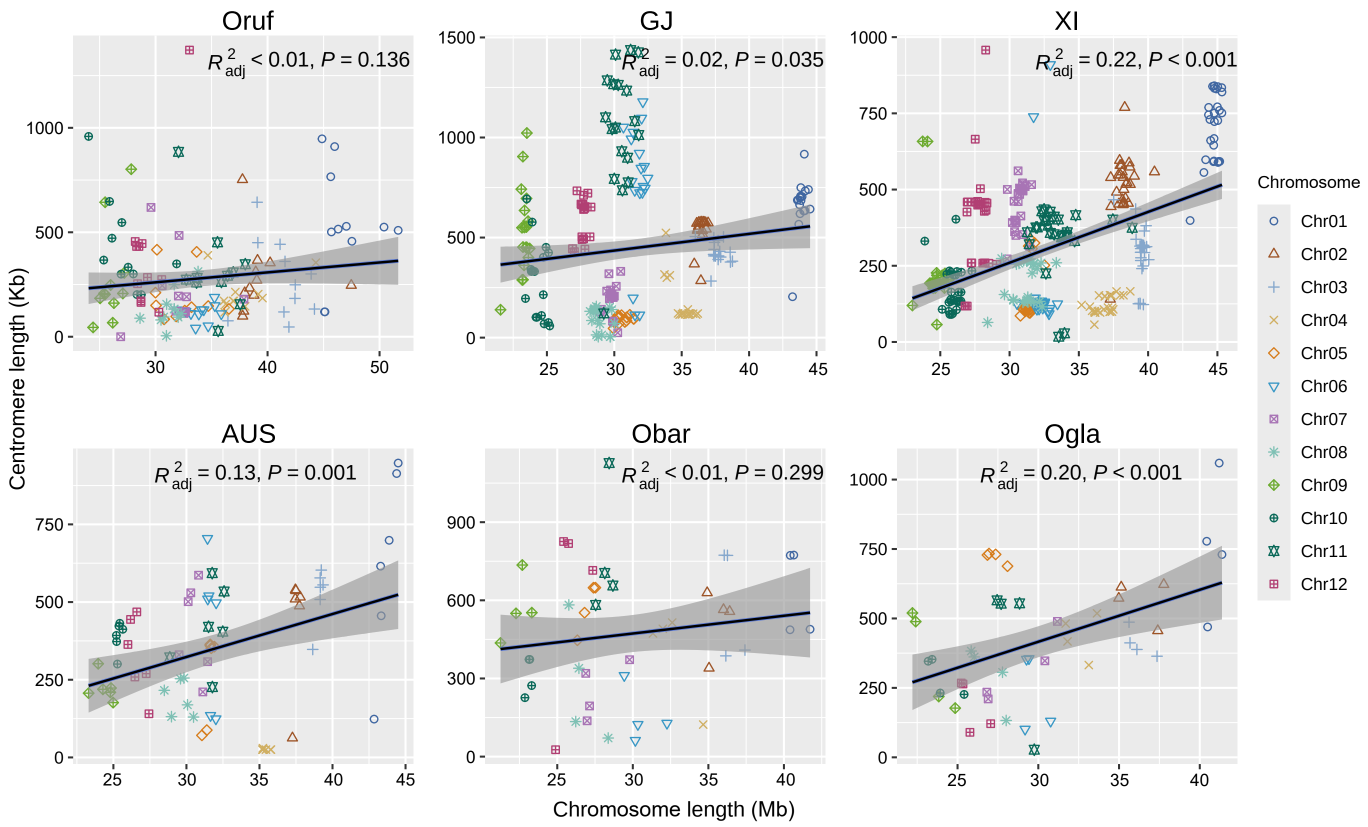


**Supplementary Fig. 3** | Relationship between centromere length and chromosome length across 12 chromosomes among different taxa.

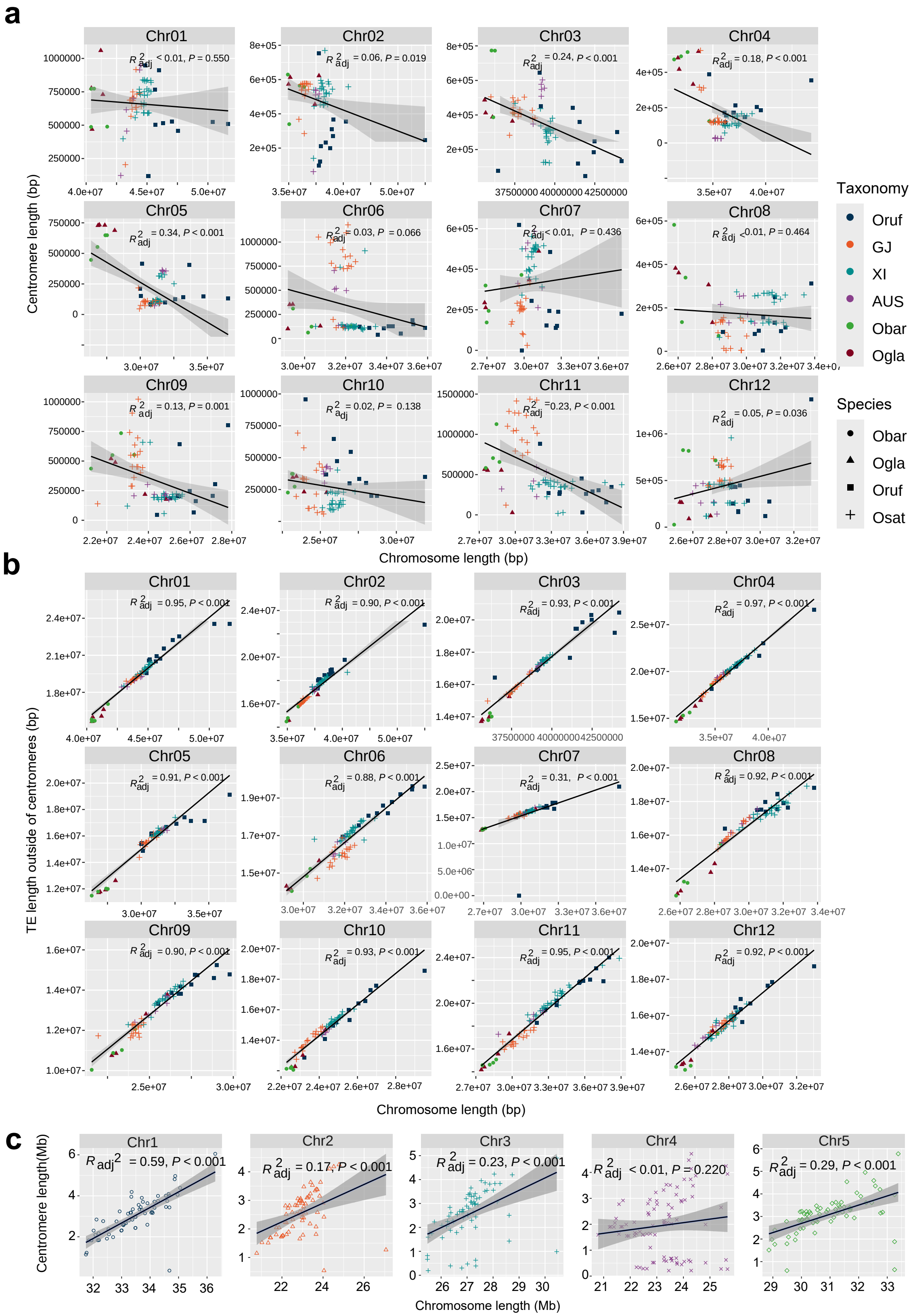


**Supplementary Fig. 4 |** Impacts of TEs on chromosome size variation. **a**, Relationship between centromere length and chromosome size within each chromosome. **b**, Relationship between TE sequence length outside of centromeres and chromosome size. c, Relationship between centromere length and chromosome size in *Arabidopsis thaliana* genomes.

### Chromosome 1

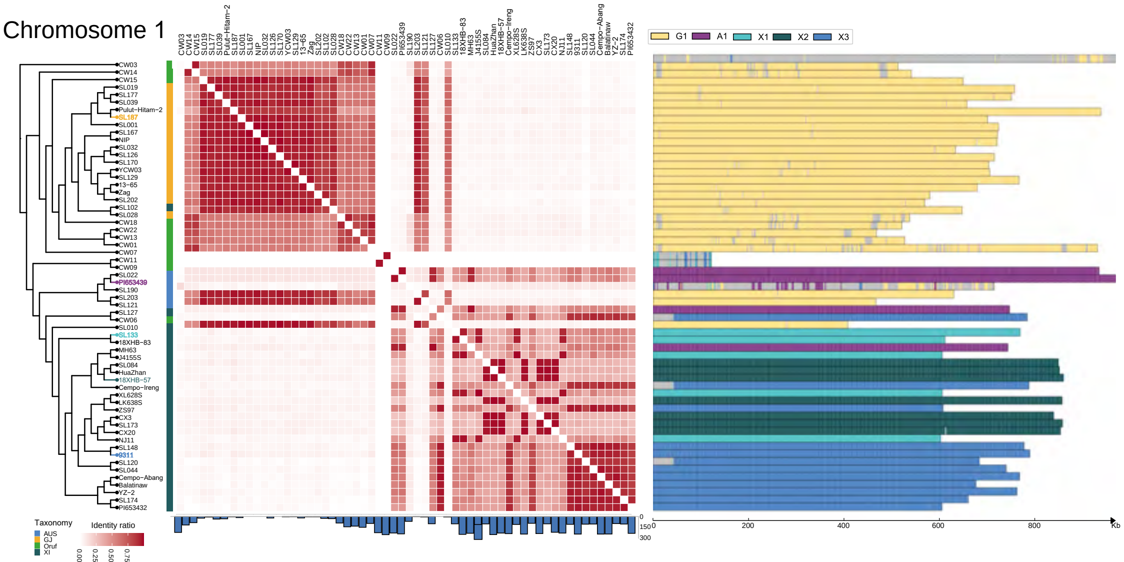


### Chromosome 3

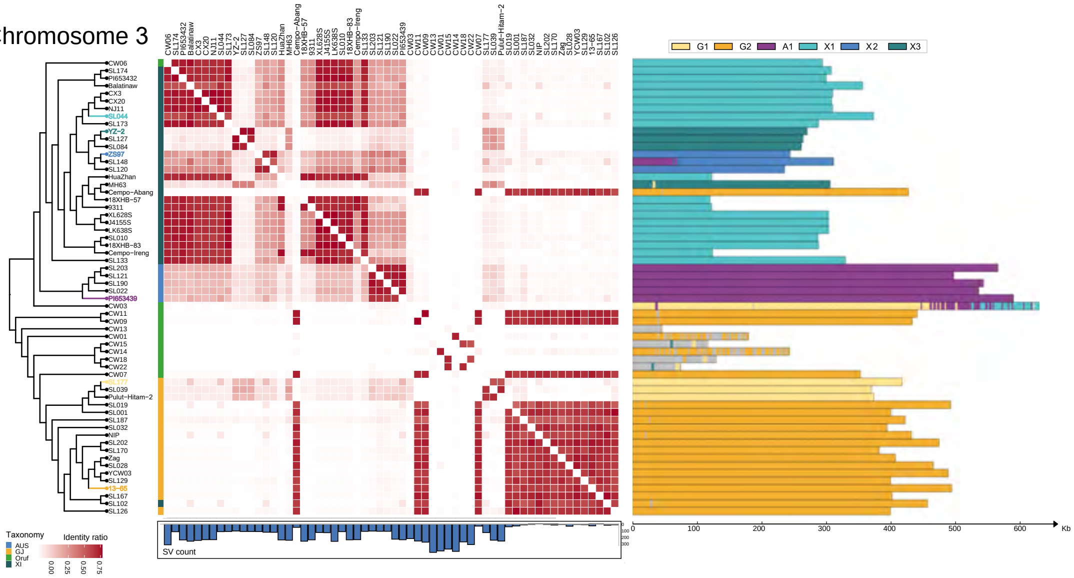


### Chromosome 6

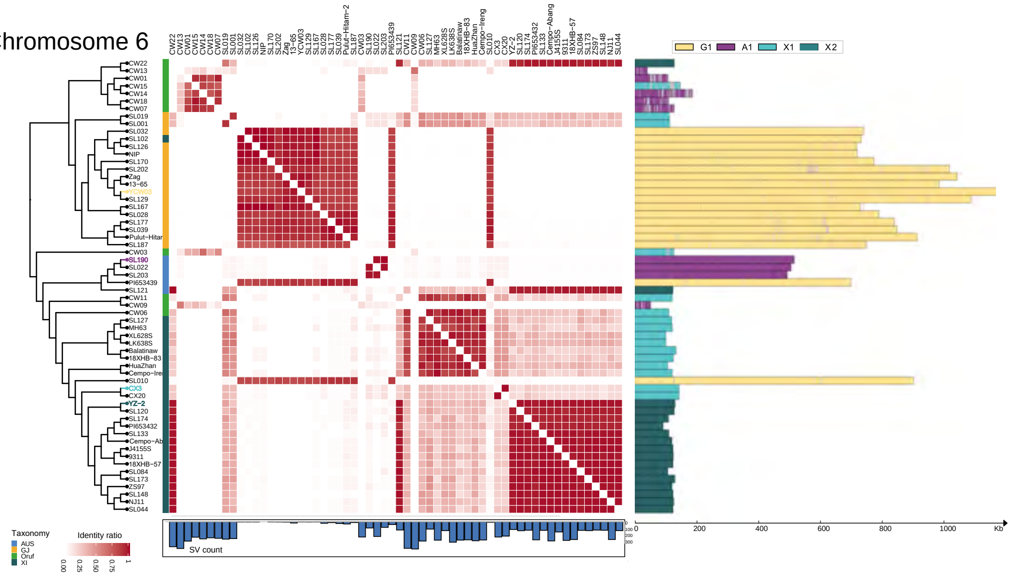

Chromosome 7

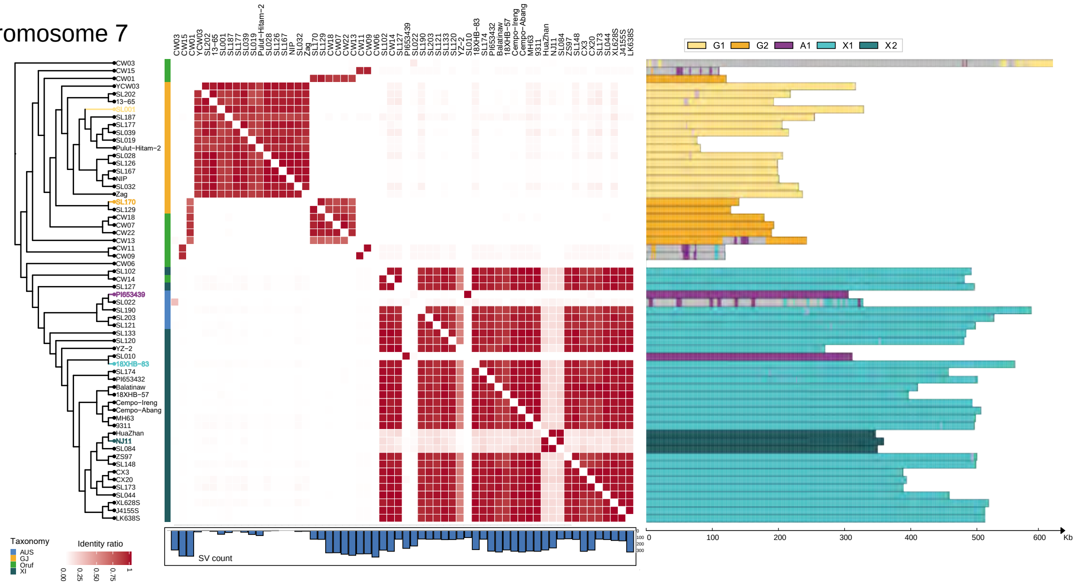


Chromosome 8

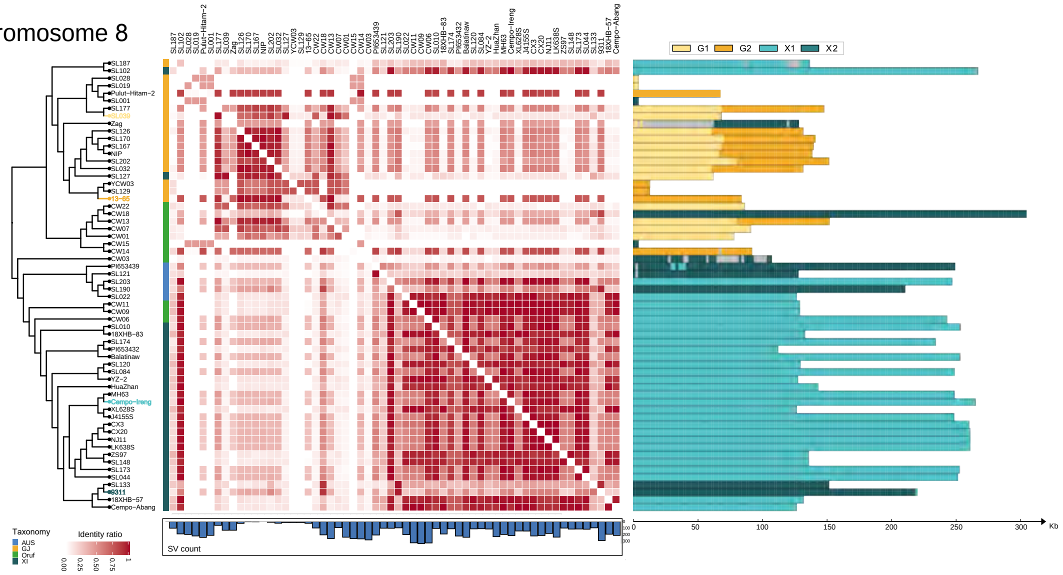


Chromosome 9

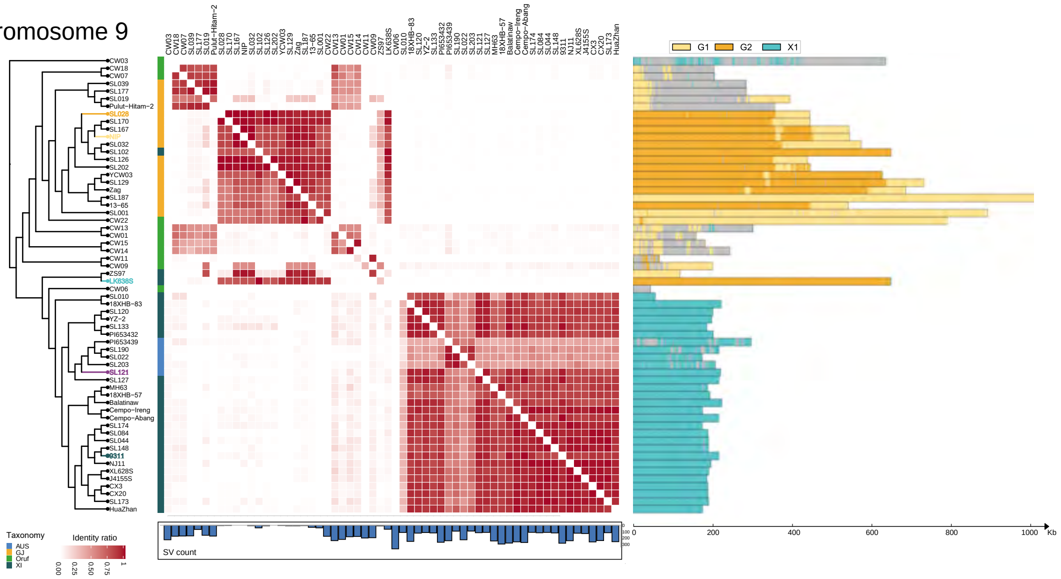

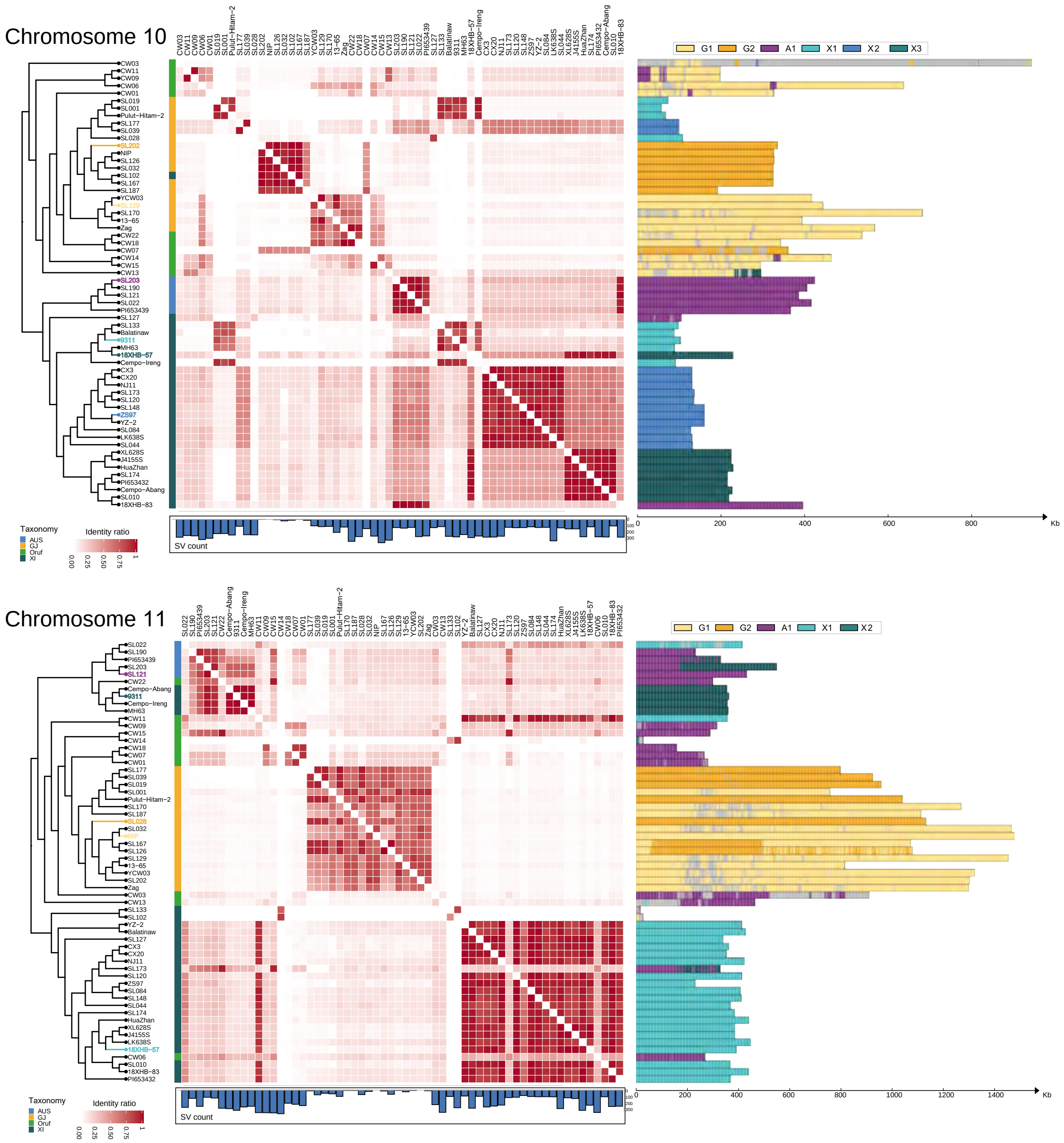


**Supplementary Fig. 5 |** Sequence similarity and haplotypes of rice centromeres revealing intra-species introgression and rearrangement events across chromosomes. Left, phylogenetic tree based on chromosome arm SNPs. Middle, heat maps showing the percentage of sequence similarity shared by pairwise centromeres. Right, haplotype assignments of rice centromeres.

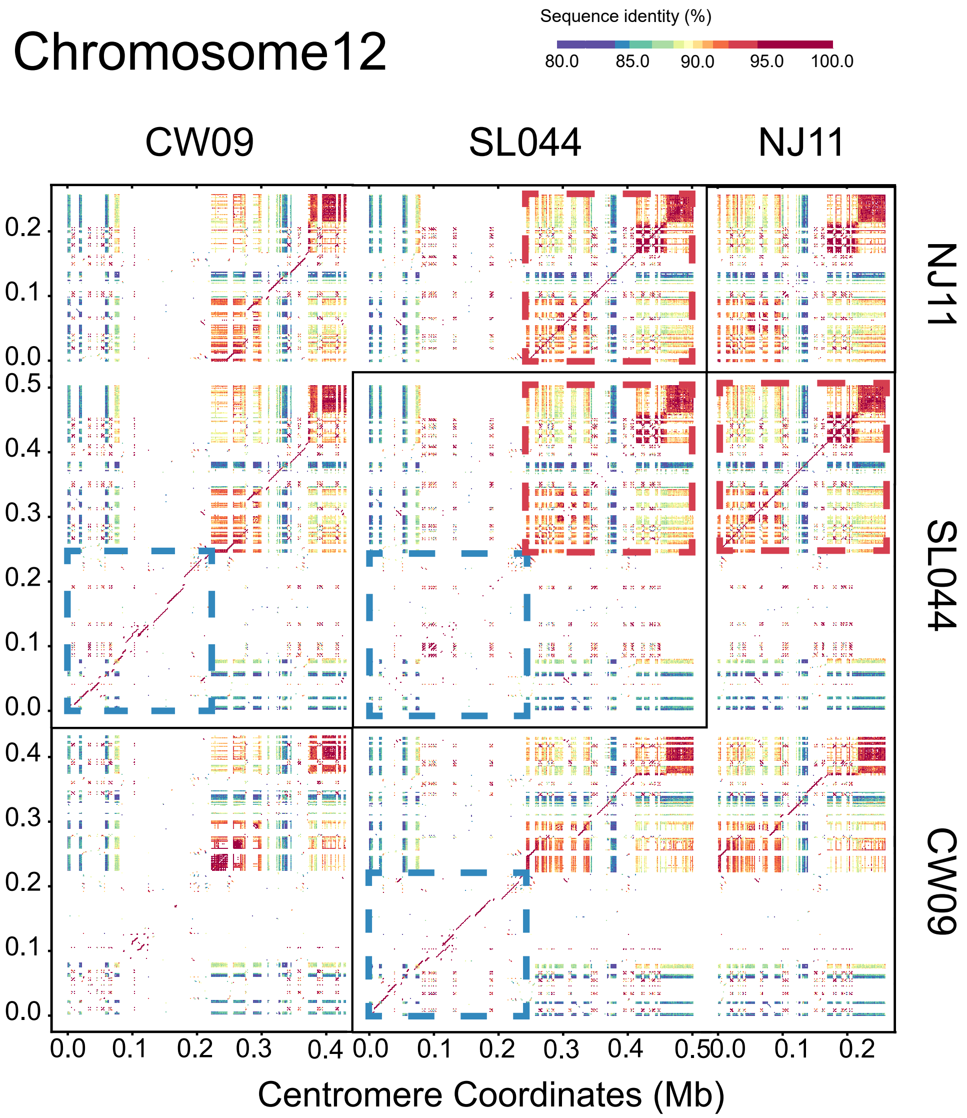


**Supplementary Fig. 6** | StainedGlass sequence similarity heat maps comparing within- and between the Chr12 centromeres of CW09, SL044 and NJ11.

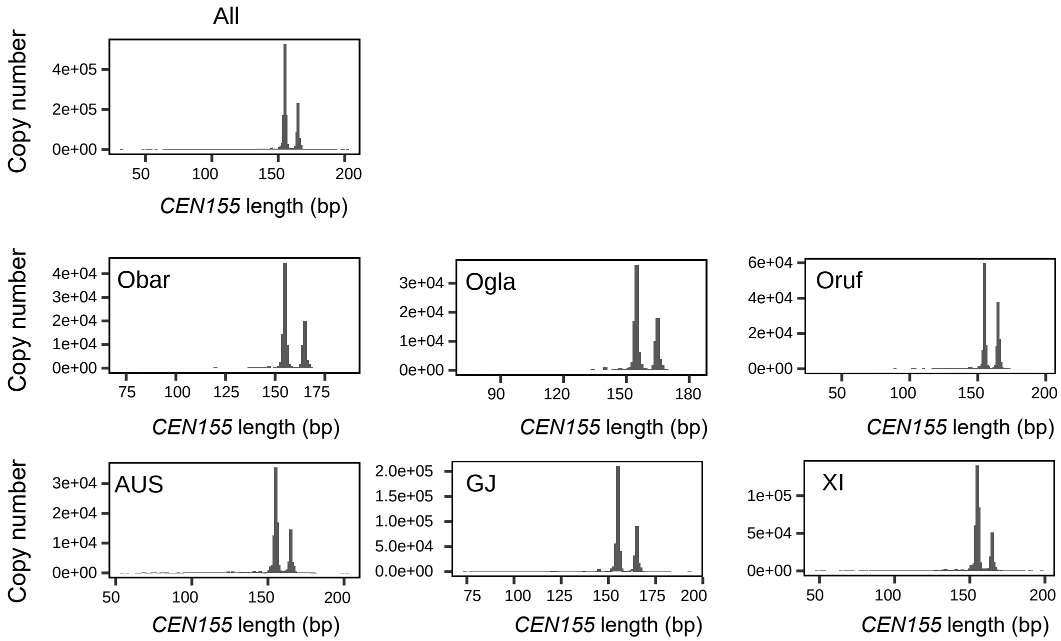


**Supplementary Fig. 7 |** Satellite repeat length across 70 rice accessions in each taxonomic group (Obar, Ogla, Oruf, AUS, GJ, and XI).

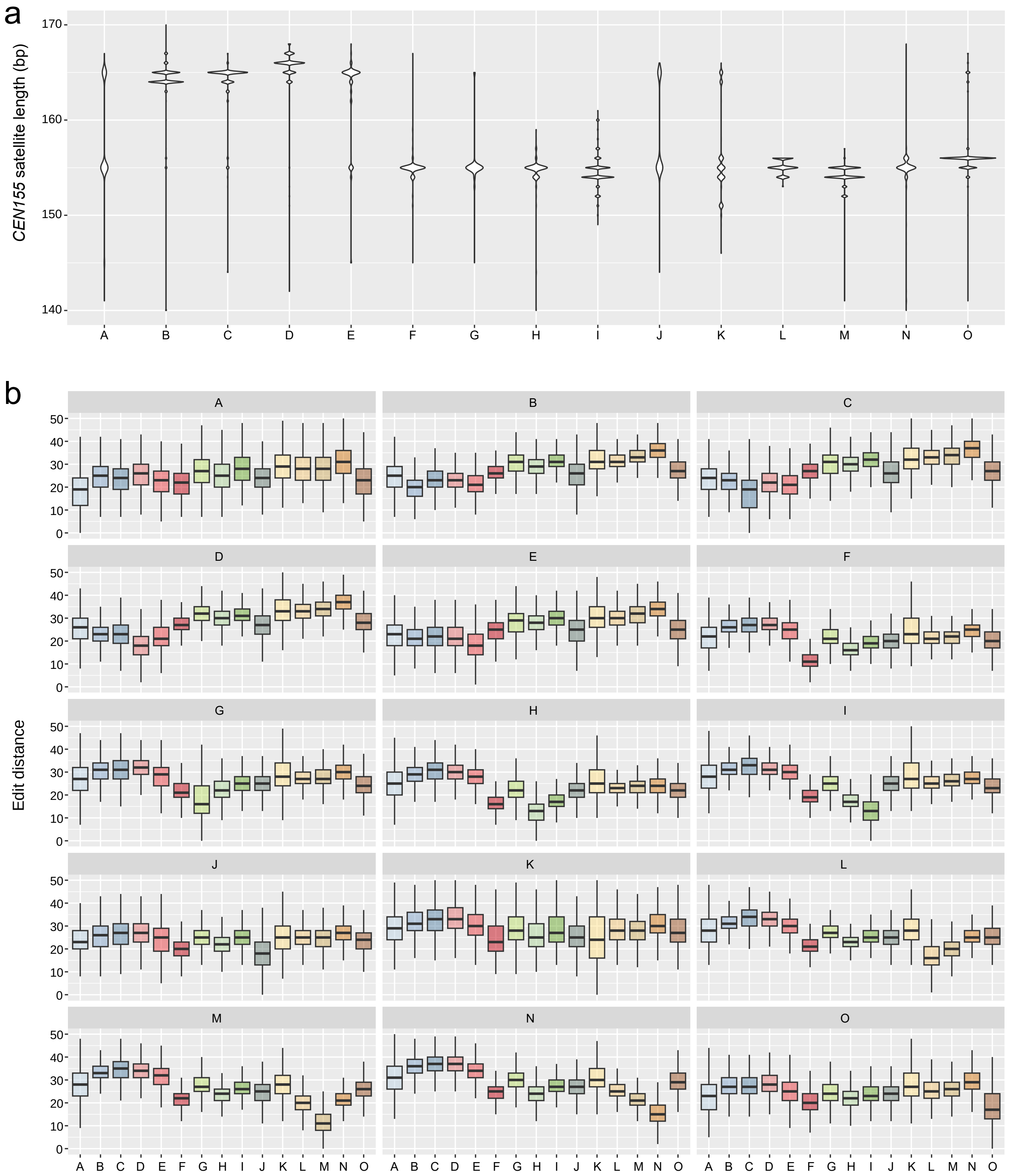


**Supplementary Fig. 8 |** Characteristics of the fifteen *CEN155* superfamilies. **a**, Distribution of *CEN155* length within each superfamily. **b**, Edit distances among superfamilies.

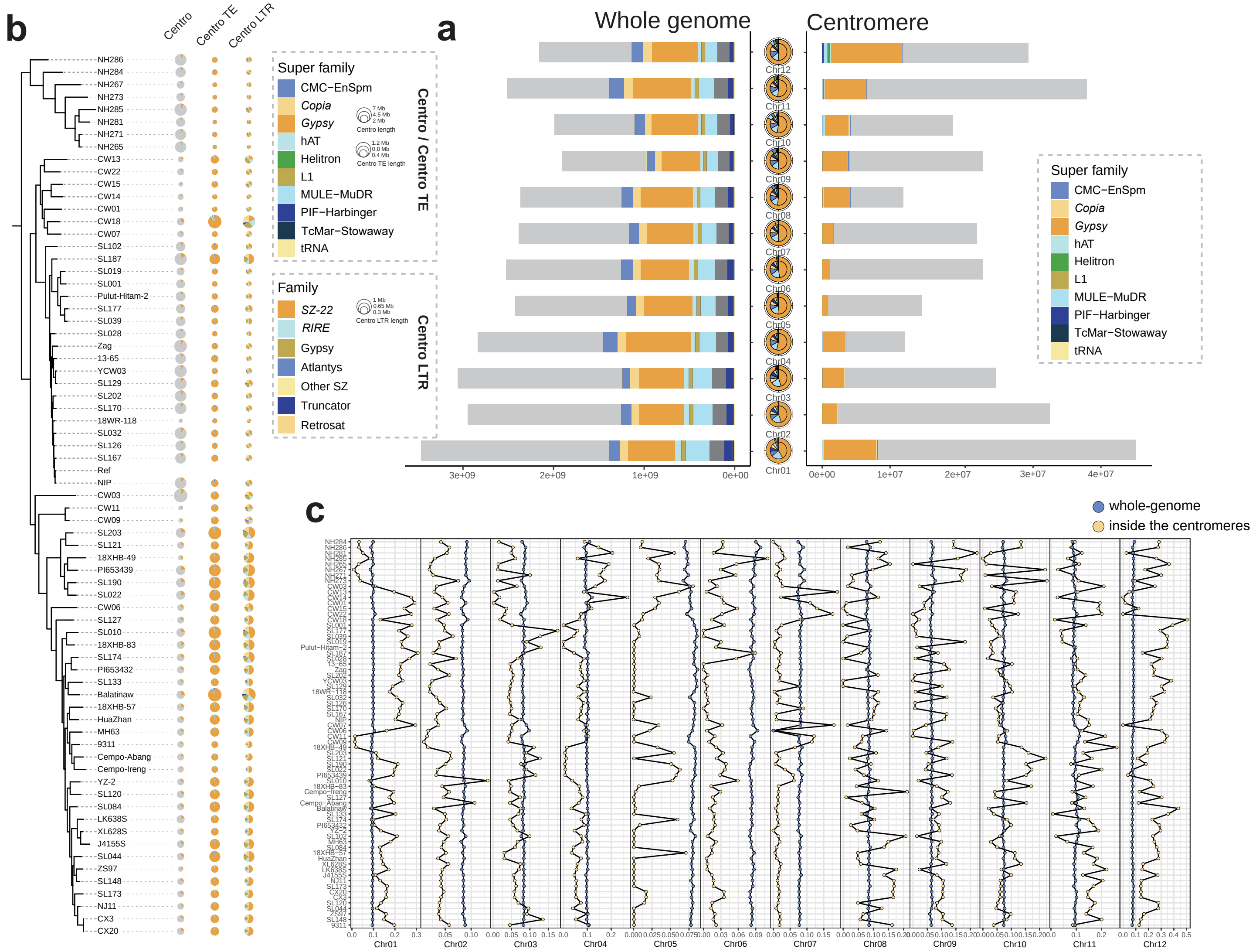


**Supplementary Fig. 9 |** TE invasion landscape in rice centromeres. **a**, TE proportions and accumulated lengths in centromere regions across chromosomes, colored by different TE super-families. **b**, TE proportions among rice genomes, shown according to the phylogenetic tree. From left to right, the circled diagrams represent repeat proportions in centromere, composition in centromeric TEs, and LTR composition in centromeric TEs, respectively. The size of the circle represents the sum of cumulative length. **c**, TE proportions across chromosomes with orange point representing centromere region and blue points stand for genome-wide region.

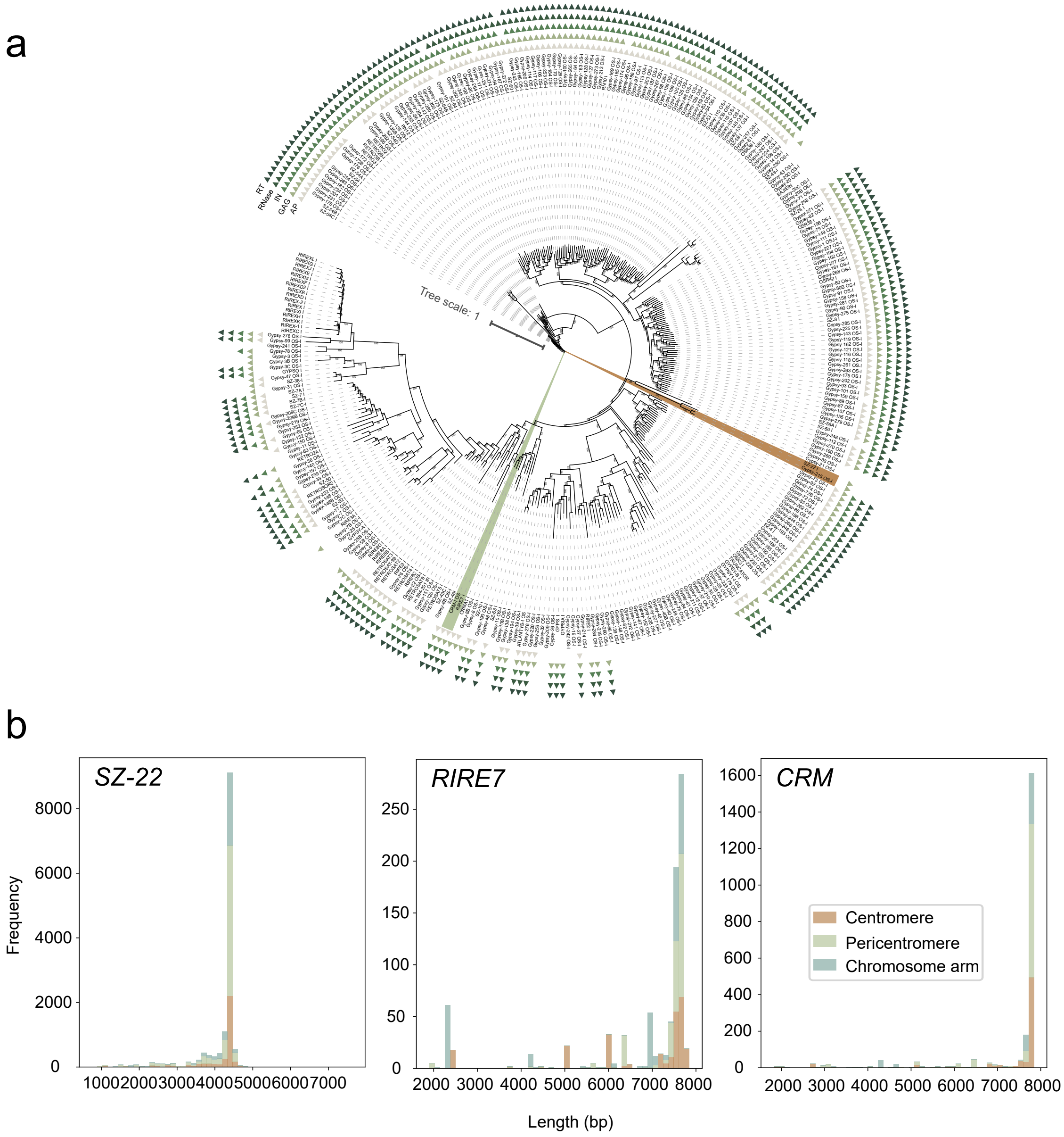


**Supplementary Fig. 10 |** Features of intact centrophilic LTR families (*SZ-22*, *RIRE7* and *CRM*) in rice centromere. **a**, Phylogeny of all LTRs in the pan-TE library based on interval sequences, showing distant relationship between *SZ-22* and other two centrophilic LTRs. Triangles in different colors were assigned to domain gag, RVT, RNaseH, Integrase_H2C2 and rve, respectively. **b**, Length distributions of *SZ-22*, *RIRE7,* and *CRM* identified within centromere, pericentromere, and chromosome arm regions.

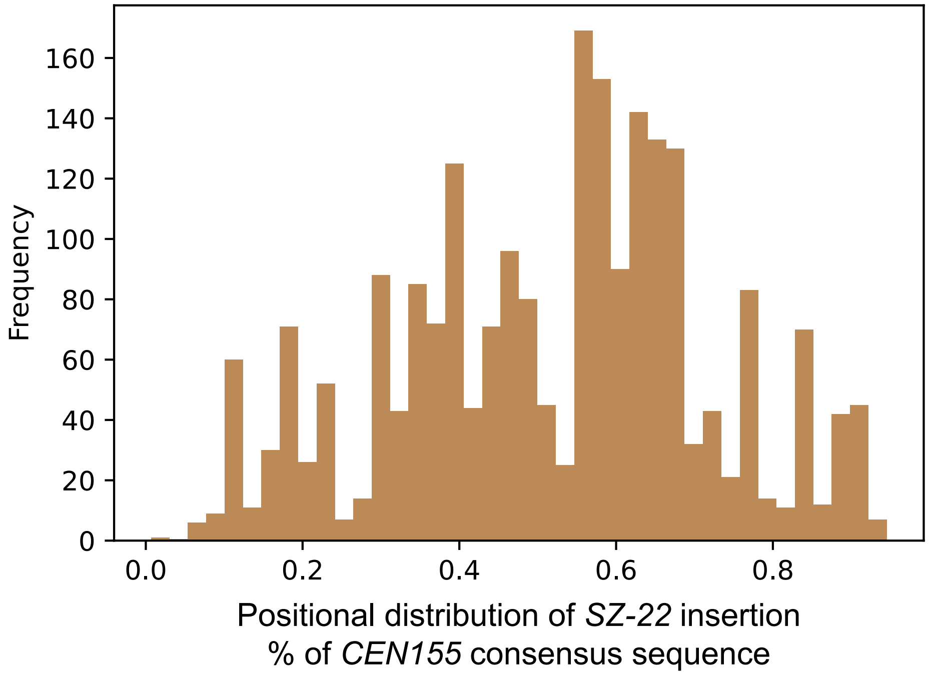


**Supplementary Fig. 11 |** Integration frequency of centrophilic LTR *SZ-22* along the *CEN155* consensus sequence.

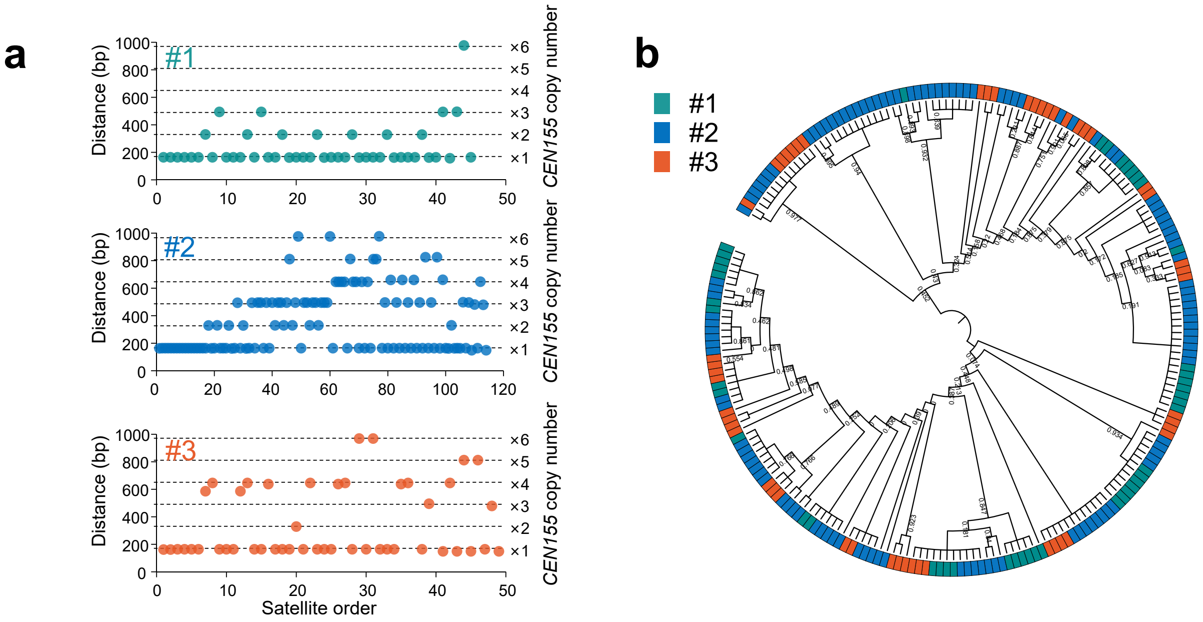


**Supplementary Fig. 12 |** Organization features of non-canonical satellites *sati94*. **a**, The genomic distances (measured by physical distance and *CEN155* copy number) between adjacent non-canonical satellite *sati94* in the three CEN155+ arrays indicating local homogenization. **b**, The maximum-likelihood phylogeny of *sati94* sequences from the three CEN155+ arrays in Chr10 centromeres of NH284 and NH265. Bootstrap values are shown at the branch.

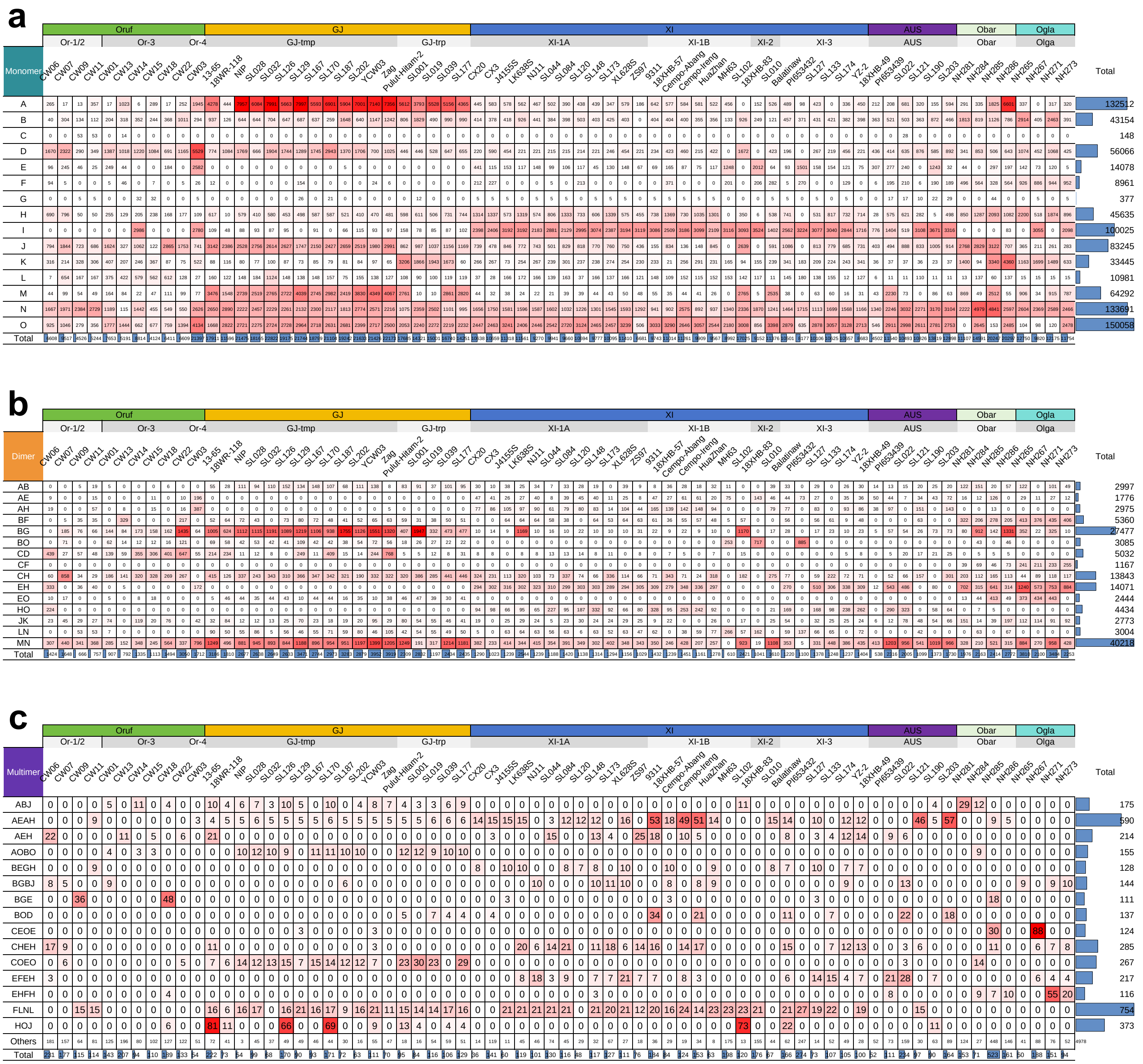


**Supplementary Fig. 13 |** Local homogenization at the monomer, dimer, and multimer scales. **a**, Distribution of *CEN155* satellites exhibiting monomer homogenization across the fifteen superfamilies in 70 rice accessions from various taxa. Numbers of dimer and multimer for the top fifteen groups are shown in **b** and **c**. Each rice accession is colored by taxonomic groups at the top, and summed up at the bottom. The total numbers of monomers, dimers, and multimers are showed on the right.

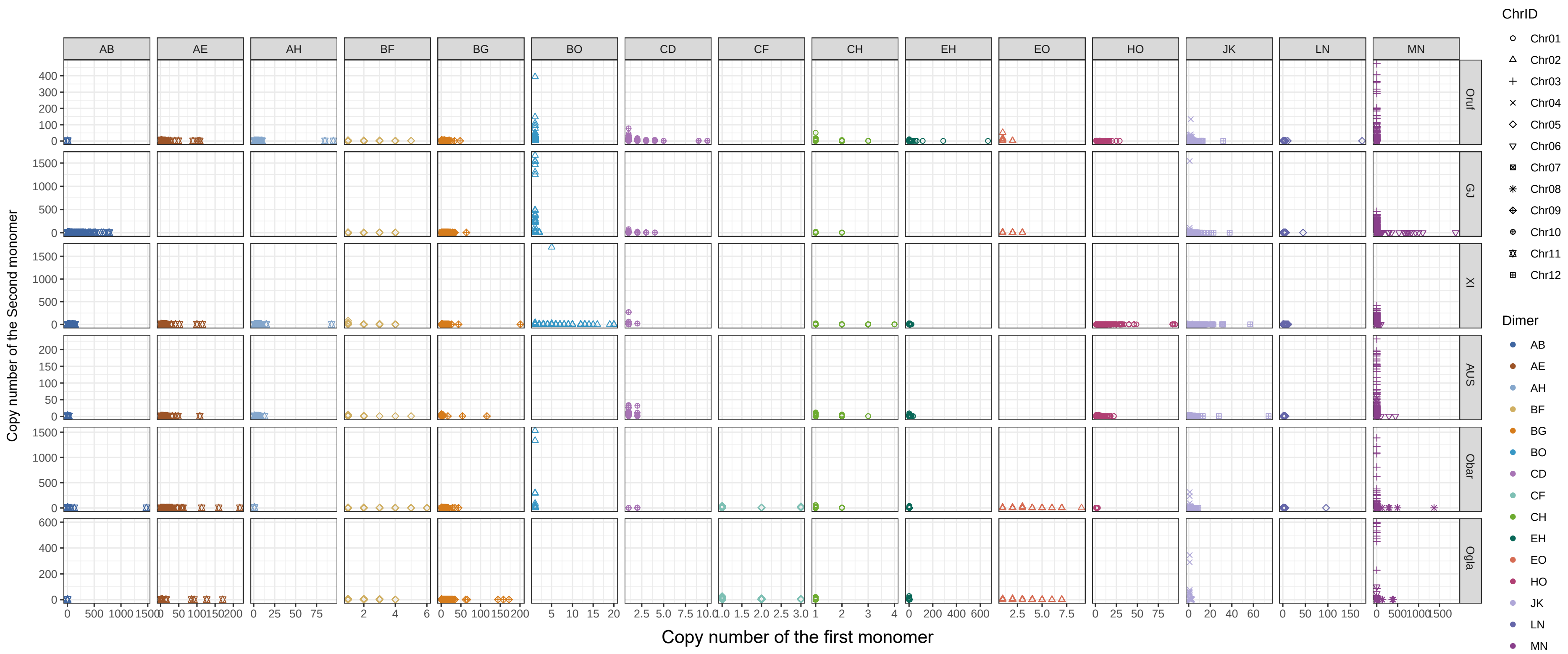


**Supplementary Fig. 14 |** Biased monomer copy number acorss dimers and taxonomic groups. One monomer has a relatively constrained and stabilized copy number (nearly 1), while the other exhibits variable copies (forming moHRs) within a diHR, represented by XYm or XmY.

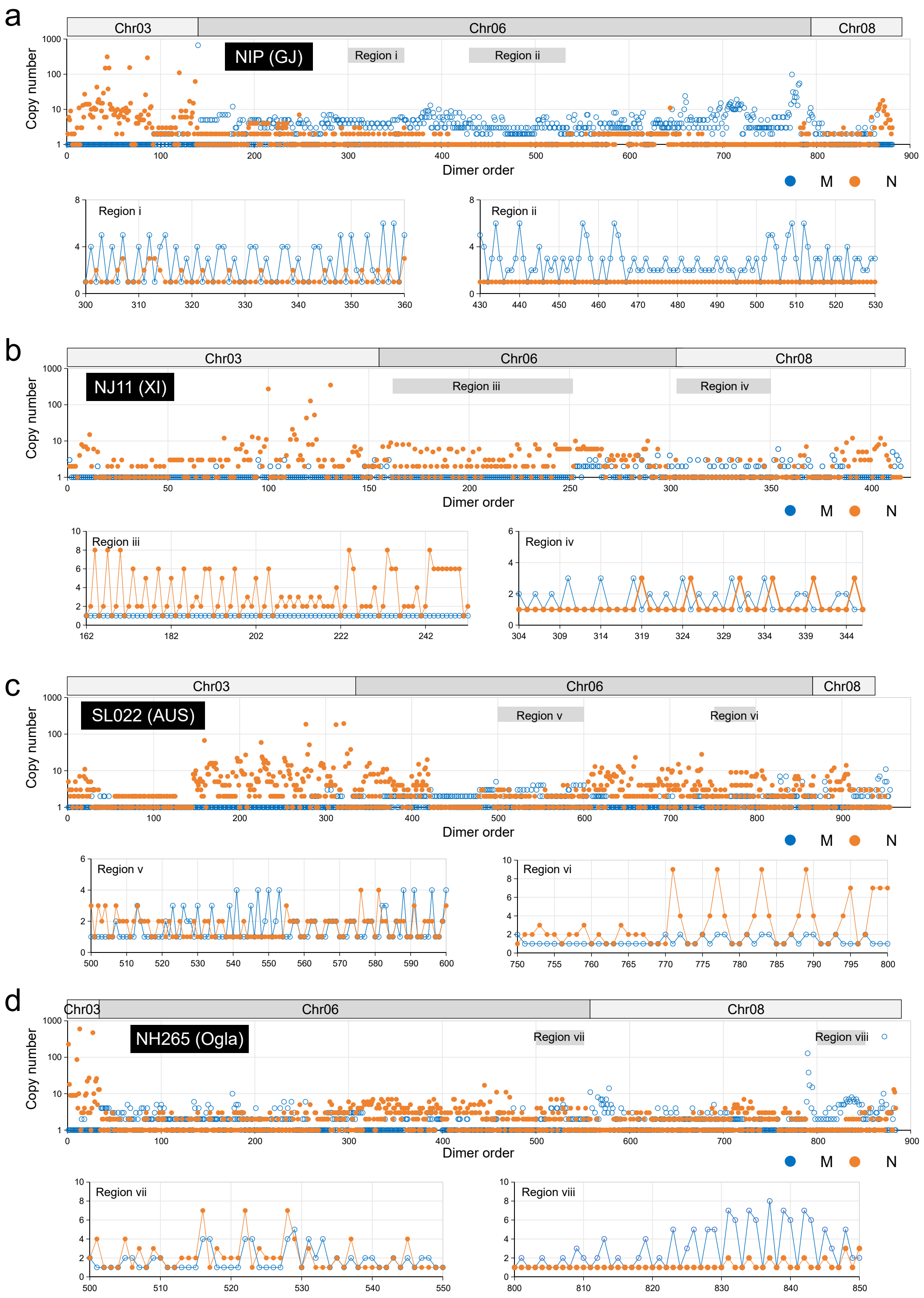


**Supplementary Fig. 15 |** Local homogenization of copy number bias within a dimer through tandem duplication. **a-d**, Repeat of bias pattern in monomer copy number of dimers in NIP(GJ), NJ11(XI), SL022(AUS) and NH265(Ogla).

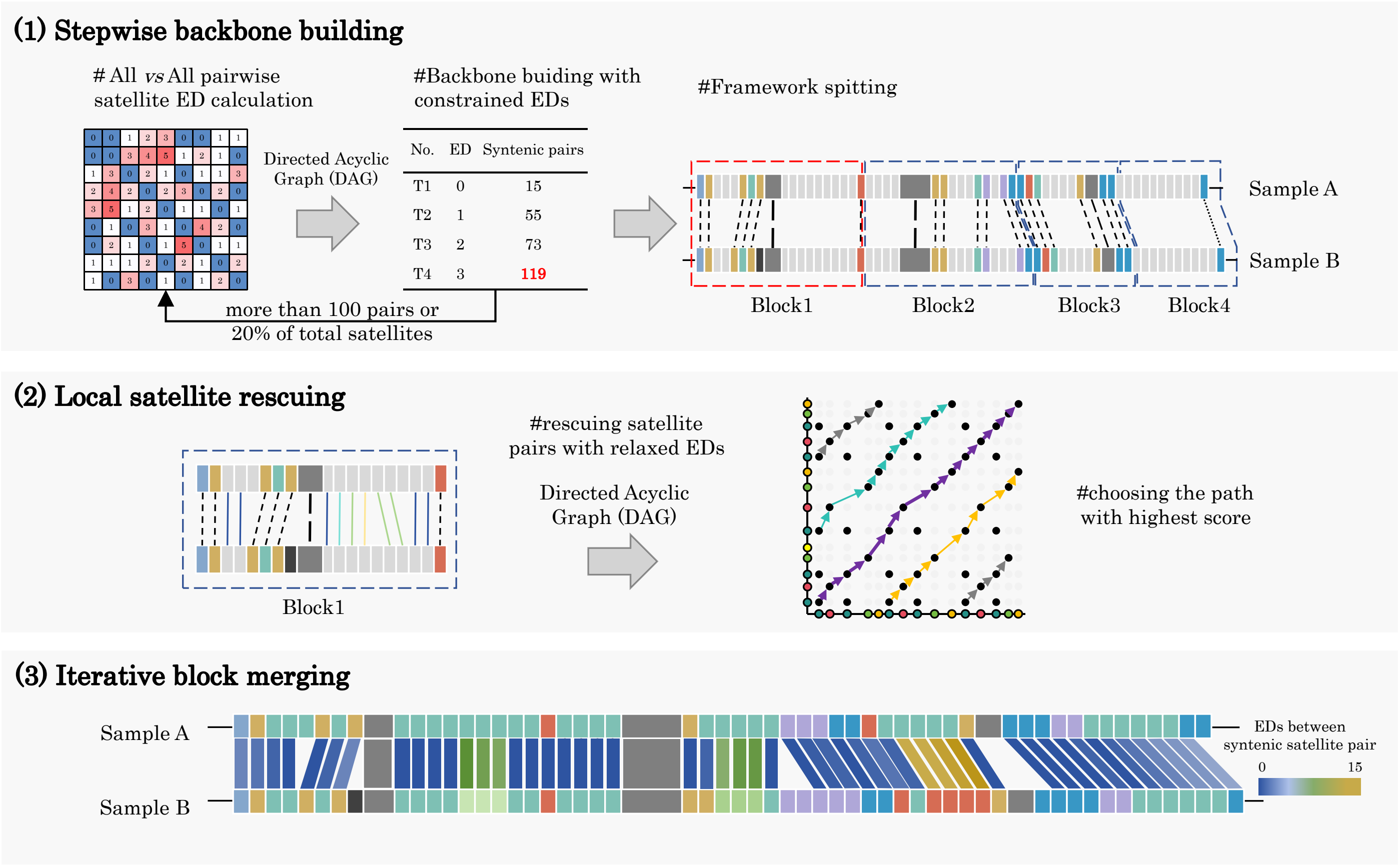


**Supplementary Fig. 16** | Workflow of SynPan-CEN. (1) Stepwise backbone building. All vs all pairwise satellite edit distance (ED) values were used to construct the Directed Acyclic Graph (DAG). To enhance accuracy in constructing synteny, the chain-backbone was built with the most conservative and highly-ordered monomer pairs by gradually increasing the ED value thresholds until the framework contained more than 20% of the total monomers or 100 monomers. This framework was then split by 10 monomer pairs within one window for downstream analysis. (2) Local satellite rescuing. DAGs were used again to rescue chains of syntenic satellite pairs within each window, employing relaxed EDs of 15. The sub-blocks with the highest scores were retained, and any non-overlapping monomer pairs were added to complete the matches. (3) Blocks from the chain-backbone were iteratively merged.

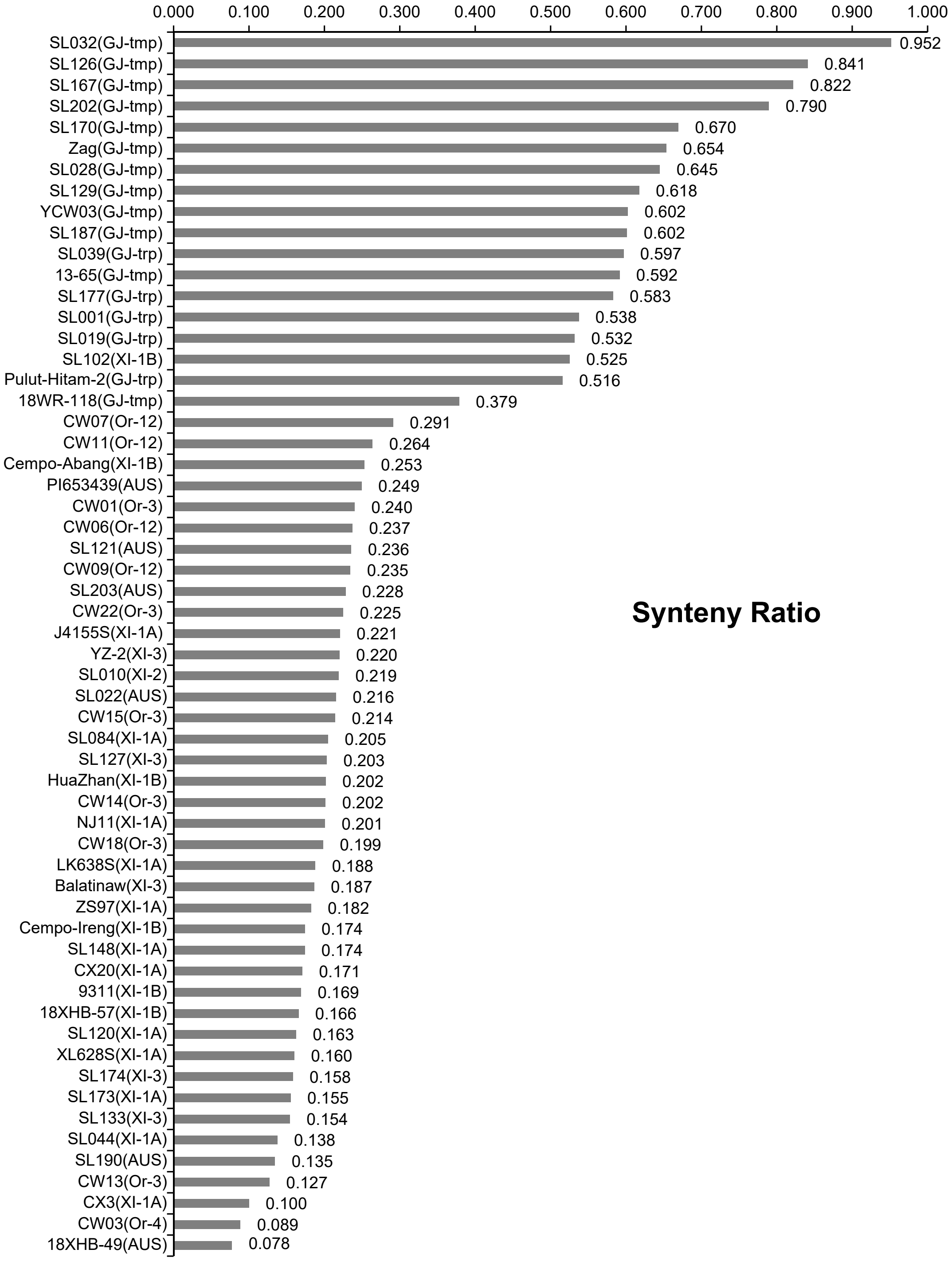


**Supplementary Fig. 17 |** Synteny ratio of *CEN155* satellites in rice centromeres against NIP (GJ-tmp) centromeres.

| Accession | Taxa | Chr01 | Chr02 | Chr03 | Chr04 | Chr05 | Chr06 | Chr07 | Chr08 | Chr09 | Chr10 | Chr11 | Chr12 |
|---|---|---|---|---|---|---|---|---|---|---|---|---|---|
| NH281 | Obar | 0.07 | 0.12 | 0.27 | 0.07 | 0.08 | 0.10 | 0.14 | 0.19 | 0.14 | 0.13 | 0.09 | 0.01 |
| NH284 | | 0.40 | 1.08 | 0.18 | 0.87 | 0.07 | 0.10 | 0.14 | 1.05 | 1.02 | 0.24 | 0.06 | 0.41 |
| NH285 | | 0.24 | 0.03 | 0.35 | 0.03 | 0.08 | 0.16 | 0.00 | 0.05 | 0.03 | 0.05 | 0.08 | 0.08 |
| NH286 | | 0.42 | 0.99 | 0.95 | 0.08 | 0.07 | 0.11 | 0.54 | 1.03 | 0.97 | 0.22 | 0.56 | 0.08 |
| NH265 | Ogla | 0.05 | 0.05 | 0.32 | 0.09 | 0.07 | 0.32 | 0.11 | 0.25 | 0.17 | 0.18 | 0.08 | 0.09 |
| NH267 | | 0.14 | 0.10 | 0.34 | 0.07 | 0.06 | 0.11 | 0.02 | 0.52 | 0.13 | 0.31 | 0.00 | 0.03 |
| NH271 | | 0.11 | 0.10 | 0.29 | 0.07 | 0.06 | 0.32 | 0.15 | 0.19 | 0.20 | 0.18 | 0.06 | 0.06 |
| NH273 | | 0.12 | 0.17 | 0.29 | 0.08 | 0.06 | 0.07 | 0.04 | 0.23 | 0.12 | 0.31 | 0.07 | 0.09 |
| CW06 | Or-12 | 0.42 | 0.26 | 0.49 | 0.26 | 0.12 | 0.08 | 0.00 | 0.51 | 0.00 | 0.21 | 0.05 | 0.35 |
| CW07 | | 0.42 | 0.38 | 0.86 | 0.47 | 0.37 | 0.15 | 0.07 | 0.69 | 0.08 | 0.35 | 0.01 | 0.21 |
| CW09 | | 0.15 | 0.19 | 0.83 | 0.24 | 0.74 | 0.08 | 0.00 | 0.53 | 0.05 | 0.18 | 0.01 | 0.26 |
| CW11 | | 0.18 | 0.20 | 0.82 | 0.21 | 0.74 | 0.07 | 0.00 | 0.54 | 0.17 | 0.24 | 0.08 | 0.27 |
| CW01 | Or-3 | 0.45 | 0.18 | 0.31 | 0.33 | 0.29 | 0.16 | 0.10 | 0.60 | 0.57 | 0.26 | 0.01 | 0.24 |
| CW13 | | 0.26 | 0.29 | 0.00 | 0.06 | 0.21 | 0.00 | 0.04 | 0.38 | 0.01 | 0.05 | 0.11 | 0.04 |
| CW14 | | 0.13 | 0.33 | 0.16 | 0.37 | 0.32 | 0.22 | 0.02 | 0.10 | 0.22 | 0.39 | 0.00 | 0.46 |
| CW15 | | 0.31 | 0.31 | 0.27 | 0.23 | 0.48 | 0.12 | 0.00 | 0.11 | 0.22 | 0.48 | 0.07 | 0.04 |
| CW18 | | 0.39 | 0.35 | 0.32 | 0.21 | 0.32 | 0.14 | 0.15 | 0.32 | 0.09 | 0.43 | 0.02 | 0.11 |
| CW22 | | 0.38 | 0.15 | 0.19 | 0.79 | 0.28 | 0.04 | 0.07 | 0.78 | 0.36 | 0.39 | 0.06 | 0.18 |
| CW03 | Or-4 | 0.04 | 0.20 | 0.16 | 0.03 | 0.09 | 0.06 | 0.01 | 0.07 | 0.00 | 0.11 | 0.06 | 0.20 |
| 13-65 | GJ-tmp | 0.81 | 0.67 | 0.21 | 0.93 | 0.79 | 0.39 | 0.85 | 0.39 | 0.61 | 0.28 | 0.63 | 0.99 |
| 18WR-118 | | 0.25 | 0.67 | 0.72 | 0.93 | 0.57 | 0.44 | 0.20 | 0.29 | 0.46 | 0.10 | 0.07 | 0.38 |
| SL028 | | 0.74 | 0.92 | 0.75 | 0.92 | 0.82 | 0.06 | 0.95 | 0.12 | 1.04 | 0.25 | 0.65 | 0.73 |
| SL032 | | 1.01 | 1.05 | 1.04 | 0.84 | 0.25 | 1.02 | 0.87 | 0.90 | 0.92 | 0.87 | 0.98 | 0.89 |
| SL126 | | 0.76 | 1.05 | 0.90 | 0.92 | 1.05 | 1.05 | 0.96 | 0.90 | 0.98 | 0.99 | 0.46 | 0.91 |
| SL129 | | 0.71 | 0.73 | 0.78 | 0.72 | 0.71 | 0.41 | 0.09 | 0.35 | 0.64 | 0.30 | 0.80 | 0.45 |
| SL167 | | 1.02 | 1.06 | 0.94 | 0.75 | 1.05 | 1.02 | 0.96 | 0.95 | 1.04 | 1.00 | 0.46 | 0.42 |
| SL170 | | 0.97 | 1.02 | 0.73 | 0.95 | 0.80 | 0.80 | 0.05 | 1.07 | 0.94 | 0.39 | 0.24 | 0.93 |
| SL187 | | 0.69 | 0.82 | 0.69 | 0.89 | 0.66 | 0.07 | 0.72 | 0.50 | 0.54 | 0.53 | 0.76 | 0.66 |
| SL202 | | 0.73 | 0.95 | 0.83 | 0.74 | 0.79 | 0.39 | 0.97 | 0.79 | 1.05 | 0.90 | 0.91 | 0.70 |
| YCW03 | | 0.77 | 0.75 | 0.72 | 0.80 | 0.71 | 0.38 | 0.59 | 0.35 | 0.63 | 0.30 | 0.73 | 0.42 |
| Zag | | 0.83 | 0.86 | 0.76 | 0.94 | 0.68 | 0.71 | 0.79 | 0.35 | 0.63 | 0.40 | 0.47 | 0.72 |
| Pulut-Hitam-2 | GJ-trp | 0.79 | 0.74 | 0.44 | 0.04 | 0.79 | 0.60 | 0.37 | 0.12 | 0.22 | 0.16 | 0.62 | 0.29 |
| SL001 | | 0.88 | 0.72 | 0.89 | 0.06 | 0.64 | 0.10 | 0.59 | 0.12 | 0.44 | 0.13 | 0.58 | 0.42 |
| SL019 | | 0.76 | 0.75 | 0.70 | 0.07 | 0.77 | 0.12 | 0.57 | 0.12 | 0.21 | 0.17 | 0.69 | 0.42 |
| SL039 | | 0.70 | 0.73 | 0.40 | 0.09 | 0.80 | 0.65 | 0.93 | 0.91 | 0.32 | 0.17 | 0.71 | 0.35 |
| SL177 | | 0.68 | 0.75 | 0.48 | 0.98 | 0.80 | 0.65 | 0.95 | 1.00 | 0.33 | 0.42 | 0.48 | 0.38 |
| 18XHB-49 | AUS | 0.06 | 0.01 | 0.21 | 0.03 | 0.04 | 0.02 | 0.07 | 0.17 | 0.01 | 0.17 | 0.06 | 0.13 |
| PI653439 | | 0.28 | 0.38 | 0.40 | 0.22 | 0.11 | 0.29 | 0.11 | 0.40 | 0.18 | 0.37 | 0.08 | 0.23 |
| SL022 | | 0.34 | 0.39 | 0.40 | 0.13 | 0.10 | 0.07 | 0.08 | 0.55 | 0.17 | 0.43 | 0.05 | 0.16 |
| SL121 | | 0.55 | 0.17 | 0.40 | 0.15 | 0.73 | 0.09 | 0.04 | 0.46 | 0.11 | 0.42 | 0.13 | 0.28 |
| SL190 | | 0.09 | 0.16 | 0.17 | 0.13 | 0.09 | 0.06 | 0.02 | 0.52 | 0.14 | 0.40 | 0.05 | 0.32 |
| SL203 | | 0.70 | 0.18 | 0.40 | 0.15 | 0.12 | 0.07 | 0.02 | 0.45 | 0.13 | 0.39 | 0.20 | 0.15 |
| CX20 | XI-1A | 0.20 | 0.18 | 0.54 | 0.20 | 0.11 | 0.06 | 0.02 | 0.58 | 0.13 | 0.14 | 0.10 | 0.17 |
| CX3 | | 0.18 | 0.19 | 0.21 | 0.04 | 0.01 | 0.00 | 0.02 | 0.11 | 0.00 | 0.01 | 0.02 | 0.08 |
| J4155S | | 0.20 | 0.17 | 0.49 | 0.82 | 0.83 | 0.10 | 0.02 | 0.53 | 0.13 | 0.42 | 0.12 | 0.32 |
| LK638S | | 0.18 | 0.18 | 0.49 | 0.17 | 0.81 | 0.09 | 0.02 | 0.38 | 0.26 | 0.14 | 0.08 | 0.13 |
| NJ11 | | 0.17 | 0.15 | 0.53 | 0.20 | 0.80 | 0.06 | 0.02 | 0.66 | 0.14 | 0.19 | 0.13 | 0.24 |
| SL044 | | 0.27 | 0.19 | 0.27 | 0.06 | 0.80 | 0.02 | 0.01 | 0.15 | 0.00 | 0.03 | 0.03 | 0.10 |
| SL084 | | 0.13 | 0.37 | 0.45 | 0.18 | 0.81 | 0.08 | 0.03 | 0.53 | 0.13 | 0.19 | 0.06 | 0.32 |
| SL120 | | 0.31 | 0.14 | 0.42 | 0.23 | 0.12 | 0.10 | 0.02 | 0.54 | 0.11 | 0.17 | 0.03 | 0.29 |
| SL148 | | 0.23 | 0.18 | 0.45 | 0.22 | 0.80 | 0.06 | 0.02 | 0.50 | 0.13 | 0.19 | 0.06 | 0.13 |
| SL173 | | 0.21 | 0.19 | 0.19 | 0.19 | 0.72 | 0.06 | 0.03 | 0.54 | 0.13 | 0.20 | 0.08 | 0.13 |
| XL628S | | 0.16 | 0.15 | 0.15 | 0.27 | 0.80 | 0.09 | 0.02 | 0.53 | 0.12 | 0.54 | 0.09 | 0.13 |
| ZS97 | | 0.36 | 0.25 | 0.44 | 0.22 | 0.77 | 0.06 | 0.02 | 0.54 | 0.06 | 0.19 | 0.04 | 0.13 |
| 9311 | XI-1B | 0.12 | 0.15 | 0.56 | 0.23 | 0.80 | 0.09 | 0.02 | 0.46 | 0.12 | 0.17 | 0.14 | 0.02 |
| 18XHB-57 | | 0.21 | 0.17 | 0.28 | 0.14 | 0.13 | 0.05 | 0.04 | 0.43 | 0.11 | 0.31 | 0.12 | 0.32 |
| Cempo-Abang | | 0.34 | 0.17 | 0.81 | 0.18 | 0.77 | 0.10 | 0.02 | 0.53 | 0.11 | 0.32 | 0.17 | 0.02 |
| Cempo-Ireng | | 0.23 | 0.15 | 0.30 | 0.12 | 0.85 | 0.12 | 0.02 | 0.39 | 0.10 | 0.20 | 0.18 | 0.03 |
| HuaZhan | | 0.18 | 0.18 | 0.56 | 0.21 | 0.77 | 0.11 | 0.03 | 0.55 | 0.12 | 0.31 | 0.08 | 0.31 |
| SL102 | | 0.70 | 0.15 | 0.94 | 0.93 | 0.71 | 1.02 | 0.03 | 0.55 | 0.50 | 1.00 | 0.00 | 0.44 |
| SL010 | XI-2 | 0.71 | 0.40 | 0.18 | 0.15 | 0.11 | 0.08 | 0.11 | 0.52 | 0.06 | 0.31 | 0.13 | 0.11 |
| Balatinaw | XI-3 | 0.24 | 0.17 | 0.44 | 0.11 | 0.76 | 0.15 | 0.04 | 0.51 | 0.12 | 0.18 | 0.08 | 0.17 |
| SL127 | | 0.30 | 0.16 | 0.46 | 0.14 | 0.79 | 0.07 | 0.02 | 0.85 | 0.12 | 0.20 | 0.09 | 0.32 |
| SL133 | | 0.21 | 0.13 | 0.23 | 0.23 | 0.80 | 0.09 | 0.02 | 0.42 | 0.11 | 0.20 | 0.00 | 0.29 |
| SL174 | | 0.31 | 0.15 | 0.37 | 0.23 | 0.15 | 0.07 | 0.02 | 0.49 | 0.10 | 0.30 | 0.07 | 0.10 |
| YZ-2 | | 0.35 | 0.22 | 0.48 | 0.27 | 0.79 | 0.10 | 0.03 | 0.54 | 0.13 | 0.19 | 0.08 | 0.29 |

**Supplementary Fig. 18 |** Heat map showing the synteny ratio between NIP (GJ-tmp) and other rice centromeres across 12 chromosomes, with Chr05 centromeres showing a significantly higher syntenic relationship between XI and GJ.

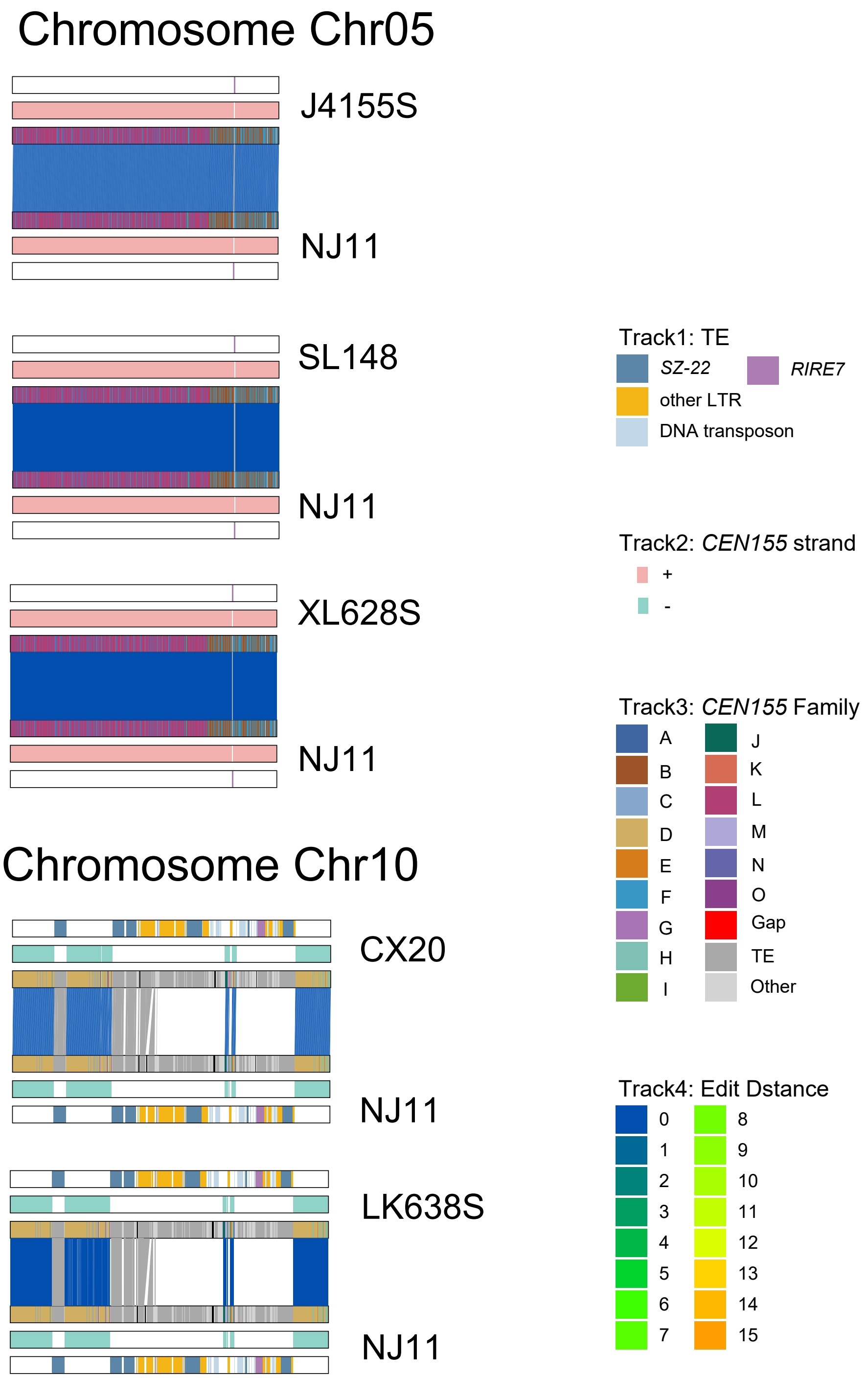


**Supplementary Fig. 19 |** Satellite syntenic relationships between centromeres of J4155S, SL148, and XL628S with NJ11 on Chr05, and CX20, LK638S on Chr10. Tracks show centromere annotation, including TE families, *CEN155* strands, *CEN155* superfamilies, and edit distances, respectively.

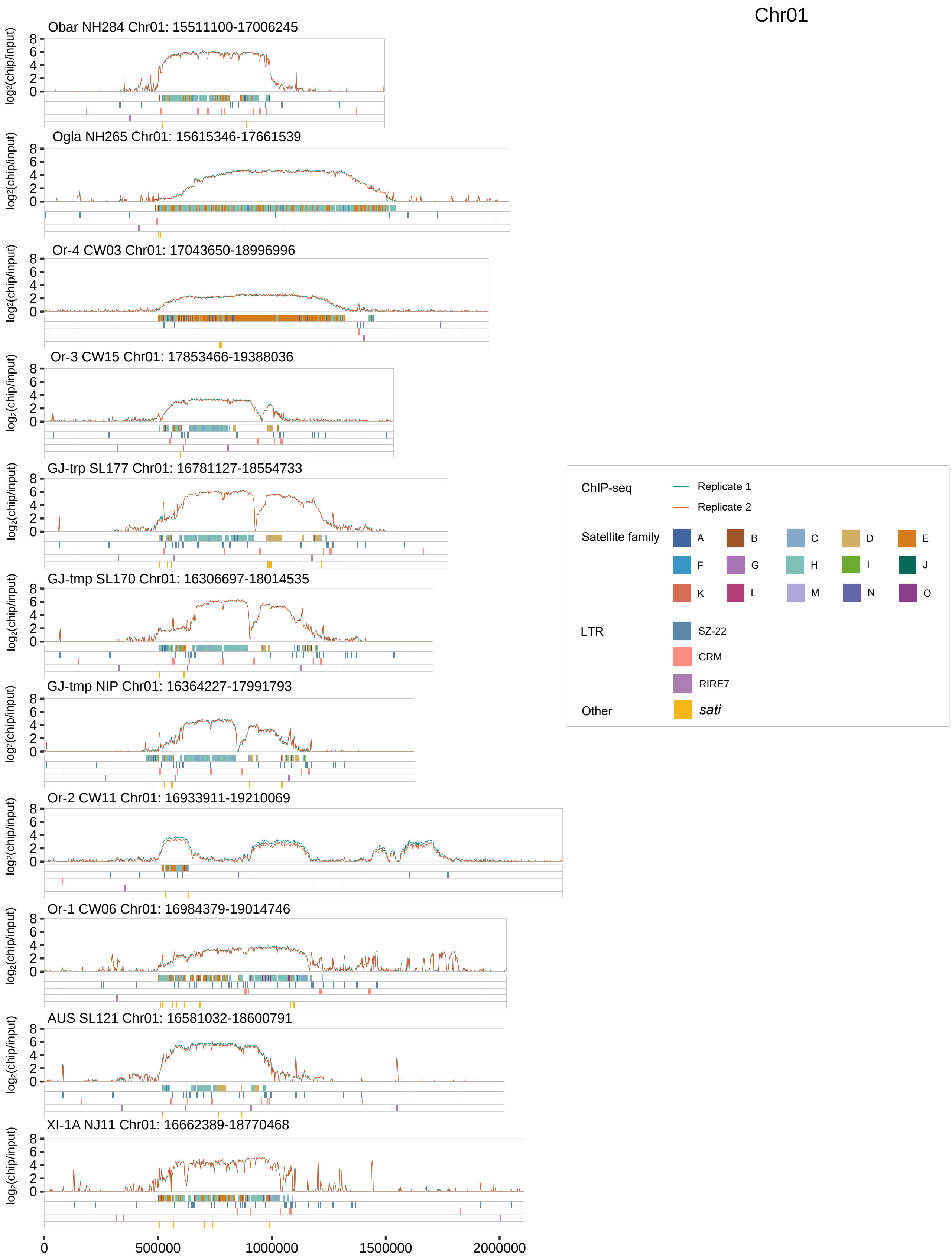


**Supplementary Fig. 20** | Genomic features of the functional centromere and its flanking regions on chromosome Chr01. Top, CENH3 ChIP-seq enrichment ($\log_2$(ChIP/input), two replicates) in 10-Kb windows. Track 1, satellite superfamilies; Track 2, LTRs *SZ-22*; Track 3, LTRs *CRM*; Track 4, LTRs *RIRE7*; Track 5, *sati* (non-canonical satellites).

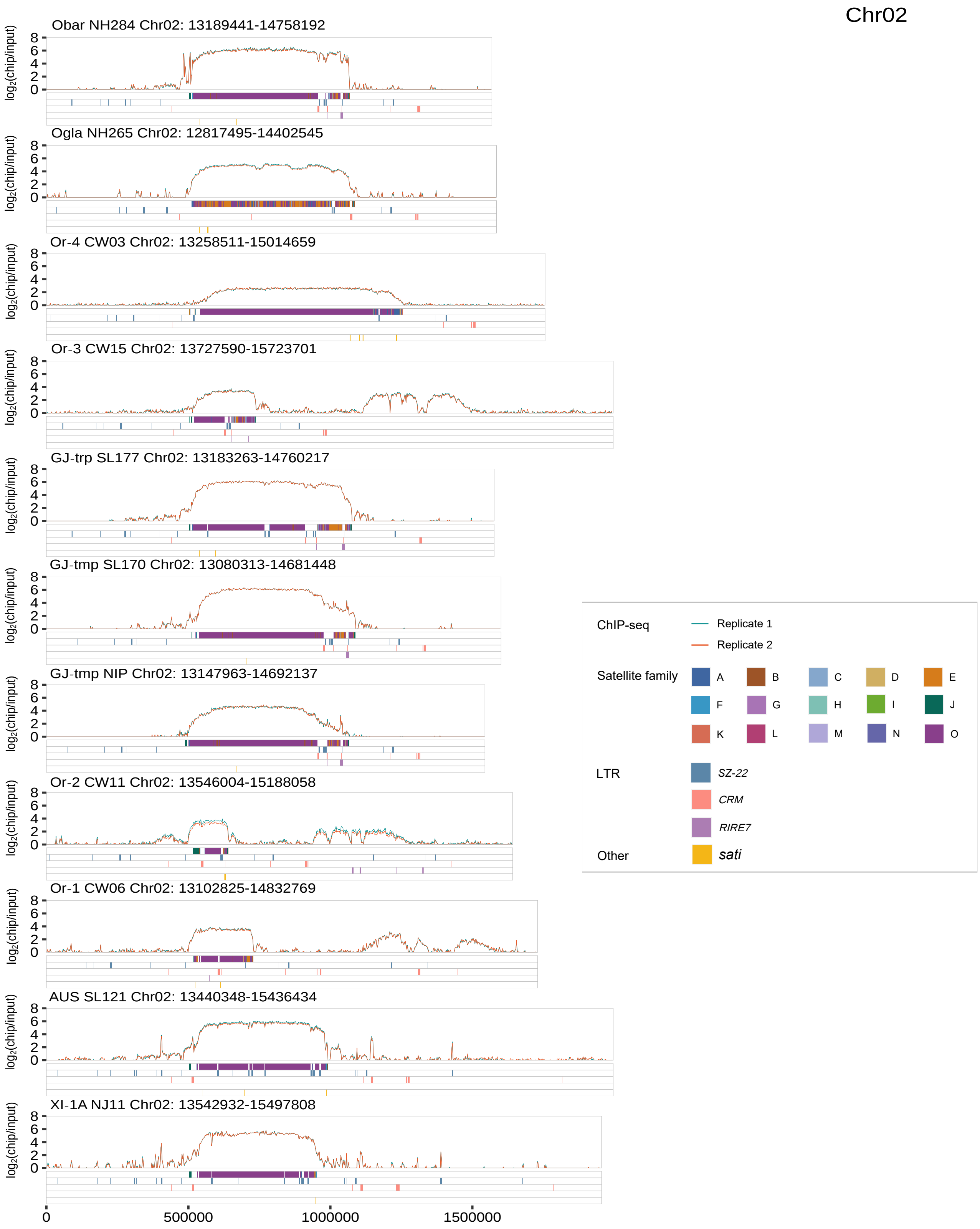


**Supplementary Fig. 21** | Genomic features of the functional centromere and its flanking regions on chromosome Chr02. Top, CENH3 ChIP-seq enrichment ($\log_2$(ChIP/input), two replicates) in 10-Kb windows. Track 1, satellite superfamilies; Track 2, LTRs *SZ-22*; Track 3, LTRs *CRM;* Track 4, LTRs *RIRE7*; Track 5, *sati* (non-canonical satellites).

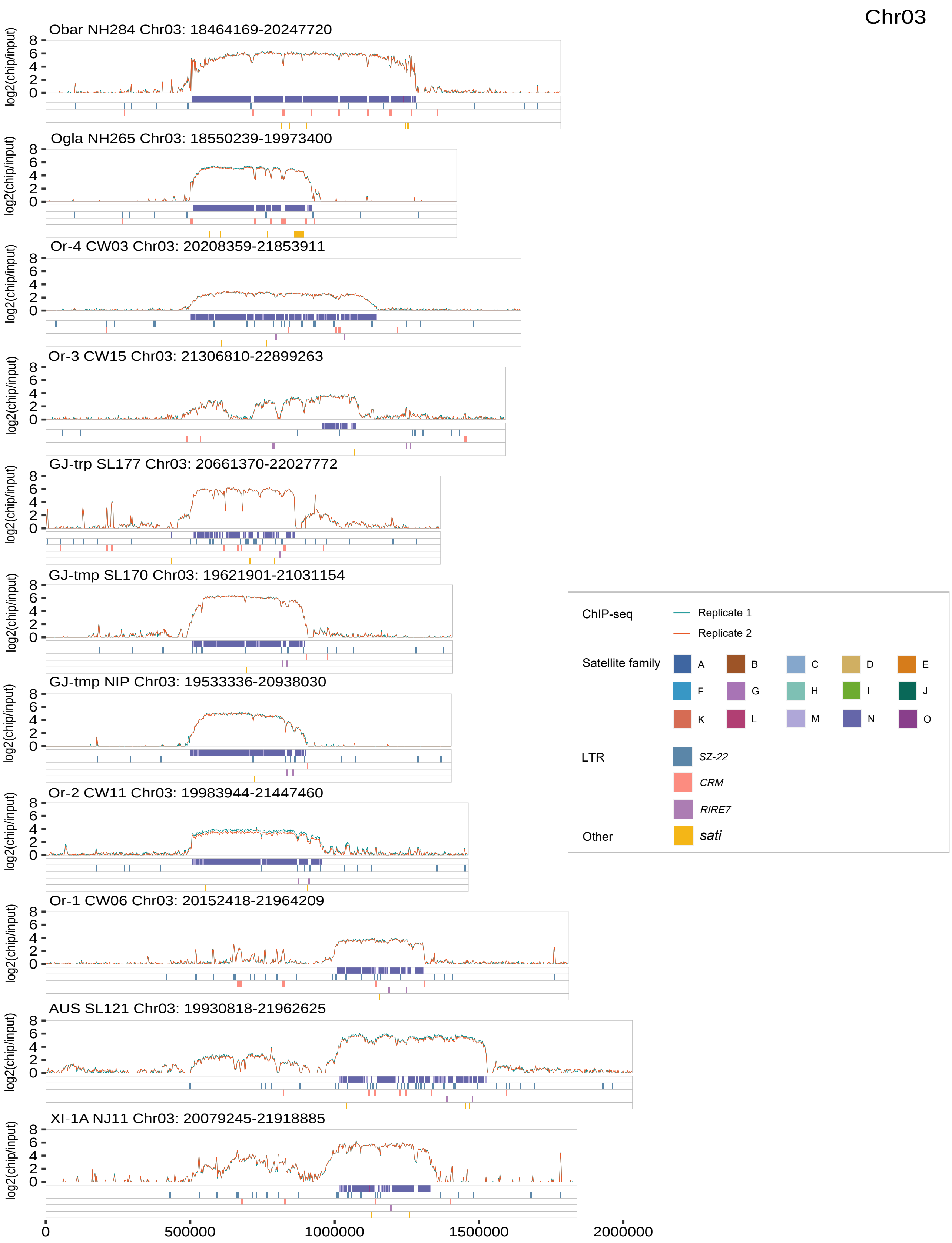


**Supplementary Fig. 22** | Genomic features of the functional centromere and its flanking regions on chromosome Chr03. Top, CENH3 ChIP-seq enrichment ($\log_2$(ChIP/input), two replicates) in 10-Kb windows. Track 1, satellite superfamilies; Track 2, LTRs *SZ-22*; Track 3, LTRs *CRM*; Track 4, LTRs *RIRE7*; Track 5, *sati* (non-canonical satellites).

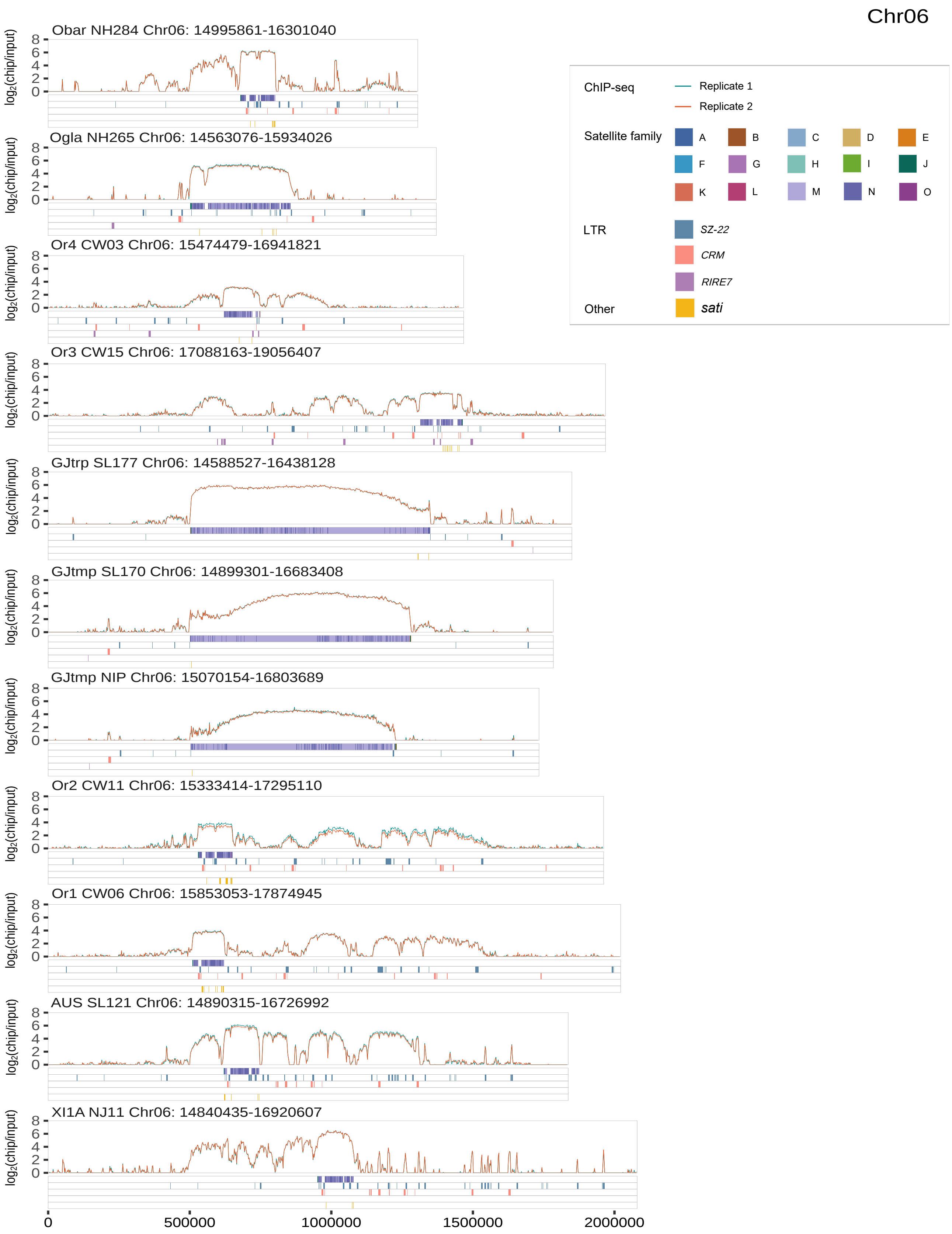


**Supplementary Fig. 23** | Genomic features of the functional centromere and its flanking regions on chromosome Chr06. Top, CENH3 ChIP-seq enrichment ($log_2$(ChIP/input), two replicates) in 10-Kb windows. Track 1, satellite superfamilies; Track 2, LTRs *SZ-22*; Track 3, LTRs *CRM*; Track 4, LTRs *RIRE7*; Track 5, *sati* (non-canonical satellites).

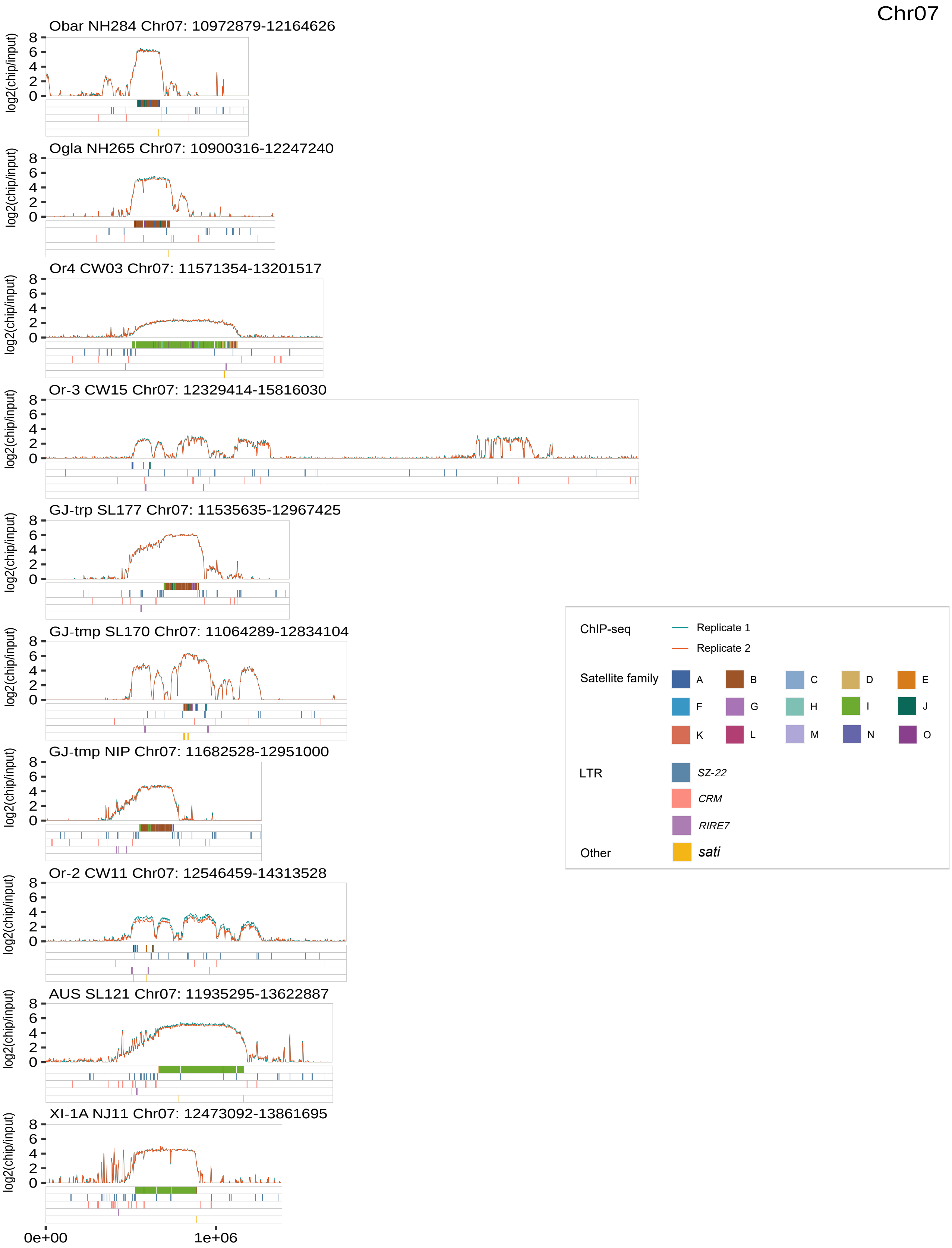


**Supplementary Fig. 24** | Genomic features of the functional centromere and its flanking regions on chromosome Chr07. Top, CENH3 ChIP-seq enrichment (log$_2$(ChIP/input), two replicates) in 10-Kb windows. Track 1, satellite superfamilies; Track 2, LTRs *SZ-22*; Track 3, LTRs *CRM*; Track 4, LTRs *RIRE7*; Track 5, short intervals (non-canonical satellites).

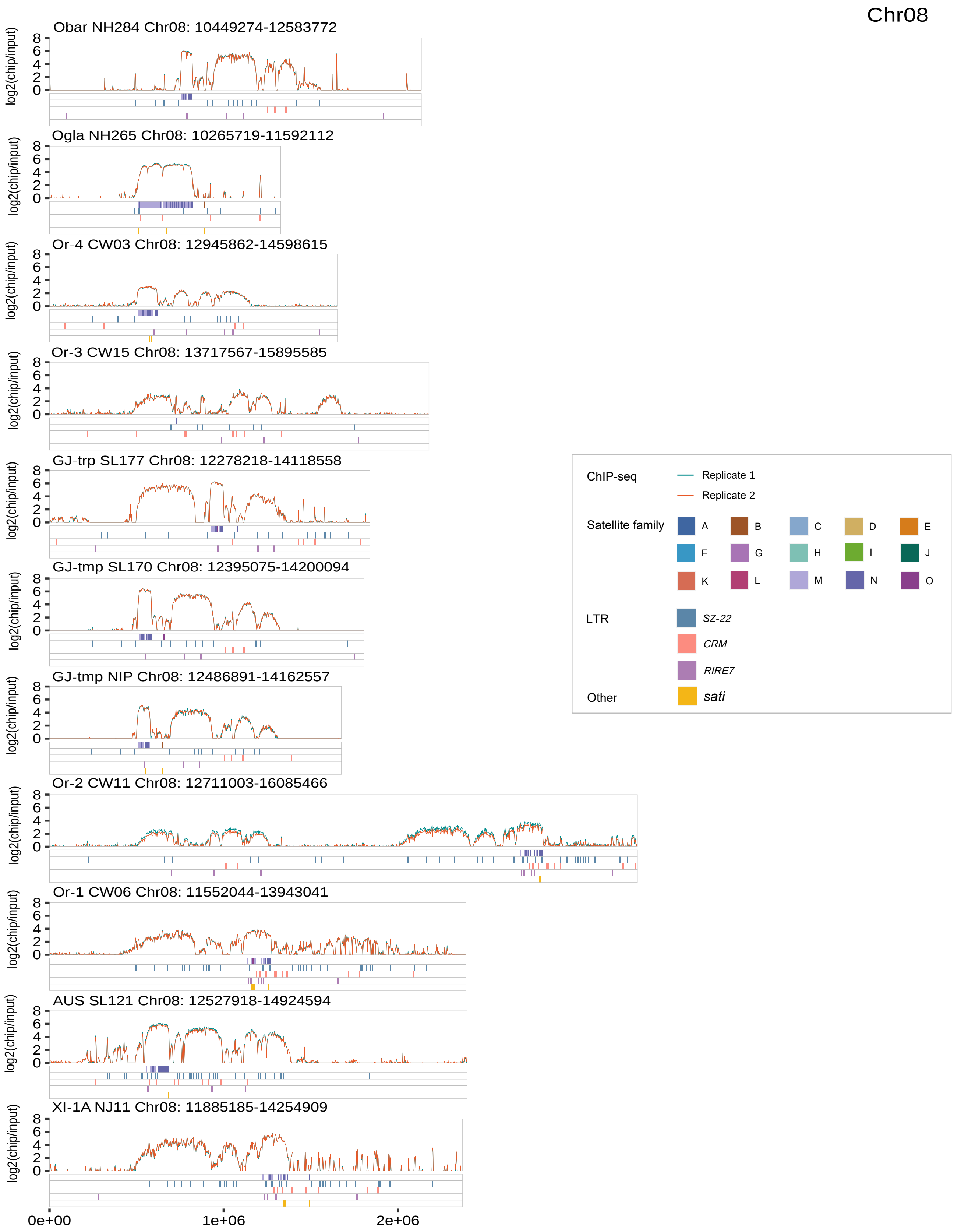


**Supplementary Fig. 25 |** Genomic features of the functional centromere and its flanking regions on chromosome Chr08. Top, CENH3 ChIP-seq enrichment ($log_2$(ChIP/input), two replicates) in 10-Kb windows. Track 1, satellite superfamilies; Track 2, LTRs *SZ-22*; Track 3, LTRs *CRM*; Track 4, LTRs *RIRE7*; Track 5, *sati* (non-canonical satellites).

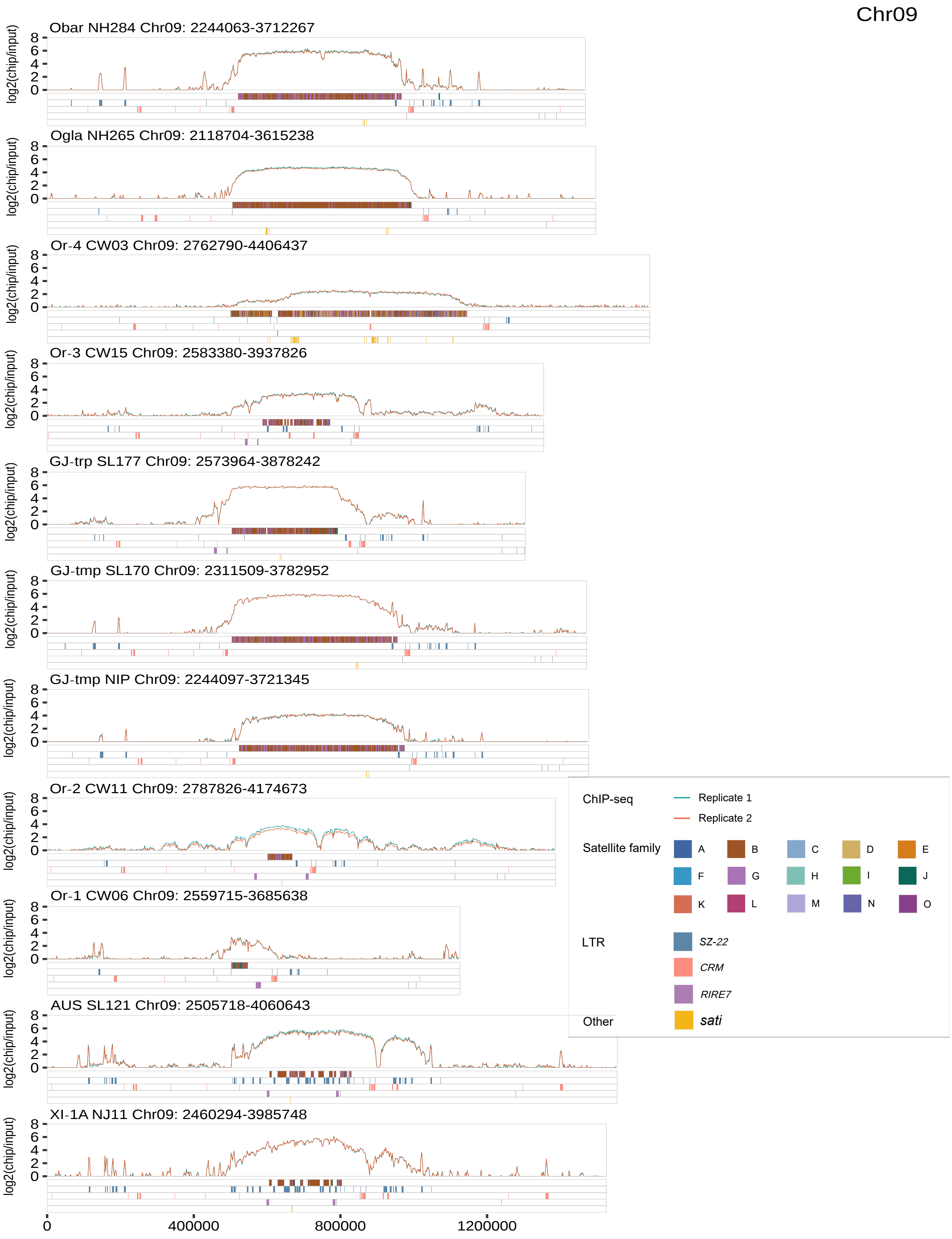


**Supplementary Fig. 26 |** Genomic features of the functional centromere and its flanking regions on chromosome Chr09. Top, CENH3 ChIP-seq enrichment (log$_2$(ChIP/input), two replicates) in 10-Kb windows. Track 1, satellite superfamilies; Track 2, LTRs *SZ-22*; Track 3, LTRs *CRM*; Track 4, LTRs *RIRE7*; Track 5, *sati* (non-canonical satellites).

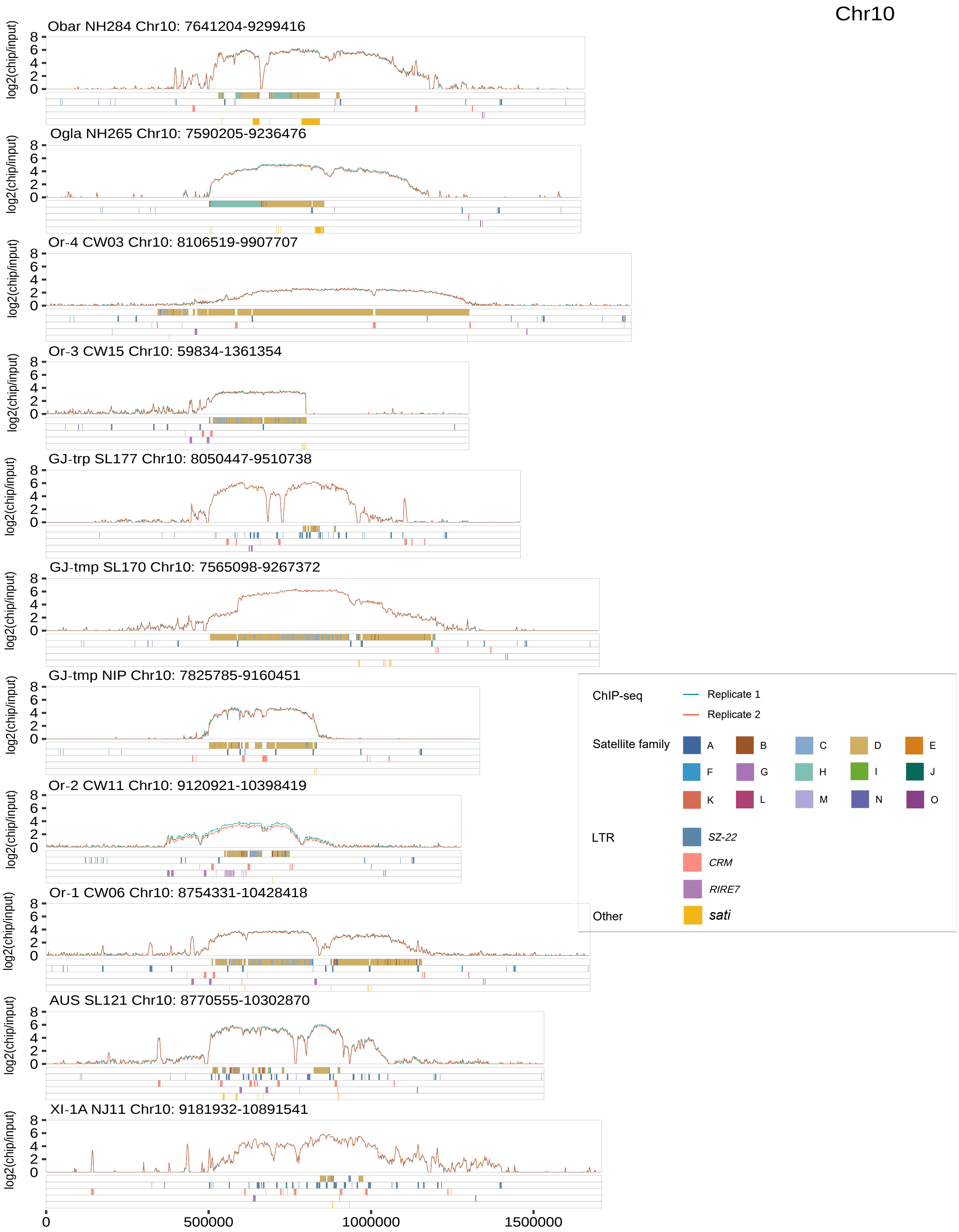


**Supplementary Fig. 27** | Genomic features of the functional centromere and its flanking regions on chromosome Chr10. Top, CENH3 ChIP-seq enrichment ($\log_2$(ChIP/input), two replicates) in 10-Kb windows. Track 1, satellite superfamilies; Track 2, LTRs *SZ-22*; Track 3, LTRs *CRM*; Track 4, LTRs *RIRE7*; Track 5, *sati* (non-canonical satellites).

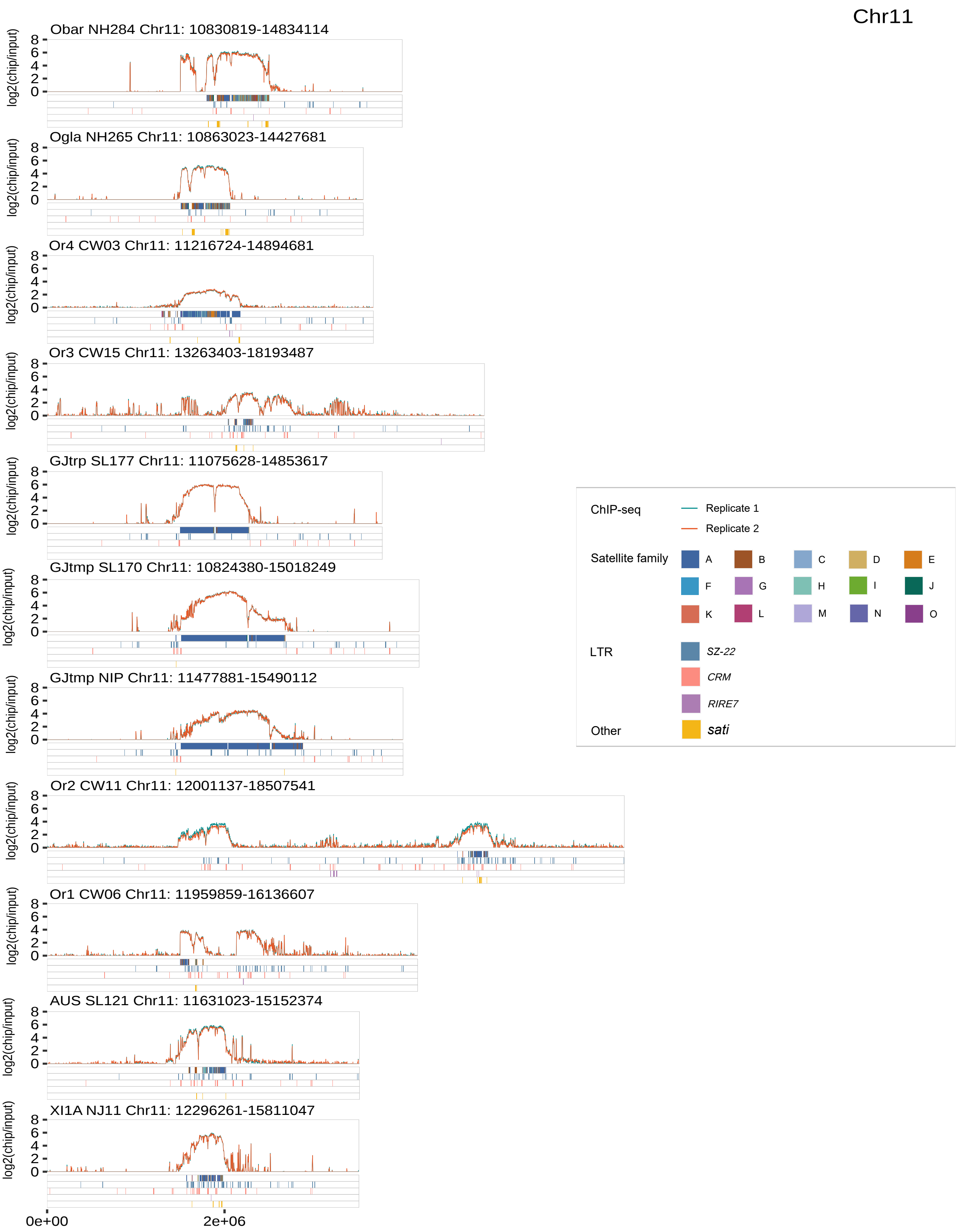


**Supplementary Fig. 28 |** Genomic features of the functional centromere and its flanking regions on chromosome Chr11. Top, CENH3 ChIP-seq enrichment ($log_2$(ChIP/input), two replicates) in 10-Kb windows. Track 1, satellite superfamilies; Track 2, LTRs *SZ-22*; Track 3, LTRs *CRM*; Track 4, LTRs *RIRE7*; Track 5, *sati* (non-canonical satellites).

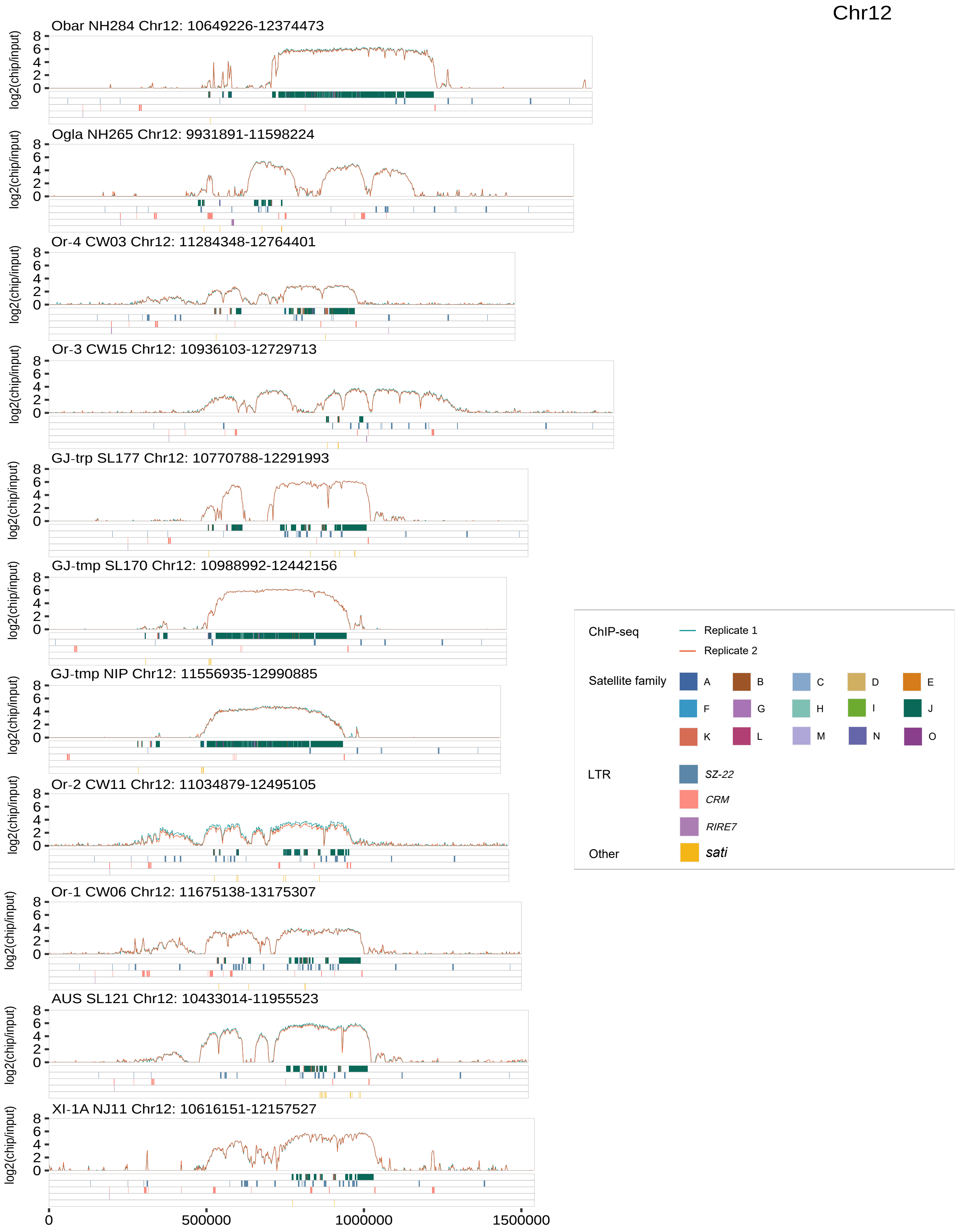


**Supplementary Fig. 29 |** Genomic features of the functional centromere and its flanking regions on chromosome Chr12. Top, CENH3 ChIP-seq enrichment (log$_2$(ChIP/input), two replicates) in 10-Kb windows. Track 1, satellite superfamilies; Track 2, LTRs *SZ-22*; Track 3, LTRs *CRM*; Track 4, LTRs *RIRE7*; Track 5, *sati* (non-canonical satellites).

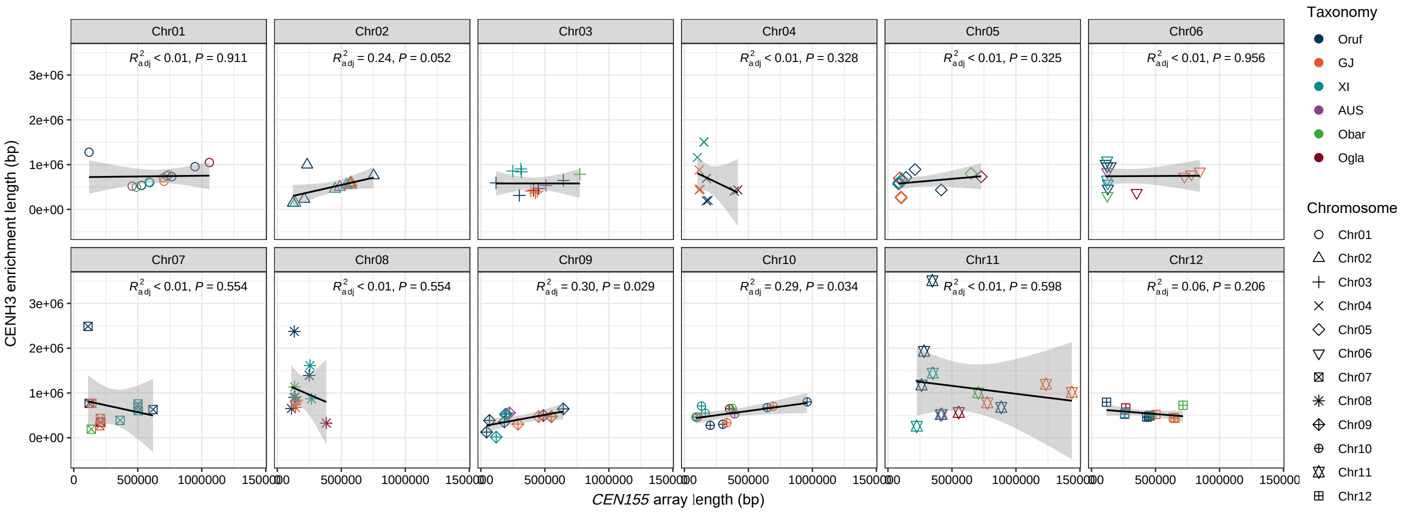


**Supplementary Fig. 30 |** Correlation between the length of CENH3 enriched regions and *CEN155*-defined centromere size for each chromosome.

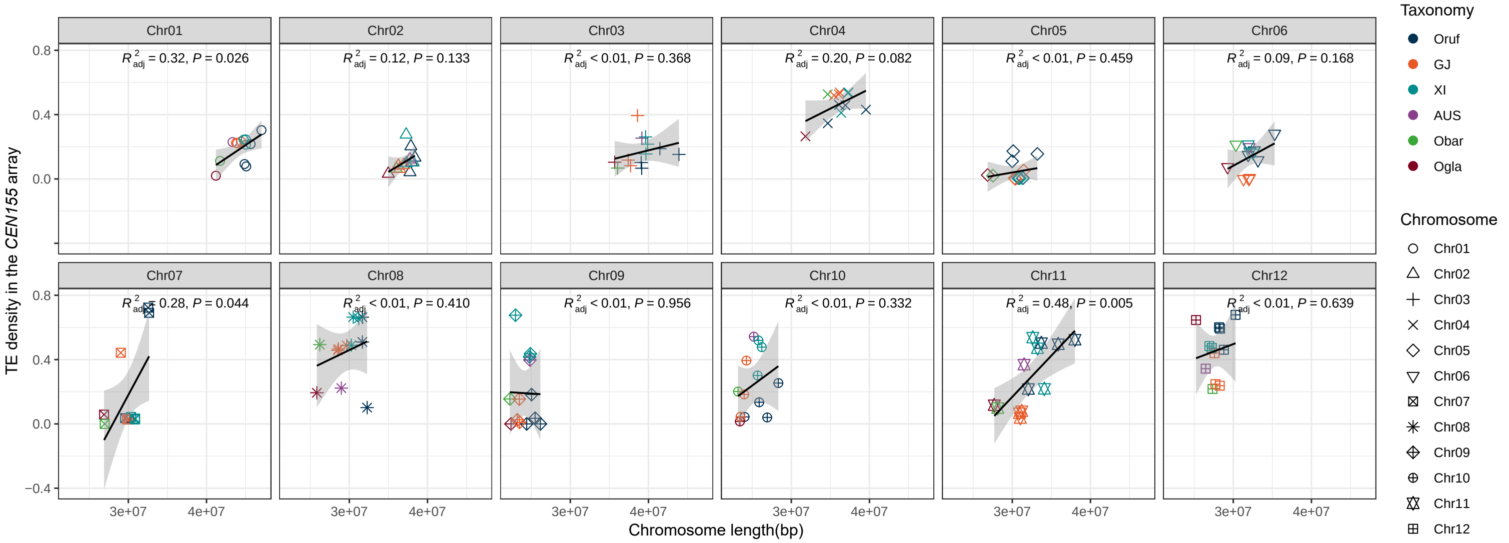


**Supplementary Fig. 31** | Correlation between chromosome length and TE density across the 12 chromosomes.

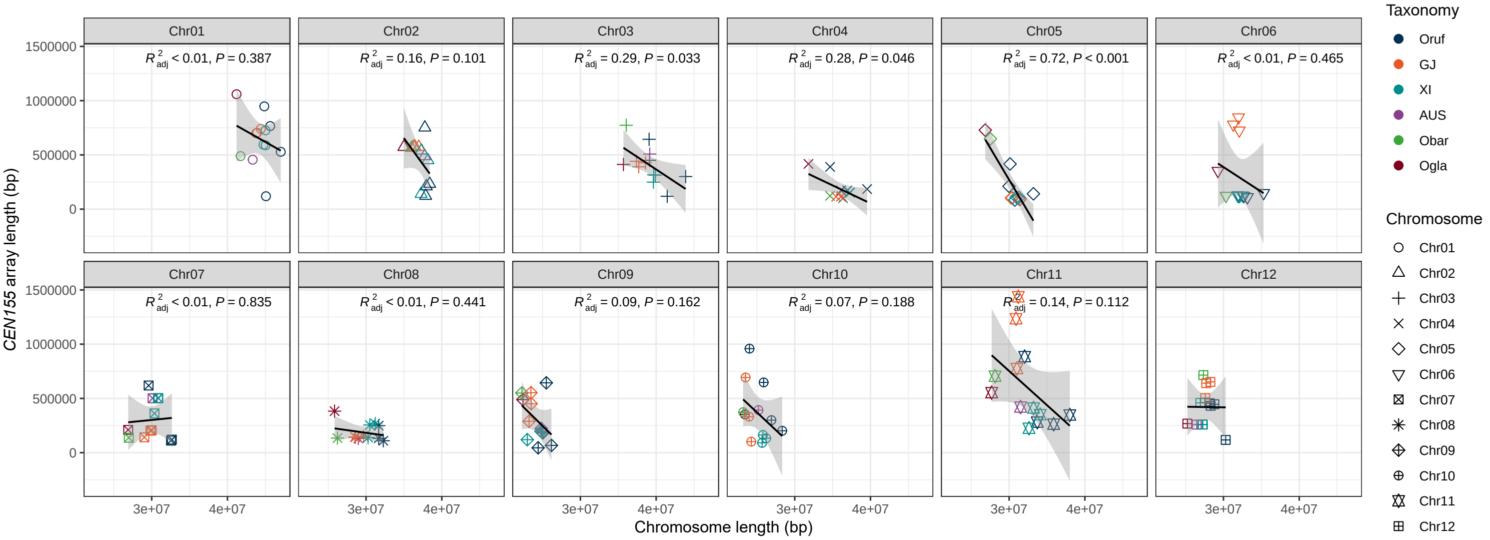


**Supplementary Fig. 32 |** Correlation between the chromosome length and *CEN155* array length across the 12 chromosomes.

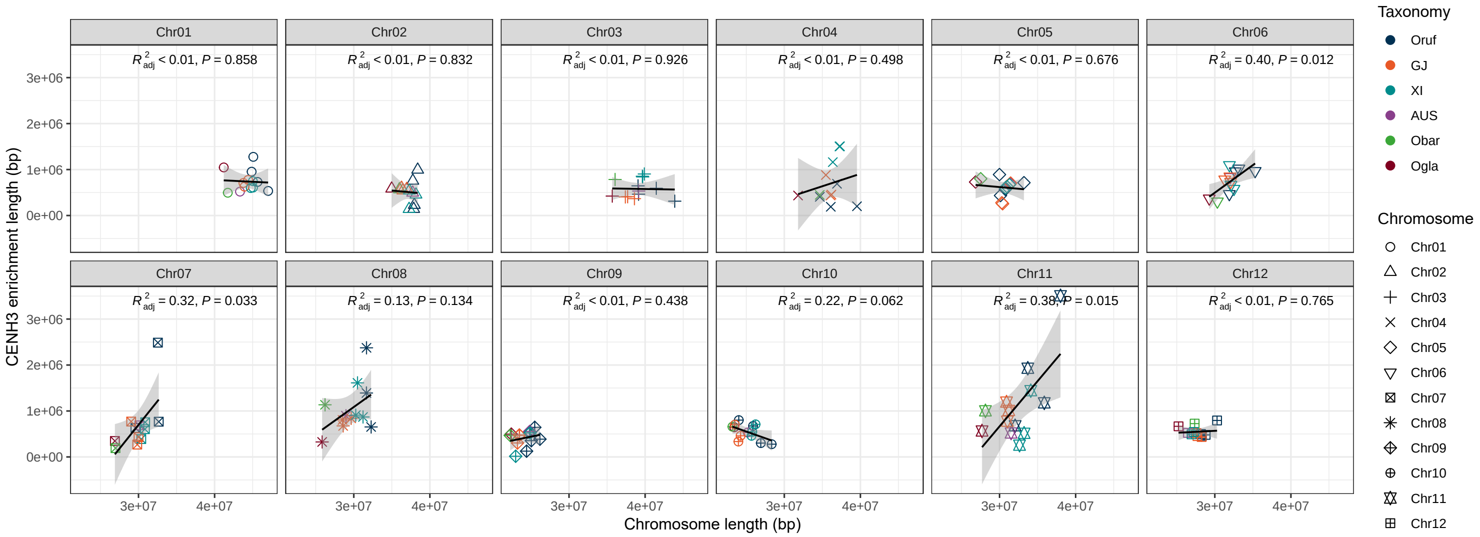


**Supplementary Fig. 33 |** Correlation between chromosome length and CENH3 enrichment length across the 12 chromosomes.

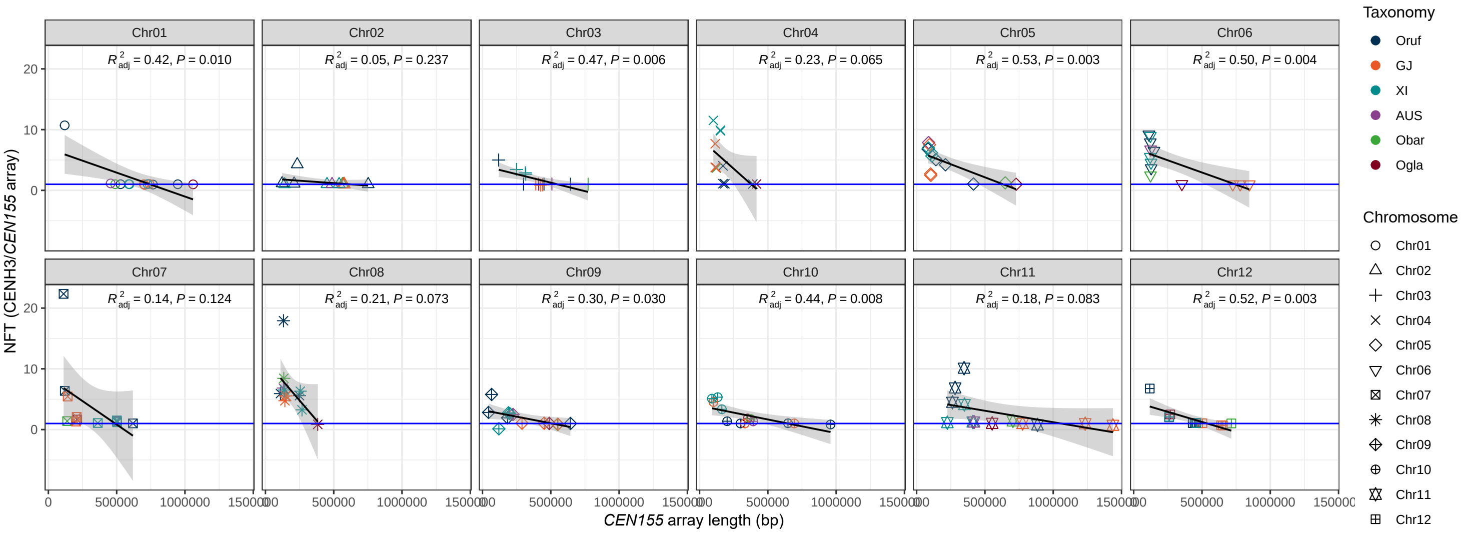


**Supplementary Fig. 34 |** Correlation between *CEN155* array length and neocentromere formation tendency (NFT) across 12 chromosomes.

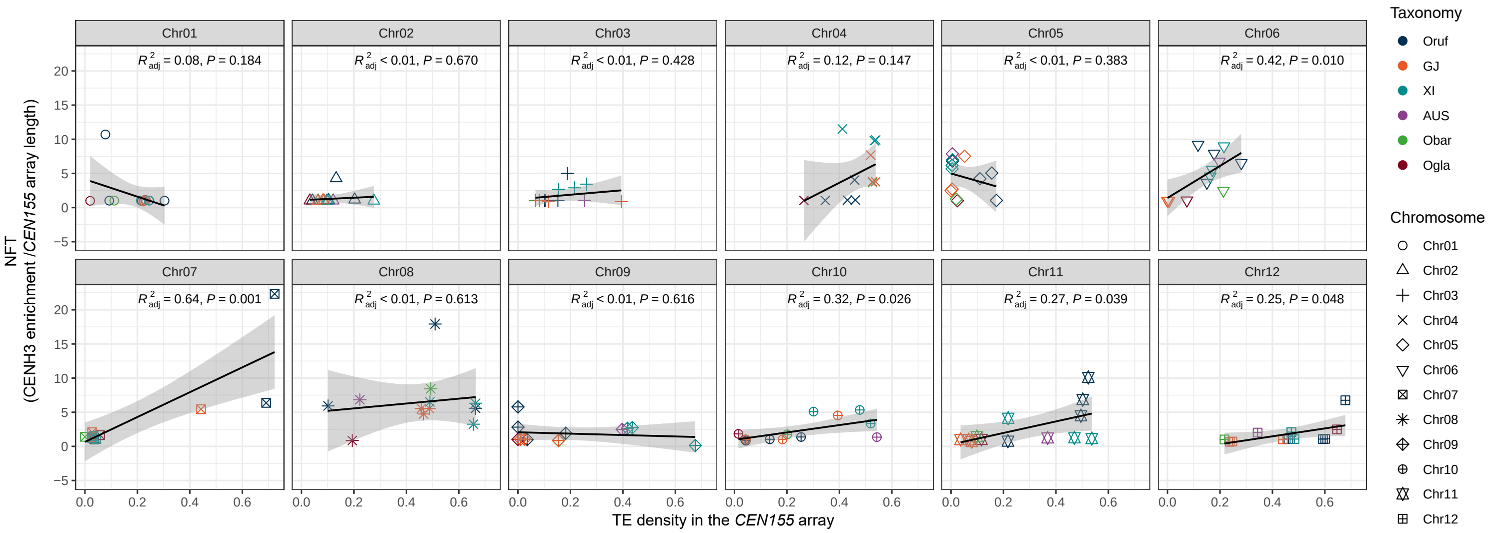


**Supplementary Fig. 35 |** Correlation between TE density in the *CEN155* array and neocentromere formation tendency (NFT) across 12 chromosomes.

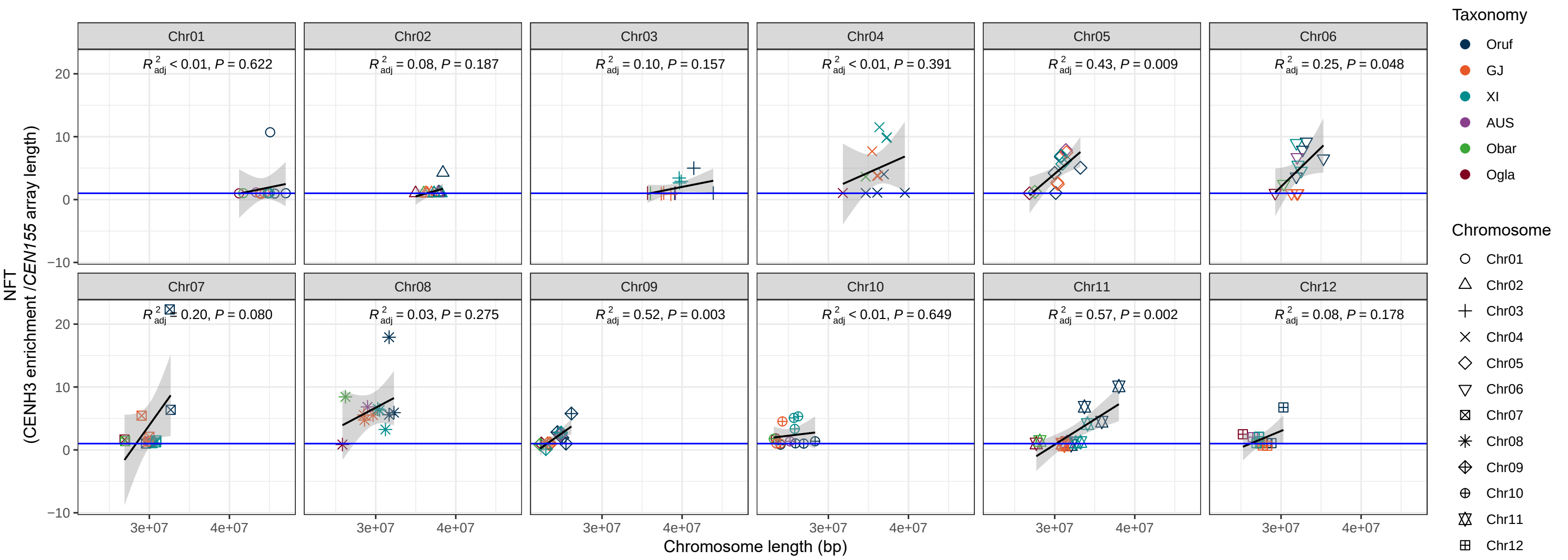


**Supplementary Fig. 36 |** Correlation between chromosome length and neocentromere formation tendency (NFT) across 12 chromosomes.

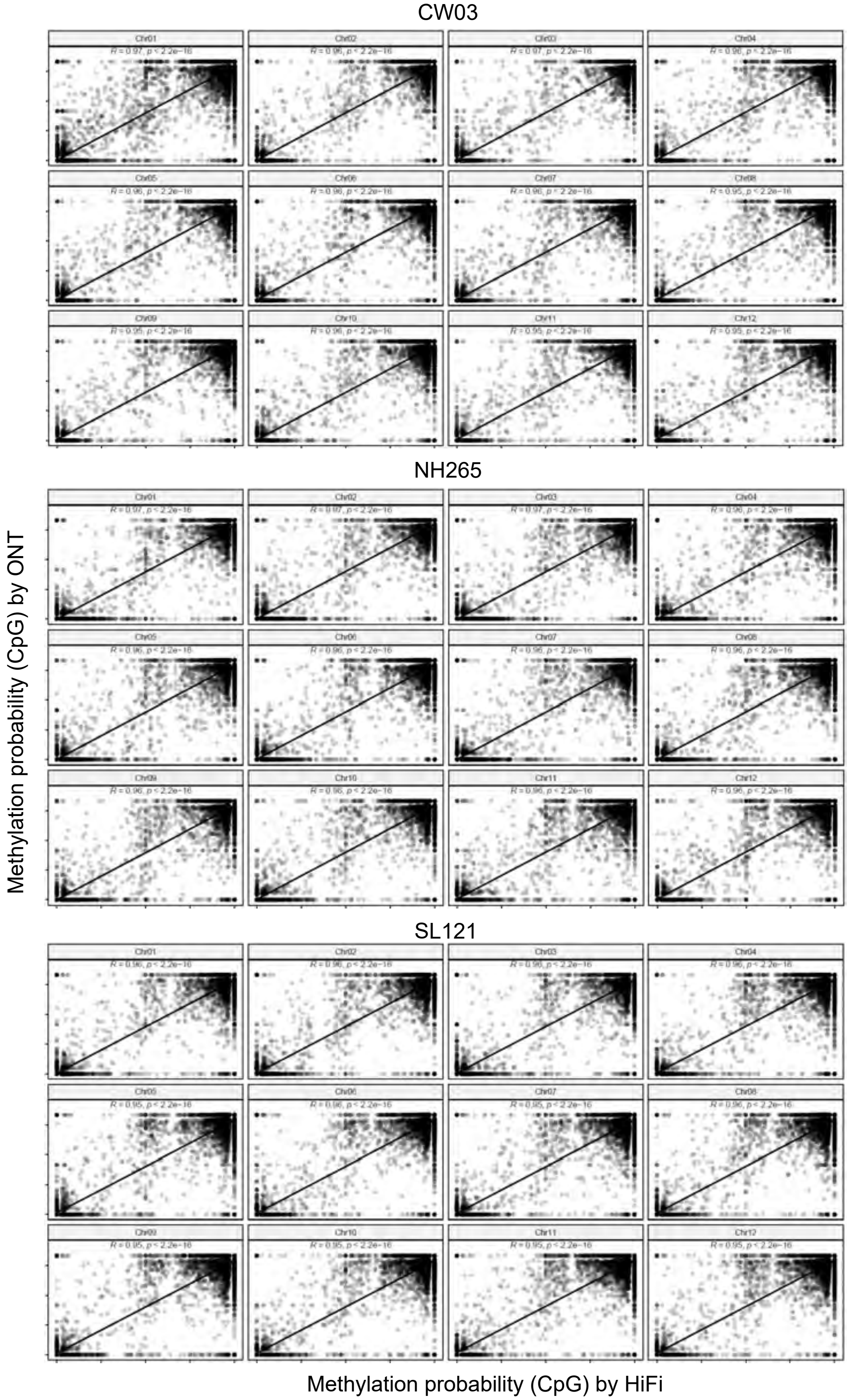


**Supplementary Fig. 37 |** Comparison of profiled DNA methylation probability from ONT sequencing data and HiFi data. Each point represents a randomly selected base C.
